# LanD-like Flavoprotein-Catalyzed Aminovinyl-Cysteine Formation through Oxidative Decarboxylation and Cyclization of a Peptide at the C-Terminus

**DOI:** 10.1101/2020.02.13.947028

**Authors:** Jingyu Liu, Yanping Qiu, Tao Fu, Miao Li, Yuqing Li, Qian Yang, Zhijun Tang, Haoyu Tang, Guangyu Li, Lifeng Pan, Wen Liu

## Abstract

Aminovinyl-cysteine residues arise from processing the C-terminal l-Cys and an internal l-Ser/l-Thr or l-Cys of a peptide. Formation of these nonproteinogenic amino acids, which occur in a macrocyclic ring of diverse ribosomally synthesized lanthipeptides and non-lanthipeptides, remains poorly understood. Here, we report that LanD-like flavoproteins in the biosynthesis of distinct non-lanthipeptides share an unexpected dual activity for aminovinyl-cysteine formation. Each flavoprotein catalyzes oxidative decarboxylation of the C-terminal l-Cys and couples the resulting enethiol nucleophile with the internal residue to afford a thioether linkage for peptide cyclization. The cyclization step, which largely depends on proximity effect by positioning the enethiol intermediate with a bent conformation at the active site, can be substrate-dependent, proceeding inefficiently through nucleophilic substitution for an unmodified peptide or efficiently through Michael addition for a dehydrated/dethiolated peptide. Uncovering this unusual flavin-dependent paradigm for thioether residue formation advances the understanding in the biosynthesis of aminovinyl-cysteine-containing RiPPs and renews interest in flavoproteins, particularly those involved in non-redox transformations. LanD-like flavoproteins activity, which is flexible in peptide substrate and amenable for evolution by engineering, can be combined with different post-translational modifications for structural diversity, thereby holding promise for peptide macrocyclization/functionalization in drug development by chemoenzymatic or synthetic biology approaches.

## INTRODUCTION

Flavin-dependent proteins are ubiquitous in living organisms and participate in numerous redox-related biochemical processes in all domains of life^1,2^. Increasing evidence indicates their critical roles in the biosynthesis of structurally-complex bioactive natural products, many of which are known to be ribosomally synthesized and post-translationally modified peptides (RiPPs)^3,4^. Each RiPP arises from post-translational modifications (PTMs) of a precursor peptide, which is typically composed of an N-terminal leader peptide (LP) and a C-terminal core peptide (CP). Only CP is transformed into mature product(s), usually in a manner dependent on LP. Using 20 proteinogenic amino acids as building blocks, nature creates stunningly diverse RiPPs largely because of unique enzymes for PTMs.

Aminovinyl-cysteine residues^3^, i.e., *S*-[(*Z*)-2-aminovinyl]-cysteine (AviCys) and *S*-[(*Z*)-2-aminovinyl]-3-methyl-cysteine (AviMeCys), are unusual thioether amino acids that occur in an array of RiPPs, including the lanthipeptide epidermin (EPI) and the antitumor non-lanthipeptides thioviridamides (TVAs) and cypemycin (CYP) (**Fig. 1**). These RiPPs share Avi(Me)Cys as part of a macrocyclic ring system that contains the C-terminal 4 or 6 residues of a precursor peptide, despite differences in their overall architectures and other residue characteristics. Knowledge concerning Avi(Me)Cys formation is poor and has gained primarily from investigations into related lanthipeptides^5-9^. Typically, it relies on processing two residues that are flanked by 2 or 4-aa, i.e., dehydration of an internal l-Ser/l-Thr, oxidative decarboxylation of the C-terminal l-Cys and the formation of a thioether linkage via a Michael addition between the resulting 2,3-didehydroalanine (Dha)/2,3-didehydrobutyrine (Dhb) and C-terminal enethiol^10^. In fact, lanthipeptides are characterized by the distinct thioether amino acids lanthionine (Lan) or 3-methyl-lanthionine (MeLan) residues (**Fig. 1**), each of which arises from dehydration of an l-Ser/l-Thr residue and subsequent cyclization that adds the side chain of an l-Cys residue onto the newly formed dehydroamino acid for macrocyclization^11^. The two reactions that occur in tandem can be attributed to the dedicated activities of a LanB dehydratase and a LanC cyclase for Class I lanthipeptides or to the combined activity of a bifunctional (Me)Lan synthetase, i.e., LanM, LanKC or LanL for Class II, III or IV lanthipeptides, respectively^11^.

**Figure 1.**
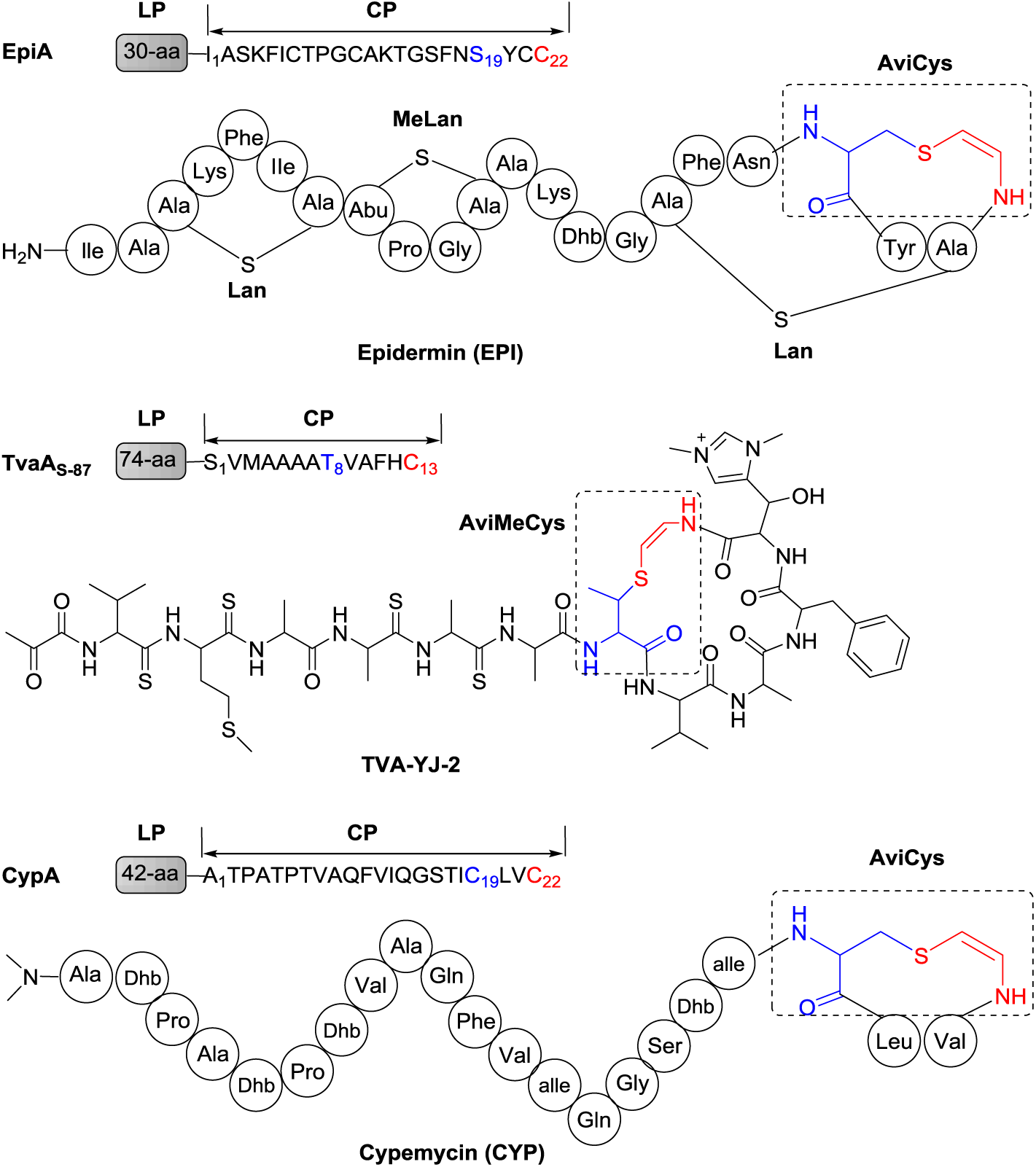
Precursor peptides and structures of Avi(Me)Cys-containing RiPPs EPI, TVA-YJ-2 and CYP. In each precursor peptide, where the LP and CP sequences are indicated, the C-terminal l-Cys and the internal l-Ser/l-Thr or l-Cys that are involved in Avi(Me)Cys formation are numbered and colored in red and blue, respectively. The Avi(Me)Cys residues that arise from processing the above two target residues are highlighted by color in dashed rectangle. Lan and MeLan residues in EPI are indicated. For other unusual residues: Dha, 2,3-didehydroalanine; Dhb, 2,3-didehydrobutyrine; alle, l-*alle*-isoleucine; and Abu, α-aminobutyric acid.

In the biosynthesis of lanthipeptides containing an Avi(Me)Cys residue, it has long been believed that the dehydration activity involved in (Me)Lan formation plays an additional role, by producing the Dha/Dhb residue lying 2 or 4-aa upstream of the C-terminal l-Cys. Transforming this l-Cys to an enethiol nucleophile depends on the activity of a LanD flavoprotein, which belongs to the family of homo-oligomeric flavin-containing cysteine decarboxylases (HFCDs)^10^. LanD activity for oxidative decarboxylation of the C-terminal l-Cys residue of a peptide was known over two decades ago^12, 13^; however, questions regarding Avi(Me)Cys formation remain in terms of timing this reaction and l-Ser/l-Thr dehydration and determining whether the following Michael addition occurs non-enzymatically or, if enzymatically, which enzyme acts as the catalyst, e.g., a LanD flavoprotein, a monofunctional LanC cyclase, or a LanM, LanKC or Lanl-represented bifunctional (Me)Lan synthetase^9,14^. Here, we report the unexpected dual activity of LanD-like flavoproteins in the biosynthesis of the non-lanthipeptides TVAs and CYP, where these flavoproteins can oxidatively decarboxylate and cyclize a peptide, either unprocessed or dehydrated/dethiolated, through the formation of an Avi(Me)Cys residue at the C-terminus. This study advance the understanding in the biosynthesis of different Avi(Me)Cys-containing RiPPs, where associated PTM logics and enzymatic mechanisms have not been established since they were discovered.

## RESULTS AND DISCUSSION

### TvaF_S-87_ can oxidatively decarboxylate and cyclize a peptide

We began investigating Avi(Me)Cys formation in the biosynthesis of the poorly understood non-lanthipeptides TVAs, a class of Avi(Me)Cys-containing thioamide RiPPs (**Fig. 1**)^15,16^. Recent survey of the published genome sequences revealed a number of new biosynthetic gene clusters (BGCs) that share nearly head-to-tail homology with the prototypal TVA BGC (*tva*) characterized in *Streptomyces olivoviridis*^*17*^, and subsequent product mining in selected microorganisms led to a rapid enrichment of the TVA family^18-22^, in which several members appear to be promising for cancer treatment. All these BGCs do not encode any homologs of LanC, LanM, LanKC or LanL proteins that possess cyclase activity, consistent with the absence of (Me)Lan in the structures of TVAs. Comparative analysis of the biosynthetic pathways of different Avi(Me)Cys-containing RiPPs revealed that only the LanD-like, HFCD-fold flavoprotein is common (**Supplementary Fig. 1**), indicating the importance of this type protein in the formation of the shared characteristic residue Avi(Me)Cys. To determine the function of this LanD-like flavoprotein in TVA biosynthesis, we overexpressed TvaF_S-87_ (201-aa, a homolog of TvaF from *Streptomyces sp*. NRRL S-87) in *Escherichia coli* (**Supplementary Fig. 2**).

The recombinant TvaF_S-87_ flavoprotein, which proved to non-covalently bind with flavin mononucleotide (FMN), was used to *in vitro* process the N-terminally truncated substrate **1**, a 21-aa-long, C-terminal mimic of the TVA precursor peptide TvaA_S-87_ (**Fig. 2** and **Supplementary Figs. 3** and **4**). This synthetic substrate contains the intact CP sequence of TvaA_S-87_, **S**_**1**_**VMAAAAT**_**8**_**VAFHC**_**13**_ (numbering relates to residue position in CP, as below), and the 8-aa sequence of LP adjacent to CP. The TvaF_S-87_-catalyzed conversion of **1** produced a −46 Da derivative, **1-I**. Analysis by high-performance liquid chromatography with high-resolution mass spectrometry (HPLC-HR-MS) confirmed that oxidative decarboxylation occurred on the C-terminal l-Cys to generate an enethiol, as previously observed in LanD catalysis for Avi(Me)Cys-containing lanthipeptide biosynthesis^9,12,13^. **1-I** is unstable and tends to be dethiolated to yield a dominant aldehyde derivative (**1-II**) via hydrolysis. Similar results were recently observed when coexpressing the TvaF and TvaA homologs in *E. coli*^23^. Different from this *in vivo* transformation, we observed an additional −64 Da product, **1-III**, in TvaF_S-87_ catalysis. The observation of **1-III** is unexpected, given the loss of molecular weight (MW) that indicates the occurrence of an additional H_2_O elimination (−18 Da). Likely, TvaF_S-87_ possesses more than the activity for oxidative decarboxylation of a peptide.

**Figure 2.**
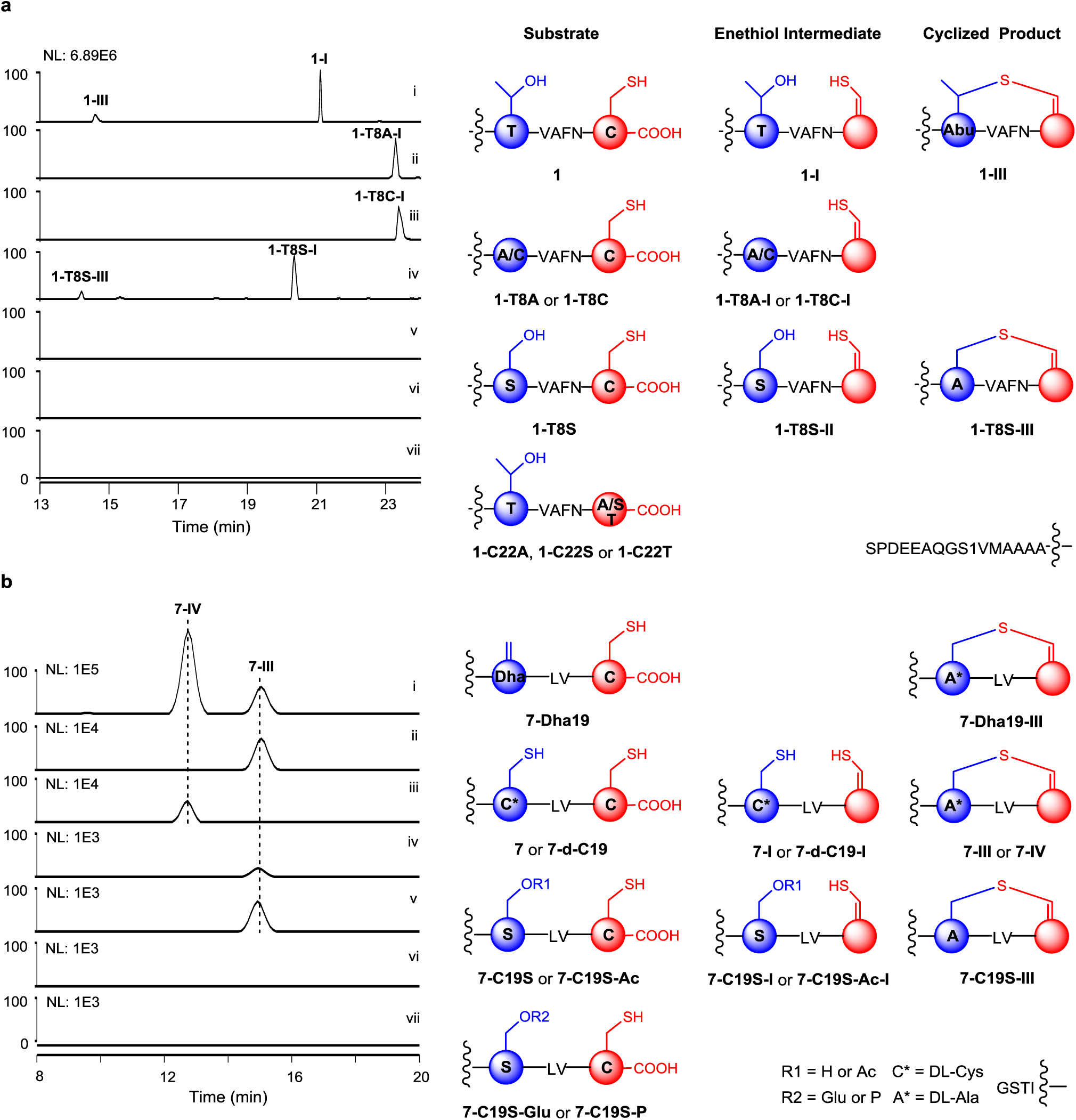
LanD-type flavoprotein-catalyzed Avi(Me)Cys formation. Left, HPLC-MS traces; and right, substrates and their associated enethiol intermediates and cyclized products. The two residues involved in Avi(Me)Cys formation, i.e., the C-terminal l-Cys (red) and its upstream l-Thr/l-Ser or l-Cys (blue), are shown in balls. Conversions were conducted at 30°C for 2 hr in the presence of TvaF_S-87_ or CypD. (**a**) *In vitro* assays of TvaF_S-87_ activity using the 21-aa-long, C-terminal CP sequence of the precursor peptide TvaA_S-87_ and its variants. i, **1**; ii, **1-T8A**; iii, **1-T8C**; iv, **1-T8S**; v, **1-C13A**; vi, **1-C13S**; and vii, **1-C13T**. (**b**) *In vitro* assays of CypD activity using the 8-aa-long, C-terminal CP sequence of the precursor peptide CypA and its variants. i, **7-Dha19**; ii, **7**; iii, **7-d-C19**; iv, **7-C19S**; v, **7-C19S-Ac**; vi, **7-C19S-Glu**; and vii, **7-C19S-P**. For details of the HR-MS and HR-MS/MS data, **Supplementary Fig. 6b** and **Table 4**.

Based on HR-MS/MS analysis, the difference of **1-III** from **1, 1-I** and **1-II** was narrowed to the C-terminal sequence **MAAAAT**_**8**_**VAFHC**_**13**_, in which only l-Thr8 can be the target residue subjected to H_2_O elimination (**Supplementary Fig. 4** and **Table 4**). Specifically, no fragment ions were observed from the C-terminal **T**_**8**_**VAFHC**_**13**_ sequence of **1-III**, and corresponding fragment ions were indeed found when analyzing **1, 1-I** and **1-II**, all of which are linear. Furthermore, **1-III** was resistant to thiol derivatization with N-ehtylmaleimide (NEM); in contrast, **1** and **1-I**, both of which bear a free (ene)thiol group, were sensitive to this treatment, yielding the derivatives conjugated by one NEM molecule (**Supplementary Fig. 5**). These findings supported the notion that **1-III** is a cyclized product resulting from the dual activity of TvaF_S-87_, which can selectively process both l-Cys13 and l-Thr8 and form an AviMeCys linkage. During this process, **1-I** can be released from TvaF_S-87_ as a linear enethiol intermediate.

To further determine the tolerance of TvaF_S-87_ catalysis to changes of the target residues, **1** was subjected to site-specific mutagenesis, resulting in the substrate variants **1-T8A, 1-T8S, 1-T8C, 1-C13A, 1-C13S** and **1-C13T** (**Fig. 2** and **Supplementary Fig**.**3** and **Table 4**). The mutations C13A, C13S and C13T caused no substrate conversion. The mutations T8A, T8S and T8C did not affect l-Cys13 processing, leading to the production of the enethiol intermediates **1-T8A-I, 1-T8S-I** and **1-T8C-I**, respectively. Only **1-T8S-I** can be further transformed to the cyclized product, **1-T8S-III**, and form a distinct AviCys residue. Clearly, TvaF_S-87_ activity tolerates the change of l-Thr8 to the chemically similar residue l-Ser. We then assayed TvaF_S-87_ activity using a series of further shortened mimics of TvaA_S-87_ CP, i.e., **2** (12-aa), **3** (11-aa), **4** (10-aa), **5** (8-aa) and **6** (6-aa, **T**_**8**_**VAFHC**_**13**_), all of which lack the LP sequence and vary in truncation extent at the N-terminus of CP (**Supplementary Fig. 3** and **Table 4**). Consequently, both the −46 Da enethiol intermediate (**2-I** or **3-I**) and the −64 Da cyclized product (**2-III** or **3-III**) were observed in the transformation of **2** or **3**; however, the yields were much lowered. Although a trace of enethiol product (−46 Da, **4-I, 5-I** or **6-I**) was detected when **4, 5** or **6** was used as the substrate, coupling l-Thr8 via H_2_O elimination-associated cyclization appears to occur only in the presence of a > 4-aa upstream flanking sequence. Therefore, the upstream residues of Avi(Me)Cys, either from LP or CP, appear to be less dependent but do have impact on the efficiency and completeness of TvaF_S-87_ catalysis.

### CypD shares the ability for AviCys formation

To examine the generality of LanD-like flavoprotein activity for distinct non-lanthipeptides, we *in vitro* assayed the activity of CypD (190-aa) involved in the biosynthesis of the CYP. This RiPP is an AviCys-containing linaridin-type RiPP that possesses several l-Thr-derived Dhbs but lacks MeLan residues characteristic of lanthipeptides (**Fig. 1**)^24,25^. Similar to *tva*, the CYP BGC (*cyp*) does not contain any genes encoding the homologs of proteins with (Me)Lan cyclase activity (**Supplementary Fig. 1**). The biosynthesis of this non-lanthipeptide is particularly interesting because two l-Cys residues of the precursor peptide are processed into an AviCys residue. We overexpressed CypD in *E. coli* (**Supplementary Fig. 2**). Different from TvaF_S-87_, purified CypD was observed to bind non-covalently with the cofactor flavin adenine dinucleotide (FAD). Using the synthetic substrate **7** (**GSTIC**_**19**_**LVC**_**22**_), an N-terminally truncated 8-aa CP sequence of the precursor peptide CypA, CypD catalysis produced a −46 Da enethiol intermediate **7-I** (along with its dethiolated derivative **7-II**) and, particularly, a −80 Da cyclized product, **7-III**, in a manner similar to that found in TvaF_S-87_ catalysis (**Figs. 2** and **3** and **Supplementary Figs. 3,6a** and **Table 4**). The structural differences between **7-I** and **7-III** were further distinguished by using NEM for thiol derivatization (**Supplementary Fig. 7**). Similar to the substrate **7**, the intermediate **7-I** contains two (ene)thiol moieties and can be conjugated by two NEM molecules. The product **7-III** cannot be modified and thus has no free thiol group. The formation of an AviCys residue was first indicated by HR-MS/MS analysis, which revealed a specific ion fragment ([M+H]^+^ *m/z*: calcd. 258.1271, obs. 258. 1271) containing this residue (**Fig. 3** and **Supplementary Table 4**). Then, the CypD-catalyzed transformation of **7** was scaled up, leading to the accumulation of **7-III** in ∼1000 mL of the reaction mixture for structural elucidation. Consequently, selected Nuclear Magnetic Resonance (NMR) analyses of purified **7-III** supported the presence of an AviCys residue (**Fig. 3** and **Supplementary Fig. 8** and **Table 6**). Together, the above results indicate that resembling TvaF_S-87_, CypD catalyzes oxidative decarboxylation and cyclization of a peptide to form an AviCys linkage along with H_2_S elimination.

**Figure 3.**
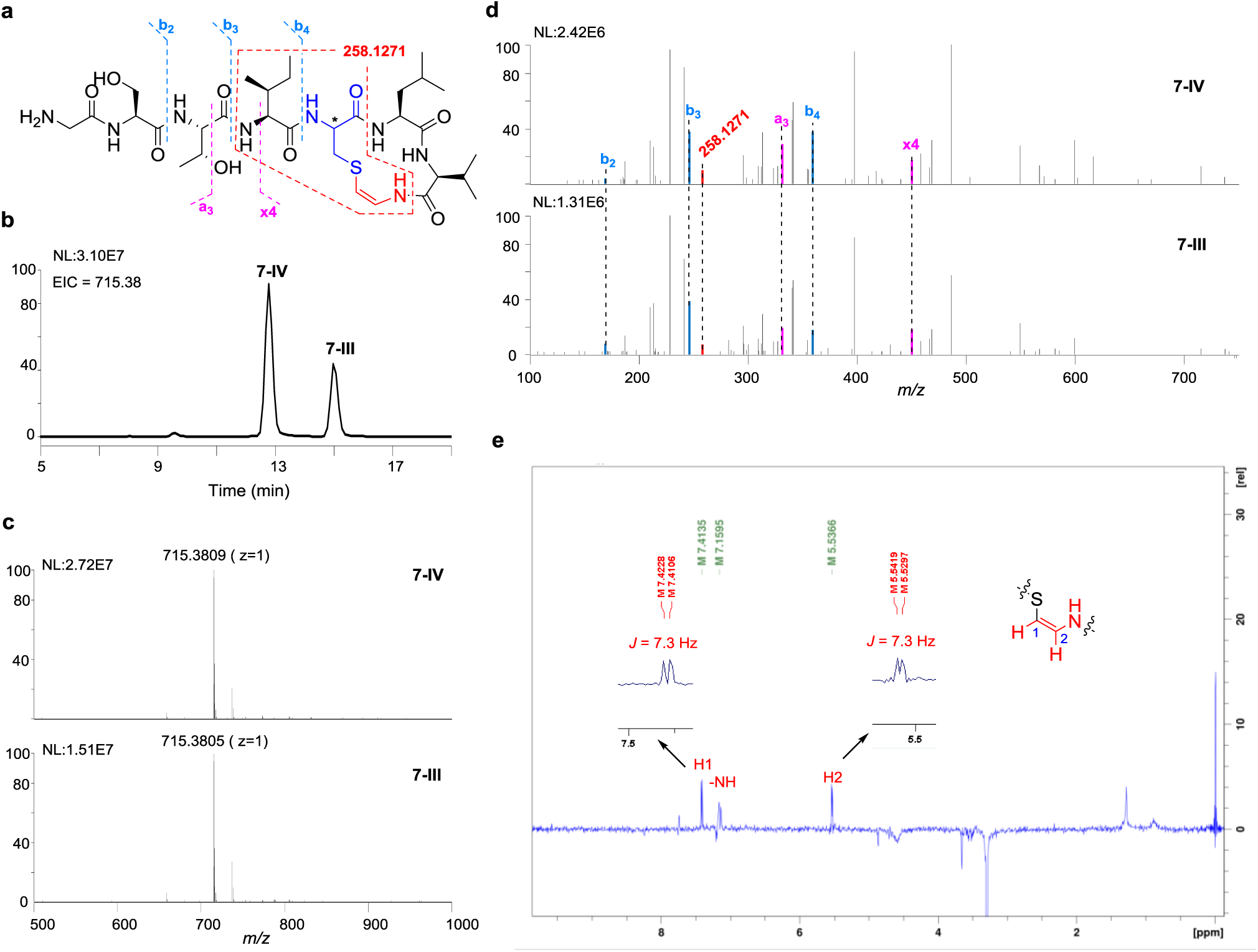
Spectral analysis of the cyclized AviCys-containing products **7-III** and **7-IV. (a)** Structures of **7-III** and **7-IV**. Fragmentations by MS/MS are indicated. The carbon atom of which **7-III** and **7-IV** are different in chirality is labeled with asterisk. (**b**) HPLC-MS. (**c**) HR-MS. (**d**) MS/MS. (**e**) Selected ^1^H NMR signals for **7-III**. 2D TOCSY f1 slice at f2 chemical shifts (600 MHz, DMSO-*d*_6_) of H-1 of **7-III** and the key of spin-spin coupling constant (*J*_H1/H2_ = 7.3 Hz) was highlighted. For details of the spectral data, see **Supplementary Fig. 8** and **Tables 4** and **6**.

To examine the tolerance of the target residues in CypD catalysis, we synthesized **7** variants, i.e., **7-C19A, 7-C19S** and **7-C19T**, by mutating the internal l-Cys19, and **7-C22A, 7-C22S** and **7-C22T**, by mutating the C-terminal l-Cys22, respectively (**Supplementary Fig. 3** and **Table 4**). In the presence of CypD, both **7-C19A** and **7-C19T** underwent oxidative decarboxylation to produce −46 Da enethiols (i.e., **7-C19A-I** and **7-C19T-I**). Only **7-C19S** can be transformed further into the cyclized product **7-III** (−64 Da, in a yield ∼50-fold lower than that in the transformation of **7**) under the same reaction conditions, while a distinct enethiol intermediate, **7-C19S-I** (−46 Da), accumulated. Clearly, l-Ser can substitute for the internal l-Cys19 during AviCys formation, indicating that in this CypD catalysis, l-Ser shares with l-Cys a common mechanism for H_2_O or H_2_S elimination. CypD did not transform the variants **7-C22A, 7-C22S** and **7-C22T**, again validating the necessity of the C-terminal l-Cys.

### TvaF_S-87_ and CypD are dodecameric Rossmann-fold flavoproteins

To gain structural insights into the flavoproteins TvaF_S-87_ and CypD for non-lanthipeptide biosynthesis, we determined their apo-form crystal structures for comparative analysis (to 2.27 Å and 2.38 Å resolutions for TvaF_S-87_ and CypD, respectively) (**Fig. 4** and **Supplementary Table 7**). Despite low homology observed in sequence, TvaF_S-87_ and CypD are highly similar in overall structure to each other and to characterized LanD proteins (**Figs. 4a-e** and **Supplementary Fig. 9a-e**) (e.g., EpiD and MrsD in the biosynthesis of the lanthipeptides EPI and mersacidin^26,27^, respectively). Each protein forms a stable dodecamer composed of four trimeric units that are located at the vertices of a tetrahedron. In a trimer unit, each Rossmann-fold monomer associates with a flavin cofactor (**Supplementary Figs. 10** and **11**). This factor is located in a cavity at the interface of two monomers, of which one plays a predominant role for hydrophobic contacts and polar interactions, facilitating packing with the third monomer. Compared to CypD, which binds FAD, TvaF_S-87_ is more similar to the archetypal LanD protein EpiD largely because they share a manner to accommodate the FMN cofactor (**Supplementary Fig. S9b, c**). Recently, two research groups reported similar observations based on the crystallization of CypD and a different TvaF homolog, respectively^23,28^. Notably, we observed two slightly different conformations of the monomers in the crystal structure of TvaF_S-87_ (**Supplementary Fig. S9f**). The region connecting β5 and α9 is disordered and unsolved in some monomers. Whereas in others, it forms a short anitparallel β-sheet resembling that as part of the substrate-binding clamp observed in the EpiD complex with a peptidyl substrate mimic (PDB ID: 1G5Q) (**Supplementary Fig. 9g**), suggesting that TvaF_S-87_ and EpiD adopt a similar mode for substrate-binding in the first oxidative decarboxylation step.

**Figure 4.**
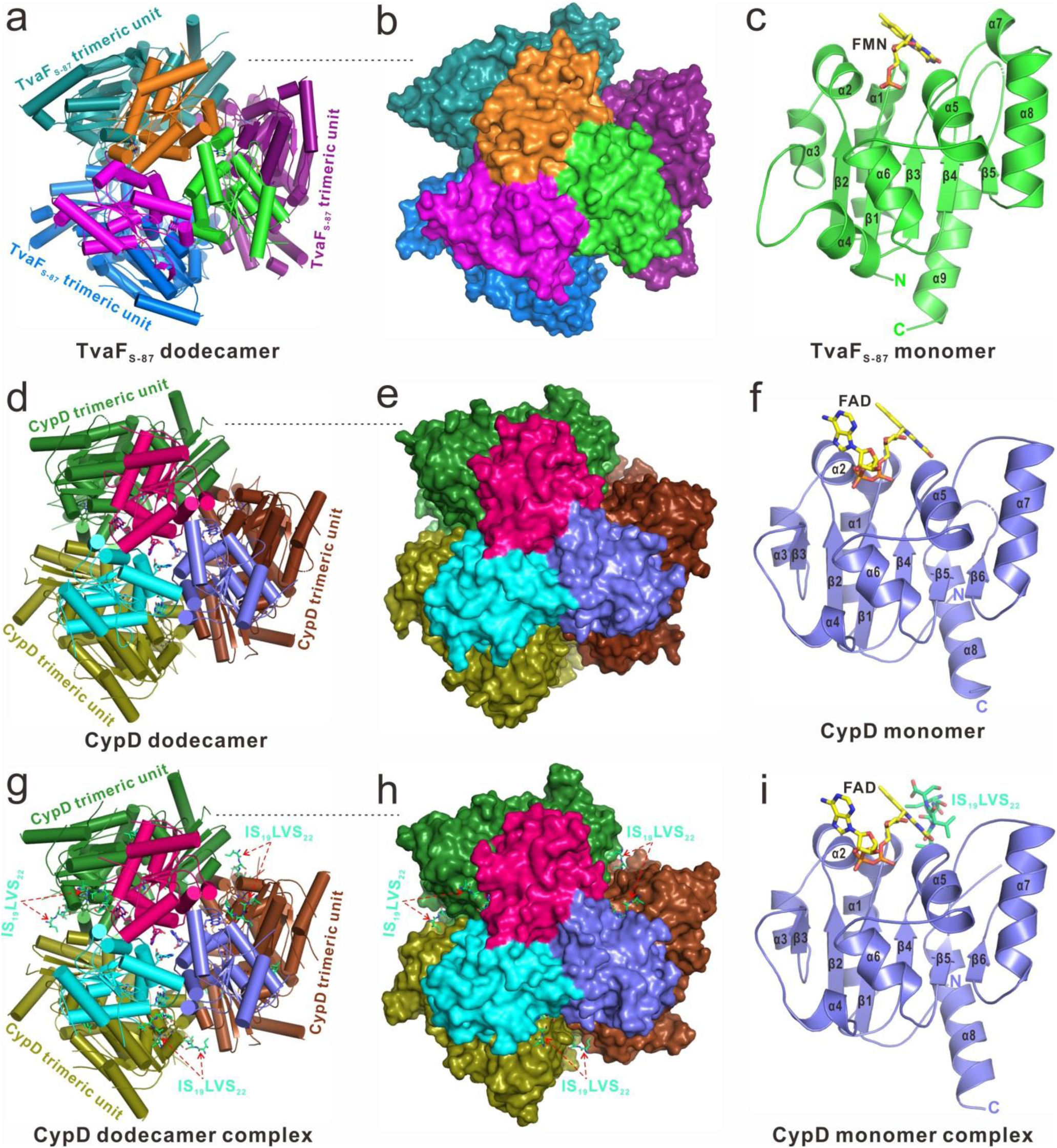
Crystal structures of LanD-like flavoproteins. For TvaF_S-87_ (**a-c**) or CypD (**d-f**) in apo-form, the dodecamer is shown by viewing along a 3-fold axis and in a ribbon-stick diagram (left) or surface representation (middle), while the monomer in complex with FMN or FAD (stick model) is shown by ribbon diagram (right). For CypD in complex with **IS**_**19**_**LVS**_**22**_ (the C-terminal pentapeptide sequence of **8, g-i**), the dodecamer complex is shown in a ribbon-stick diagram (left) and surface representation (middle), and the monomer in complex with FAD and **IS**_**19**_**LVS**_**22**_ (stick model) is shown by ribbon diagram (right).

CypD and TvaF_S-87_ share a great majority of the residues responsible for interaction with the flavin cofactor, either FMN or FAD, through hydrophobic contacts and hydrogen bonds with the common isoalloxazine ring (**Supplementary Figs. 10-12**). In their structures, interaction between trimeric units are attributed to residues within the α1, α2, and α3 regions of the monomer (**Supplementary Figs. 12-14**). We selectively mutated the key residues V28 and M62, respectively, to l-Asp in these regions to destabilize this interaction in TvaF_S-87_. The two mutations exclusively led to the decomposition of the active TvaF_S-87_ dodecamer to stable trimer, which retains the ability to bind FMN but losses the activity to transform the peptide substrate **1** (**Supplementary Figs. 4** and **13c**). Thus, the formation of a dodecamer is necessary for the activity of LanD-like flavoproteins. This structural nature cannot be applied to 4’-phosphopantothenoylcysteine (PPC) decarboxylases, which fall into the same HFCD family but function in an active trimer form during the biosynthesis of coenzyme A^29^.

### Capture of a cyclization transition state in the CypD complex

After numerous attempts, we eventually resolved the co-crystal structure of CypD in complex with **8** (**GSTIS**_**19**_**LVS**_**22**_) to 2.3 Å resolution (**Fig. 4g-i** and **Supplementary Table 8**). **8** is a synthetic derivative of the peptide substrate **7** and fused with CypD by a GS-repeat linker. Compared to **7**, this derivative, in which the two residues involved in AviCys formation (i.e., l-Cys19 and l-Cys22) are replaced with l-Ser, cannot be transformed by CypD. The complex of CypD highly resembles its apo-form, as twelve FAD-associated monomers adopt a similar conformation for assembly to a homododecamer accommodating twelve **8** through the formation of four trimeric units (**Fig. 4d-i** and **Supplementary Fig. 15a**). For each bound octapeptide, while the N-terminal tripeptide **GST** cannot be modeled due to a lack of electron density, the remaining pentapeptide **IS**_**19**_**LVS**_**22**_ can be clearly defined. The **IS**_**19**_**LVS**_**22**_ sequence adopts an obvious bent conformation and packs extensively with a solvent-exposed, highly charged pocket formed between two trimeric units (**Fig. 5a** and **Supplementary Fig. 16**). This pocket, which is close to a bound FAD cofactor, overlaps the putative substrate binding region revealed by superimposition of CypD and the mutant EpiD (H67N) complex with a pentapeptide substrate mimic (**Supplementary Fig. 17**)^26^.

**Figure 5.**
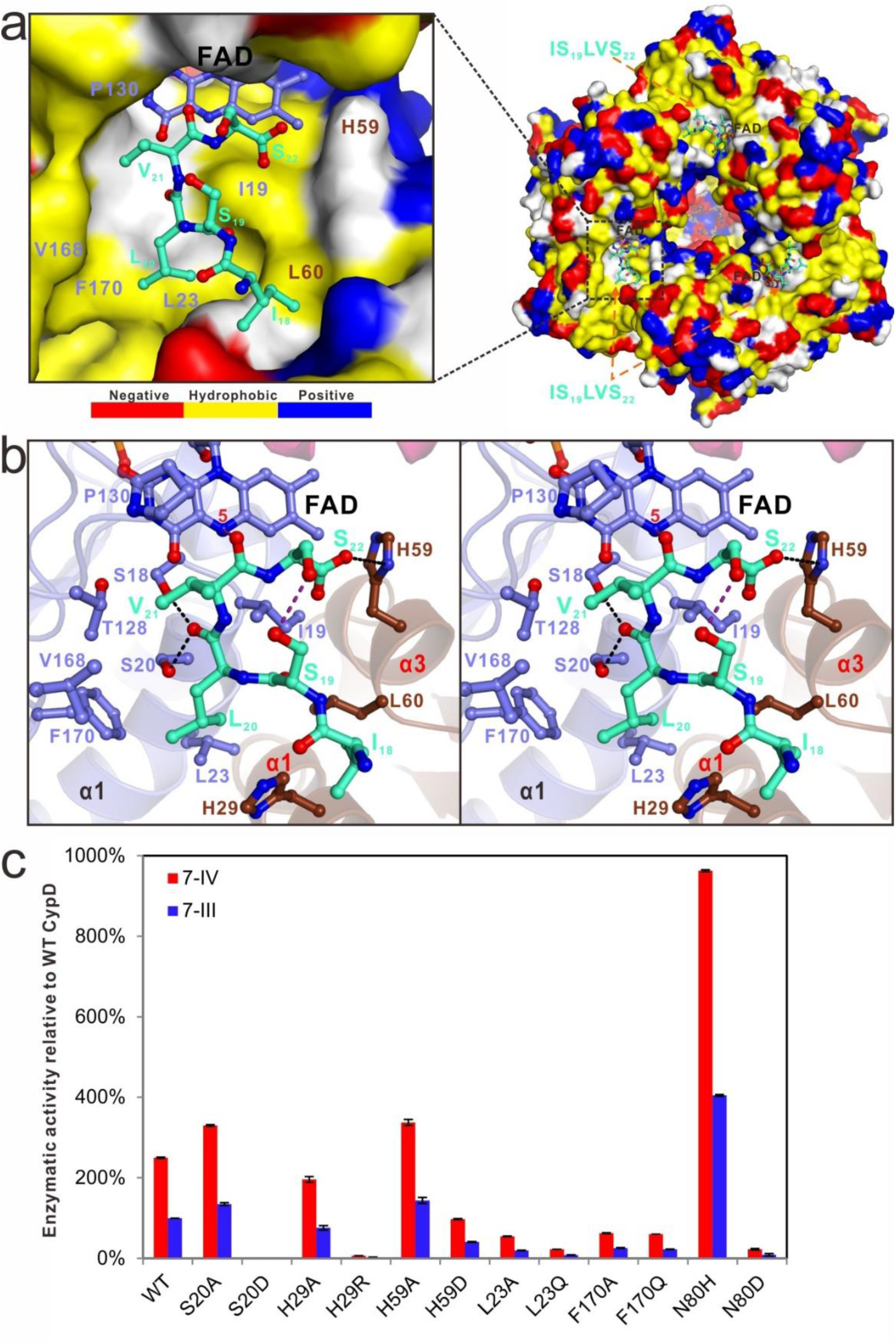
Identification of the residues involved in CypD catalysis at its active site. (**a**) The combined surface representation and the stick-ball model showing the hydrophobic binding interface between CypD and **IS**_**19**_**LVS**_**22**_. The FAD cofactor and bound **IS**_**19**_**LVS**_**22**_ are displayed in the stick-ball model, and the CypD dodecamer in the surface view is colored by amino acid types. In the surface model of CypD, the hydrophobic amino acid residues are drawn in yellow, the positively charged residues in blue, the negatively charged residues in red, and the uncharged polar residues in gray. (**b**) Stereoviews of the ribbon-stick models showing the detailed interactions of CypD with **IS**_**19**_**LVS**_**22**_ in the complex. The hydrogen bonds involved in the interaction are shown as dotted lines. The N5 atom on the isoalloxazine ring of FAD and the putative hydrogen bond between the backbone carboxyl group of V_21_ and reduced N5 atom are also indicated. (**c**) Comparison of the enzymatic activities of CypD with its variants based on examination of the production of **7-III** and **7-IV**.

In the CypD complex (**Fig. 5**), the hydrophobic side chains of the residues l-Val21, l-Leu20, and l-Ile18 of **8** occupy a hydrophobic groove formed by the residues I19, L23, T128, P130, V168 and F170 of one monomer and the residue L60 of another monomer within the neighboring trimeric unit. The negatively charged, C-terminal carboxyl group of **8** forms a weak hydrogen bond with the monomer residue H59, while the backbone carboxyl group of l-Leu20 interacts with the polar side chains of the monomer residues S18 and S20 via two highly specific hydrogen bonds. Theses interactions allow approaching between the polar side chains of l-Ser22 and l-Ser19 to form an intramolecular hydrogen bond in **8**, where the bent conformation is further stabilized (**Fig. 5b**). This conformation leads to a hypothesis that during the transformation of **7**/**7-C19S**, cyclization to form an AviCys residue in CypD catalysis proceeds via nucleophilic substitution between l-Cys19/l-Ser19 and the C-terminal enethiol resulting from oxidative decarboxylation. Distinct from the mutant EpiD (H67N) complex, the CypD complex with **8** unlikely restores a transition state for l-Cys22 oxidative decarboxylation given their significant differences in substrate binding. Instead, enethiol production might be bypassed (assumedly due to replacing the C-terminal l-Cys with l-Ser) in this CypD complex, where a transition state for the following cyclization reaction was captured. The mechanism for CypD-catalyzed, l-Cys22 oxidative decarboxylation was recently proposed to be similar to that in EpiD catalysis^30^; thus, repositioning the enethiol intermediate at the active site of CypD appears to be necessary for nucleophilic substitution. In the CypD complex with **8**, the backbone carboxyl group of the substrate residue l-Val21 approaches N5 of the isoalloxazine ring of a bound FAD within a distance of 3.5 Å (**Fig. 5b**). The reduction of FAD to FADH_2_ during l-Cys22 processing can protonate N5 and facilitate the formation of a hydrogen bond with the backbone carboxyl group of l-Val21. This polar interaction might compensate the loss of the weak hydrogen bond between the monomer residue H59 and the C-terminal l-Ser22 of **8** and thus can stabilize the bent conformation of the decarboxylated enethiol intermediate.

To validate this hypothesis, we assayed CypD activity using the substrate **7-C19S** in which l-Ser19 was deuterium-labeled (i.e., l-[2,3,3-D_3_]Ser). As expected, −46 Da intermediate **7-C19S-I** and −64 Da product **7-III** were observed (**Supplementary Fig. 18**). They both share +3 Da in MW compared to those produced in the control reaction where unlabeled l-Ser was used. The observation of +3 Da **7-III** strongly supported that following C-terminal l-Cys22 processing, coupling l-Ser19 with the newly formed enethiol proceeds via nucleophilic substitution because all deuterium atoms were retained. Most likely, TvaF_S-87_ shares the mechanism for Avi(Me)Cys formation in the transformation of **1**/**1-T8S**.

### Determination of the nature of CypD substrate in CYP biosynthesis

Although the dual activity of TvaF_S-87_ and CypD is experimentally and structurally evidenced, it is unlikely that these flavoproteins act on an unmodified peptide substrate for Avi(Me)Cys formation in the biosynthesis of TVAs or CYP because of the low yield of cyclized products versus the relatively high yield of enethiol intermediates (for calculation, their aldehyde derivative were included). To examine PTM dependence, we assayed CypD activity using the variants of **7**/**7-C19S** (i.e., **7-C19S-Ac, 7-C19S-Glu, 7-C19S-P** and **7-Dha19**) in which l-Cys19/l-Ser19 was replaced with acetylated l-Ser, glutamylated l-Ser, phosphorylated l-Ser and Dha, respectively (**Fig. 2**). Acetylation, glutamylation and phosphorylation can form a leaving group, which might facilitate cyclization. While CypD failed to convert **7-C19S-Glu** and **7-C19S-P** likely due to the steric or electronic effects that arise from glutamyl or phosphoryl substitution, this flavoprotein tolerated the substrate **7-C19S-Ac** and produced **7-III** with a yield ∼2-fold higher than that in the conversion of the unmodified substrate **7-C19S**. In both conversions, the yields of the enethiol intermediates were comparable (**Supplementary Fig. 6b**), indicating that l-Ser acetylation does motivate enethiol coupling.

In contrast, the CypD-catalyzed conversion of **7-Dha19** produced **7-III** with a yield much higher than those in the conversions of unmodified **7** (by ∼10-fold) and **7-C19S** (by ∼500-fold). Notably, this conversion resulted in an additional product, **7-IV**, which was even ∼2.5-fold higher than **7-III** in yield (**Fig. 2**). **7-IV** and **7-III** were validated to be identical to each other in MW, MS/MS analysis, AviCys-containing and chemical derivatization (**Fig. 2** and **Supplementary Fig. 19**), suggesting that they are diastereomers different from each other in the Cα configuration of the AviCys residue. **7-III** is believed to have an l-AviCys residue because nucleophilic substitution does not change the configuration at Cα; thereby, **7-IV** might have a D-AviCys residue. This prediction was further supported by the success in converting the substrate **7-d-C19** (in which l-Cys19 was replaced with D-Cys) to produce **7-IV** only (**Fig. 2** and **Supplementary Fig. 6b**). Remarkably, enethiol intermediate (i.e., **7-Dha19-I**) did not accumulated during the CypD-catalyzed conversion of **7-Dha19**, indicating that Michael addition to the Dha19 residue immediately follows oxidative decarboxylation of the C-terminal l-Cys22 and occurs efficiently for cyclization. Together, CypD activity prefers a dethiolated/dehydrated substrate (rather than hijacking the l-Cys19/l-Ser19-functionalized intermediate) and tends to produce a D-AviCys-containing product. In fact, D-AviCys residue was found exclusively in related natural products that were characterized in stereochemistry^31,32^. The above results argue against the recent report by Zhang *et al*. who failed to observe AviCys formation in the conversion of a Dha-containing peptide and thus concluded that CypD has no cyclase activity^33^. In this study, two l-Lys residues were added at the N-terminus of the 7-aa-long, C-terminal CP mimic of CypA, yielding **KKSTI-Dha**_**19**_**-LVC**_**22**_, which CypD can oxidatively decarboxylate but cannot cyclize. Instead, this finding is consistent with that observed in aforementioned TvaF_S-87_ catalysis, i.e., cyclization is an enzymatic process and the sequence upstream of those involved in Avi(Me)Cys formation are necessary for the completeness of CypD catalysis.

### Examination of the active site residues in CypD

Based on the crystal structure of CypD in complex with **8**, we first utilized **7-Dha19** as the substrate to examine the involvement of the active site residues in CypD catalysis (**Fig. 5**). While the mutation S20D completely abolished CypD activity, the mutations L23A, L23Q, F170Q, F170A, H59D and H29R, which change the physiochemical and/or steric properties of residues, significantly lowered the yields but did not alter the ratio between the diastereomers **7-III** and **7-IV** (∼1:2.5) (**Fig. 5c**). No enethiol intermediates (and their aldehyde derivatives) accumulated in these assays. Given the fact that CypD employs a same pocket for accommodating **7-Dha19** for oxidative decarboxylation and the enethiol intermediate for subsequent Michael addition, the above residues might contribute to the both reactions for substrate/enethiol intermediate binding during AviCys formation. Processing the C-terminal l-Cys22 may serve as the rate-limiting step, for which CypD and TvaF_S-87_ might share a flavin-dependent mechanism with the archetypal LanD protein EpiD^10^. At the active site, the peptide substrate approaches flavin via the thiol group of the C-terminal l-Cys, allowing oxidation occurring to initiate decarboxylation and form a reactive enethiolate ion. During this process, the residues l-His67 in EpiD could act as a base and participate in C-terminal l-Cys activation. Consistently, the TvaF_S-87_ mutant in which the corresponding residue H85 is changed to l-Ala completely lost activity (**Supplementary Fig. 4**). In contract, CypD possesses a distinct N80 according to sequence alignment (**Supplementary Fig. 12**). Whereas the mutation N80D nearly completely abolished **7-III**/**7-IV** production, the mutation N80H resulted in a ∼4-fold increase of the both cyclized products, indicating that a stronger base facilitates the deprotonation of the C-terminal l-Cys (**Fig. 5c**). The mutations H59A and S20A also improved the yields of **7-III** and **7-IV** (by ∼1-fold for each), supporting that the hydrophobicity of the binding pocket can affect the interaction with the substrate/ennethiol intermediate (**Fig. 5**). We then assayed the activities of the above CypD mutants by using the unprocessed substrate **7**. Overall, these mutants produced **7-III** and the enethiol intermediate/aldehyde derivative (**7-I**/**7-II**) with a similar yield trend (**Supplementary Fig. 20**), indicating that the cyclization activity of CypD or its variants can be examined using the readily prepared, unmodified peptide substrates (albeit lower in yields). Therefore, the proximity effect resulting from the bent conformation at the active site play a critical role in coupling l-Cys19 and l-Cys22 for AviCys formation.

## CONCLUSION

Following an unexpected finding that LanD-like flavoproteins can function alone to cyclize a peptide, we uncovered a new flavin-dependent PTM paradigm for the formation of thioether amino acids occurring in non-lanthipeptides (**Fig. 6**). We discovered this paradigm by assaying TvaF_S-87_ activity in the biosynthesis of the thioamide RiPPs TVAs, and subsequently validated its generality upon more detailed biochemical and structural investigations into CypD in the biosynthesis of the linaridin RiPP CYP. LanD-like flavoprotein first catalyzes oxidative decarboxylation of the invariable C-terminal l-Cys residue; then, it adds the resulting enethiol nucleophile to the 2 or 4-aa upstream internal residue, which is variable and can be l-Ser, l-Cys or l-Thr, and forms an Avi(Me)Cys linkage for peptide cyclization. This cyclization is an enzymatic process and occurs largely due to proximity effect at the active site of the enzyme by repositioning and forcing the enethiol intermediate to a bent conformation. Mechanistically, it can be substrate-dependent. To process an unmodified peptide, the cyclization goes inefficiently through nucleophilic substitution that causes the accumulation of the enethiol intermediate, and the upstream LP and CP sequences can affect the catalytic efficiency. In contrast, cyclizing a Dha-containing peptide, as shown in CypD catalysis where reduced flavin cofactor appears to be necessary to stabilize the bent conformation of the enethiol intermediate, proceeds efficiently via Michael addition to produce two diastereomers, of which the product with a D-AviCys residue is dominant. Consistent with the finding that CypD catalysis favors a dethiolated substrate, a Dha19-containing peptide intermediate was observed by heterologous overexpression of the *cyp* gene cluster in which *cypD* was inactivated^25^. TvaF and CypD are capable of oxidatively processing the C-terminal l-Cys and non-oxidatively coupling the internal residue, raising a question of whether LanD proteins share this dual activity in the biosynthesis of related lanthipeptides, where (Me)Lan cyclase activity has long been suggested to participate in Avi(Me)Cys formation. Recently, the combined activities of LanKC and LanD homologs proved to be necessary to form an AviCys-like avionin motif in the biosynthesis of lipolanthines; however, it remains to determine the individual roles of the either homologs in this process^34^. The study presented here renews great interest in flavoproteins, as their catalytic mechanisms have not been fully appreciated in various biochemical processes, particularly those involving additional non-redox reactions^35-37^. The LanD-like flavoproteins tolerate a wide variety of peptide substrates, is evolvable by engineering, and can be combined with different PTM enzymes for structural diversity. Using chemoenzymatic or synthetic biology strategies, enzymatic Avi(Me)Cys formation has potential for peptide drug development by application in peptide macrocyclization and functionalization, which remain a challenge to current chemical synthesis approaches.

**Figure 6.**
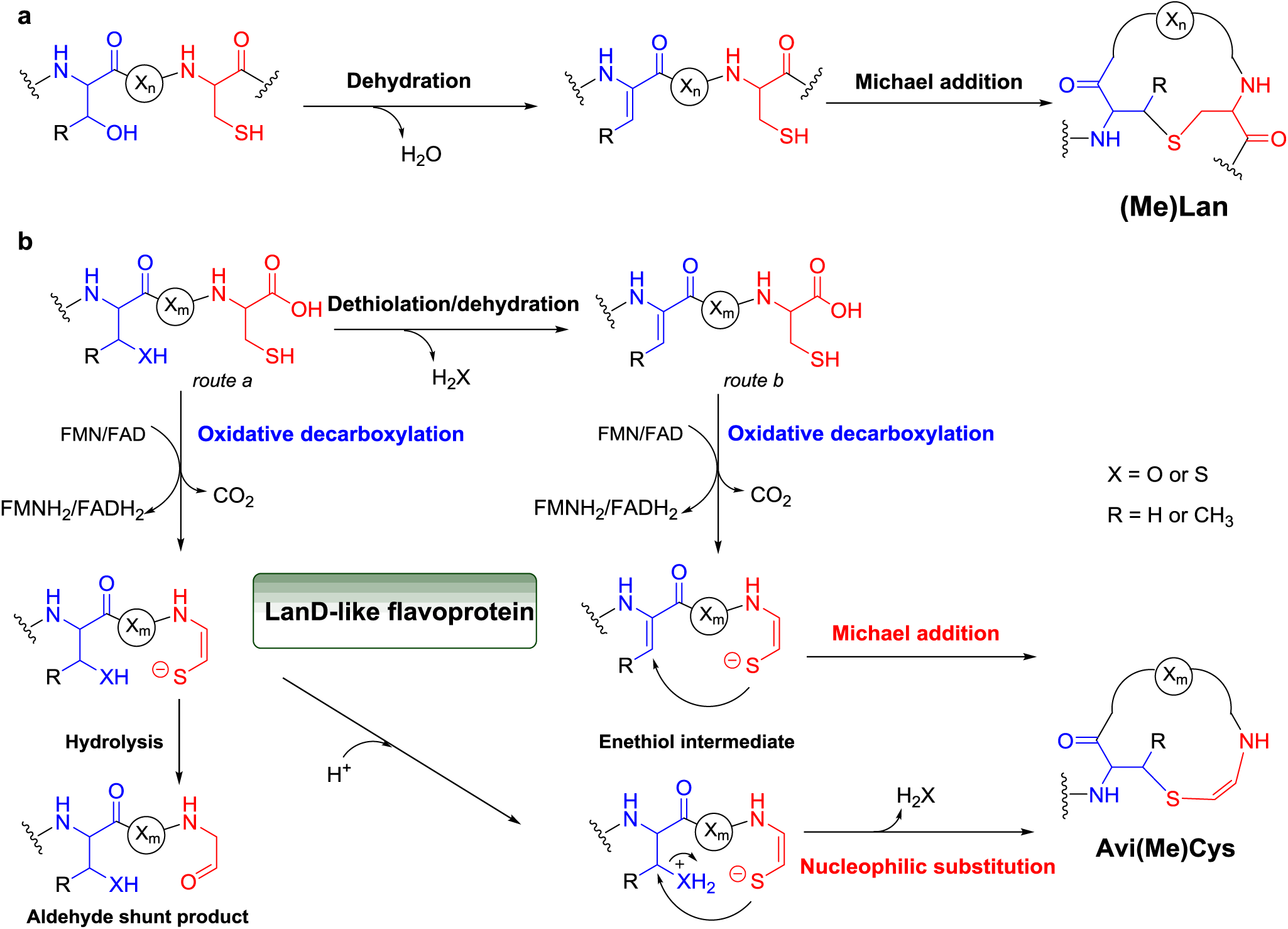
Mechanistic comparison in the formation of the thioether amino acids. (**a**) Formation of (Me)Lan residues. (**b**) Formation of Avi(Me)Cys residues. The cyclization activity of bifunctional LanD-like flavoproteins in the biosynthesis of non-lanthipeptides can be substrate-dependent, i.e., nucleophilic substitution for processing an unmodified peptide (*route a*) and Michael addition for processing a Dha-containing peptide (*route b*).

## Supplementary Information

is available in the online version of the paper.

## Data availability

Atomic coordinates and structural factors for the reported crystal structures of TvaF_S-87_ and CypD are deposited in the Protein Data Bank under the accession numbers 6KTP, 6KTT, 6KTI, and 6KT9.

## Acknowledgements

We thank SSRF BL19U1 for X-ray beam time. This work was supported in part by grants from NSFC (21672253, 21822705, 21621002), STCSM (17JC1405200), CAS (XDB20020200) (for L.P.); and NSFC (21750004 and 21520102004), CAS (QYZDJ-SSW-SLH037), STCSM (17JC1405100), MST (2018ZX091711001-006-010) and K. C. Wang Education Foundation (for W.L.).

