## Supplementary Materials for "LanD-like Flavoprotein-Catalyzed Aminovinyl-Cysteine Formation through Oxidative Decarboxylation and Cyclization of a Peptide at the C-Terminus"

**Supplementary Methods**

1.1 General Materials and Methods.

1.2 Protein Expression and Purification.

1.3 *In vitro* Assays of LanD-like Flavoprotein Activity.

1.4 Peptide Purification and Characterization

1.5 Protein crystallization and related structural elucidation.

1.6 Chemical synthesis of **7-Dha19**.

**Supplementary Figures**

**Supplementary Figure 1.** Alignment of the biosynthetic gene clusters of TVAs, CYP and EPI.

**Supplementary Figure 2.** Purified recombinant enzymes and cofactor analysis.

**Supplementary Figure 3.** Summary of LanD-like flavoprotein-catalyzed transformations in this study.

**Supplementary Figure 4.** HPLC-MS analysis of enethiol intermediates and associated shunt aldehyde derivatives in the $TvaA_{S\text{-}87}$-catalyzed conversions of **1** and its variants.

**Supplementary Figure 5.** Characterization of (ene)thiol and aldehyde peptides in the $TvaF_{S\text{-}87}$-catalyzed conversion of **1** by chemical derivatization.

**Supplementary Figure 6.** HPLC-MS analysis of enethiol intermediates and associated shunt aldehyde derivatives in the CypD-catalyzed conversions of **7** and its variants.

**Supplementary Figure 7.** Characterization of (ene)thiol peptides in the CypD-catalyzed conversion of **7** by chemical derivation.

**Supplementary Figure 8.** 2D-TOCSY f2-slices at related f1 experiments (600 MHz, DMSO-$d_6$) for residue identification in **7-III**.

**Supplementary Figure 9.** Overall structural comparison of the monomers of $TvaF_{S\text{-}87}$, CypD, MrsD and the EpiD complex (shown by ribbon diagram).

**Supplementary Figure 10.** FMN binding interface in the $TvaF_{S\text{-}87}$ structure.

**Supplementary Figure 11.** FAD binding interface in the CypD structure.

**Supplementary Figure 12.** Structure-based sequence alignment of CypD, EpiD and $TvaF_{S\text{-}87}$.

**Supplementary Figure 13.** Structural and biochemical analyses of the binding interface between two trimeric units of the $TvaF_{S\text{-}87}$ dodecamer.

**Supplementary Figure 14.** Structural analysis of the binding interface between two trimeric subunits of the CypD dodecamer.

**Supplementary Figure 15.** Structural analysis of the CypD monomer in the CypD complex with the peptide substrate **8**.

**Supplementary Figure 16.** Structural analysis of the substrate-binding pockets of CypD dodecamer.

**Supplementary Figure 17.** Structural comparison of the CypD complex and the EpiD H67N mutant complex.

**Supplementary Figure 18.** *In vitro* assays of CypD activity using the substrates, **7**, **7-C19S** and **7-C19S-D$_3$**.

**Supplementary Figure 19.** Characterization of Dha-containing peptides in the CypD-catalyzed conversion of **7-Dha** by chemical derivation.

**Supplementary Figure 20.** Comparison of the enzymatic activities of CypD with its variants based on examination of the production of **7-I**, **7-II** and **7-III**.

**Supplementary Tables**

**Supplementary Table 1.** Related bacterial strains and plasmids used in this study.

**Supplementary Table 2.** Primers used in this study.

**Supplementary Table 3.** Gene sequences of *cypD*, $tvaF_{S\text{-}87}$, and *3c*.

**Supplementary Table 4.** HR-MS and HR-MS/MS Data collection of all compounds analyzed in this study.

**Supplementary Table 5.** The $^{1}$H and $^{13}$C NMR data of **7** in DMSO-$d_6$.

**Supplementary Table 6.** Key data in the 2D-TOCSY f2-slice at f1 experiments.

**Supplementary Table 7.** Statistics of X-ray crystallographic data collection and model refinements of $TvaF_{S\text{-}87}$ and CypD in their apo-forms.

**Supplementary Table 8.** Statistics of X-ray crystallographic data collection and model refinement of CypD in complex with **8**.

**Supplementary References**

**Supplementary Methods**

### 1.1 General Materials and Methods

**Materials, bacteria strains and plasmids.** Biochemicals and media were purchased from Sinopharm Chemical Reagent Co., Ltd. (China), Oxoid Ltd. (U.K.) or Sigma-Aldrich Corporation (USA) unless otherwise stated. Restriction endonucleases were purchased from Thermo Fisher Scientific Co. Ltd. (USA). Chemical reagents were purchased from standard commercial sources. Synthetic peptides were purchased from Genscript Biotech (Nanjing, China). Related bacterial strains and plasmids are summarized in **Supplementary Table 1**. Primers used in this study are listed in **Supplementary Table 2**.

**DNA isolation, manipulation, and sequencing.** DNA isolation and manipulation in *Escherichia coli* or *Streptomyces* strains were carried out according to standard methods[1]. PCR amplifications were carried out on an Applied Biosystems Veriti™ Thermal Cycler either using Taq DNA polymerase (Vazyme Biotech Co. Ltd, China) for routine genotype verification or PrimeSTAR HS DNA polymerase (Takara Biotechnology Co., Ltd. Japan) for high fidelity amplification. The synthesis of primers and genes were performed at Shanghai Sangon Biotech Co., Ltd. (China). DNA sequencing was performed at Shanghai Biosune Biotech Co., Ltd. (China).

**Sequence analysis.** Biosynthetic gene clusters (BGCc) were mined from microbial genomes using the AntiSMASH web tool[2]. Open reading frames (ORFs) were identified using the FramePlot 4.0beta program (http://nocardia.nih.go.jp/fp4/)[3]. The deduced proteins were compared with other known proteins in the databases using available BLAST methods (http://blast.ncbi.nlm.nih.gov/Blast.cgi)[4]. Amino acid sequence alignments were performed using Vector NT1 and ESPript 3.0 (http://espript.ibcp.fr/ESPript/ESPript/)[5].

**Chemical analysis.** Analysis and semi-preparation by High Performance Liquid Chromatography (HPLC) were carried out on an Agilent 1260 HPLC system (Agilent Technologies Inc., USA). Analyses by HPLC-associated Electrospray ionization Mass Spectrometer (ESI-MS) and ESI-high resolution MS (ESI-HR-MS) were performed on a Thermo Fisher LTQ XL ESI-MS spectrometer and a Q Exactive™ Plus Mass Spectrometer (Thermo Fisher Scientific Inc., USA), respectively. Related data were processed using Thermo Xcalibur software. NMR data were recorded on a Bruker AV500 spectrometers (Bruker Co. Ltd., Germany) or on an Agilent PremiumCompact+ 500MHz NMR spectrometer (Agilent Technologies Inc., USA).

#### 1.2 Protein Expression and Purification

**Construction and overexpression in *E. coli*.** The gene *tvaF*$_{S\text{-}87}$ was amplified from the genome of *S*. sp. NRRL S-87 by PCR using the primer pair *tvaF*$_{S\text{-}87}$-for/ *tvaF*$_{S\text{-}87}$ -rev, in contrast to the gene *cypD*, which was synthesized by Genscript Biotech (Nanjing, China). The gene *tvaF*$_{S\text{-}87}$ and *cypD* were cloned individually into pRSFDuet-1 for the expression of the recombinant proteins TvaF$_{S\text{-}87}$ and CypD, each of which is tagged by Thioredoxin (TRX) and 6xHis at N-terminus. To prepare the variants of TvaF$_{S\text{-}87}$, i.e., TvaF$_{S\text{-}87}$-H85A, TvaF$_{S\text{-}87}$ -V28D, and TvaF$_{S\text{-}87}$-M62D, and the variants of CypD, i.e., CypD-S20A, CypD-S20D, CypD-L23A, CypD-L23Q, CypD-F170A, CypD-F170Q, CypD-H59D, CypD-H59A, CypD-H29R, CypD-H29A, CypD-N80H and CypD-N80D by Site-specific mutagenesis, the Rolling-cycle PCR amplification of each pRSFDuet-1 derivative that contains *tvaF*$_{S\text{-}87}$ or *cypD* was conducted by using corresponding primers, and subsequent *Dpn*I digestion was performed according to the standard procedure of the QuickChange Site-Directed Mutagenesis Kit purchased from Stratagene (GE Healthcare, USA) and Multi Express™ II (Vazyme Biotech Co., Ltd., China). For N-terminal tag removal, the protease-encoding gene *3c* was synthesized by Genscript Biotech (Nanjing, China), and cloned into pET28a(+) for the expression of the recombinant 3C protein.

The above derivatives of pRSFDuet-1 and pET28a(+) were introduced into *E. coli* BL21(DE3) individually. The culture of each resulting recombinant *E. coli* strain was incubated in Luria-Bertani (LB) medium (5 g of yeast extract, 10 g of tryptone and 10 g of NaCl per liter) containing 50 μg/mL kanamycin at 37°C and 250 rpm until the cell density reached 0.6-0.8 at $OD_{600}$. Protein expression was induced by the addition of isopropyl-β-D-thiogalactopyranoside (IPTG) to a final concentration of 0.1 mM, followed by further incubation for 25-30 hr at 25°C or 16°C. The cells were harvested by centrifugation at 3000 × g for 20 min, flash-frozen and then stored at -80°C.

**Purification and characterization.** *E. coli* cells were re-suspended in lysis buffer (137 mM NaCl, 2.7 mM KCl, 10 mM $Na_2HPO_4$, 1.8 mM $KH_2PO_4$, 10% glycerol and 5 mM imidazole, pH 8.0). After disruption by FB-110X Low Temperature Ultra-Pressure Continuous Flow Cell Disrupter (Shanghai Litu Mechanical Equipment Engineering Co., Ltd, China), soluble fractions were collected by centrifugation. Recombinant proteins that contain a 6xHis-tag were purified on a HisTrap HP column (GE Healthcare, USA), which was pre-treated with 10 column volumes (CVs) of lysis buffer followed by 10 CVs of wash buffer (137 mM NaCl, 2.7 mM KCl, 10 mM $Na_2HPO_4$, 1.8 mM $KH_2PO_4$, 10% glycerol and 40 mM imidazole, pH 7.4), using elution buffer (137 mM NaCl, 2.7 mM KCl, 10 mM $Na_2HPO_4$, 1.8 mM $KH_2PO_4$, 10% glycerol and 250 mM imidazole, pH 7.4). Desired protein fractions were concentrated (to 500 μM-1 mM) using Amicon® Ultra-15 Centrifugal Filter Devices (MILLIPORE, USA) and desalted using a PD-10 Desalting Column (GE Healthcare, USA) according to the manufacturer's protocols, and then quantified in concentration by Bradford assay using bovine serum albumin as the standard.

The purity of recombinant proteins was determined by sodium dodecyl sulfate polyacrylamide gel electrophoresis (SDS-PAGE). For the determination of the flavin cofactor associated with $TvaF_{S-87}$ and CypD, The UV spectra of recombinant proteins were recorded at a concentration of 30 mg/mL on a DeNovix DS-11 UV/Vis spectrophotometer (DeNovix Inc. Wilmington, DE 19810 USA). Each protein solution was incubated at 100˚C for 5 min for denaturation and then subjected to HR-ESI-MS analysis on a 6230B Accurate Mass TOF LC/MS System (Agilent Technologies Inc., USA) to examine the presence of FMN ($[M+H]^+$ *m/z* calcd. 457.1124; observed 457.1108 from the $TvaF_{S-87}$) and the presence of FAD ($[M+H]^+$ *m/z* calcd. 786.1644; observed 786.1647 from the CypD sample).

#### 1.3 *In vitro* Assays of LanD-like Flavoprotein Activity

**Transformations of synthetic peptides.** Each conversion was conducted at 30°C for 2 or 5 hr in 60 µL of the reaction mixture that contained 100 µM (or 30 µM) synthetic peptide and 30 µM (or 10 µM) N-terminally TRX-tagged $TvaF_{S-87}$ (or its variant) or CypD (or its variant) along with 50 mM Tris-HCl (pH 8.5), 10 mM TCEP, 5 µM FMN or FAD and 1 µM 3C. Conversions were quenched by adding equal volumes of acetonitrile, and after centrifugation, reaction mixtures were subjected to HPLC-HR-MS and HR-MS/MS analyses. For (ene)thiol determination, 16 µL of each quenched reaction mixture was treated with *N*-ethylmaleimide (NEM) in dark at room temperature for 30 min before 2 µL of 1 M dithiothreitol (DTT) was added to eliminate excessive NEM and prevent side reactions. For aldehyde determination, 16 µL of each quenched reaction mixture was treated with 200 mM 1-(2-hydrazinyl-2-oxoethyl) pyridin-1-ium chloride (HOPI) in dark at 37°C for 2 hr.

**HPLC-HR-MS and HR-MS/MS analyses**. Reaction mixtures were subjected to HPLC-HR-MS and HR-MS/MS analyses after centrifugation. For HPLC-HR-MS analysis, an Agilent ZORBAX column (300SB-C18, 2.1 mm × 100 mm, 3.5 µm, Agilent Technologies Inc., USA) or an Agilent 300Extend-C18 column (2.1 mm × 100 mm, 3.5 µm, Agilent Technologies Inc., USA) was used on an UltiMate 3000 UHPLC system coupled to a Thermo Scientific Q Exactive Plus Orbitrap mass spectrometer. Gradient elusion was conducted using solvent A ($H_2O$ + 0.1% formic acid) and solvent B (acetonitrile + 0.1% formic acid) with a flow rate of 0.3 mL/min over a 43 min period as follows: T = 0 min, 10% B; T = 2 min, 10% B; T = 20 min, 30% B; T = 25 min, 60% B; T = 30 min, 100% B; T = 35 min, 100% B; T = 38 min, 10% B; and T = 43 min, 10% B. Unless otherwise stated, ESI-MS was performed in positive ion mode, with a spray voltage of 3800 V, a capillary temperature of 375 °C, aux gas heater temperature 350 °C and an S-lens level 60. Full MS was examined at a resolution of 70,000 (AGC target 2e5, maximum IT 50 ms, range 300–1000 or 400-1800 m/z).

Parallel reaction monitoring (PRM) or data-dependent $MS^2$ was performed at a resolution of 35,000 (AGC target between 1e5 and 1e6, maximum IT between 100 ms and 250 ms, isolation windows in the range of 1.0 to 2.0 m/z) using a stepped NCE of 18, 20 and 28 or an NCE of 25. Scan ranges, inclusion lists, charge exclusions, and dynamic exclusions were adjusted as needed.

#### 1.4 Peptide Purification and Characterization

**Production and isolation of 7-III.** The *in vitro* enzymatic transformation of **7** into **7-III** was carried out as described above. Each 500 µL of the reaction mixture was then applied to a Thermo C18 HyperSep cartridge coupled with a Vacuum Extraction Manifold system (Agilent Technologies Inc., USA). This cartridge was equilibrated with ACN and the ACN solution (5%) containing 0.1% formic acid. After washing with the same ACN solution to remove polar contaminants and elution with the ACN solution (40%) containing 0.1% formic acid, the collection was freeze-dried using a freeze dryer (Martin Christ Inc., Germany). Further purification was conducted by RP-HPLC on a Agilent ZORBAX column (250 × 4.6 mm, 5 µm, Agilent Technologies Inc., USA) by gradient elution of solvent A ($H_2O$ + 0.01% formic acid) and solvent B (acetonitrile + 0.01% formic acid) with a flow rate of 1 mL/min over a 35 min period as follows: T = 0 min, 15% B; T = 3 min, 15% B; T = 20 min, 60% B; T = 25 min,100% B; T = 30 min, 100% B; and T = 35 min, 15% B ($\lambda$ at 202 nm). After lyophilization, the collection was subject to HPLC-HR-MS and HR-MS/MS analyses under conditions as described above.

**Structural characterization of 7-III.** For comparison, the synthetic peptide substrate **7** (Genscript Biotech, Nanjing, China) was subjected to spectral analysis. **7** was obtained as a white amorphous powder: HR-ESI-MS (positive mode) *m/z* $[M+H]^+$, calcd. 795.3739 for $C_{32}H_{58}N_8O_{11}S_2$, obs. 795.3726; $^1H$ and $^{13}C$ NMR data (600 and 150 MHz, respectively, recorded in DMSO-$d_6$), see **Supplementary Table 5**. Clearly, the structure of **7** is characterized as the sequence **G**1-**S**2-**T**3-**I**4-**C**5-**L**6-**V**7-**C**8.

HR-ESI-MS established the molecular formula of **7-III** as $C_{31}H_{54}N_8O_9S$ (*m/z* $[M+H]^+$, calcd. 715.3808, obs. 715.3805). $^1H$ NMR data of **7-III** were recorded in DMSO-$d_6$. For structural comparison, the slice spectra from 2D TOCSY were collected using a previously established TOCSY method[6-12], to identify the six residues (i.e., **G**1, **S**2, **T**3, **I**4, **L**6 and **V**7) that are identical to those in **7** and the newly formed AviCys residue that is derived from the two residues **C**5 and **C**8 of **7**. The key signals of each residue (i.e., **S**2, **T**3, **I**4, **L**6, **V**7 or AviCys) were observed according to the 2D-TOCSY f2-slice at the related f1 experiment (S**upplementary Figures 8** and **Supplementary Table 6**). Specifically, a pair of olefin protons were observed by irritating –NH (H) of the 2-aminovinyl group of the AviCys residue. Together with $^3J_{H1/H2}$ = 7.3 Hz ($^3J_{H2/H1}$ = 7.3 Hz), the existence of the vinyl group with a *Z* configuration was indicated by the topspin software (https://www.bruker.com/service/support-upgrades/).

#### 1.5 Protein Crystallization and Structural Elucidation

**Size exclusion chromatography coupled with multi-angle light scattering (SEC-MALS).** SEC-MALS experiments were performed on a dynamic light scattering detector (DynaPro NanoStar, Wyatt), a static light scattering detector (DAWN HELLOS-II, Wyatt), and a differential refractive index detector (Optilab T-rEX, Wyatt) coupled with an AKTA pure system (GE Healthcare) at 25°C. All the protein samples (concentration above 3 mg/ml) were filtered and loaded into a Superdex 200 Increase 10/300 GL column pre-equilibrated by the buffer containing 20 mM Tris-HCl (pH 7.5), 100 mM NaCl, and 1 mM DTT overnight. The data was analyzed by ASTRA software (version 7.1) and further output to the Origin 9.0 software and aligned with each other.

**Protein crystallization and structural elucidation.** Crystals of seleno-methionine labeled $TvaF_{S-87}$ and CypD, CypD fusion proteins were obtained using the sitting-drop vapor-diffusion method at 16°C. Specifically, the freshly purified seleno-methionine labeled $TvaF_{S-87}$ protein (20 or 10 mg/ml in 20 mM Tris-HCl, 100 mM NaCl, 1 mM DTT, 1 mM EDTA at pH 7.5) was mixed with equal volume of reservoir solution containing 0.1 M amino acids (0.2 M L-Na-Glutamate, 0.2 M Alanine (racemic), 0.2 M Glycine, 0.2 M Lysine HCl (racemic), 0.2 M Serine (racemic)); 0.1 M buffer system at pH 8.5 (Tris (base), Bicine); 50% precipitant mix 3 (40% Glycerol, 20% PEG4000). While, crystals of seleno-methionine labeled CypD and CypD fusion protein (20 or 10 mg/ml in 20 mM Tris-HCl, 100 mM NaCl, 1 mM DTT, 1 mM EDTA at pH 7.5) were grown from 1.8 M Ammonium sulfate, 0.1 M BIS-TRIS pH 6.5, 2% v/v Polyethylene glycol monomethyl ether 550, and 0.15 M DL-Malic acid at pH 7.0, 0.1 M Imidazole at pH 7.0, 22% v/v Polyethylene glycol monomethyl ether 550, respectively. Before diffraction experiments, appropriate glycerol with FAD was added as the cryo-protectant. X-ray data sets were collected at the beamline BL17U1 or BL19U1 of the Shanghai Synchrotron Radiation Facility[13]. The diffraction data were processed and scaled using HKL2000[14].

The phase problems of $TvaF_{S-87}$ and CypD were solved by single wavelength anomalous diffraction (SAD) method. The phase problem of CypD fusion protein was solved by the molecular replacement method using the structure of CypD with PHASER[15]. The initial structural model was rebuilt manually using COOT[16], and then refined using REFMAC[17] or PHENIX[18]. Further manual model building and adjustment were completed using COOT[16]. The qualities of the final model were validated by MolProbity[19]. The final refinement statistics of solved structures in this study were listed in **Supplementary Tables 7 and 8**. All the structural diagrams were prepared using the program PyMOL (http://www.pymol.org/). Atomic coordinates and structural factors for the reported crystal structures of $TvaF_{S-87}$ and CypD are deposited in the Protein Data Bank under the accession numbers 6KTP, 6KTT, 6KTI, and 6KT9.

### 1.6 Chemical synthesis of 7-Dha19

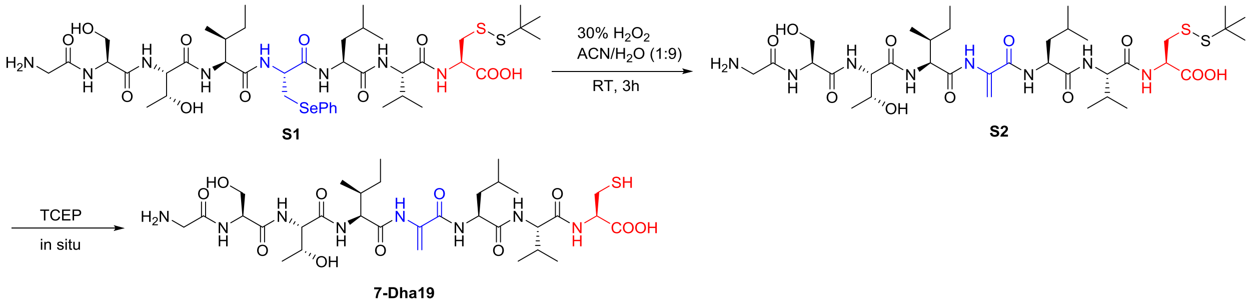


The dehydration of the synthetic substrate **S1** (Genscript Biotech, Nanjing, China) to the intermediate **S2** before *in situ* deprotection with TCEP was conducted in a reaction tube containing 100 µM $H_2O_2$ (30%, wt/wt) and 25µM **S1** by stirring at room temperature for 3 hr. The reaction mixture was quenched by addition of 10 µl dimethylsulfide. The quenched mixture was freeze-dried and then examined by HR-ESI-MS on a 6230B Accurate Mass TOF LC/MS System (Agilent Technologies Inc., USA) for the production of **S2** (*m/z* $[M+H]^+$ calcd. 849.4209; obs. 849.4214).

### Supplementary Figures

**Supplementary Figure 1.** Alignment of the biosynthetic gene clusters of TVAs, CYP and EPI. Genes coding for precursor peptides and LanD-type flavoproteins are highlighted in blue and red, respectively. Overall, only genes encoding LanD-type flavoproteins are conserved among the *cyp*, *epi* and *tva* clusters. $TvaF_{S-87}$ shares 21.8% and 18.9% with CypD and the archetypal LanD protein EpiD, respectively, while CypD share 17.5% with EpiD. For details in sequence identity, see **Supplementary Figure 12**.

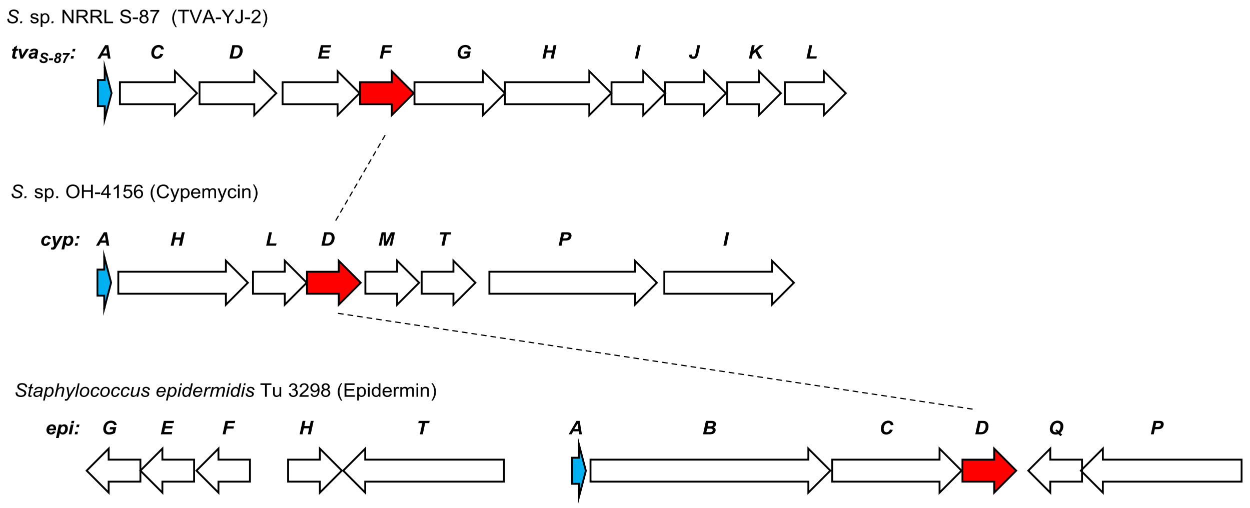

**Supplementary Figure 2.** Purified recombinant enzymes and cofactor analysis. **(a)** Coomassie-stained SDS-PAGE analysis of TRX-tagged $TvaF_{S\text{-}87}$, its variants and 3C with protein standard, TRX-tagged CypD and its variants with protein standard. **(b)** UV-Vis spectra of TRX-tagged recombinant flavoproteins $TvaF_{S\text{-}87}$ and CypD. **(c)** Determination of flavin cofactors associated with $TvaF_{S\text{-}87}$ and CypD. (i) Authentic FAD; (ii) boiled CypD, (iii) authentic FMN, and (iv) boiled $TvaF_{S\text{-}87}$. For examination by HPLC, UV absorbance at 375 nm. **(d)** Multiangle light scattering (MALS) analysis of $TvaF_{S\text{-}87}$ and CypD, showing the relative light scattering signal as a function of elution volume. The derived molecular mass of the peak is shown in blue.

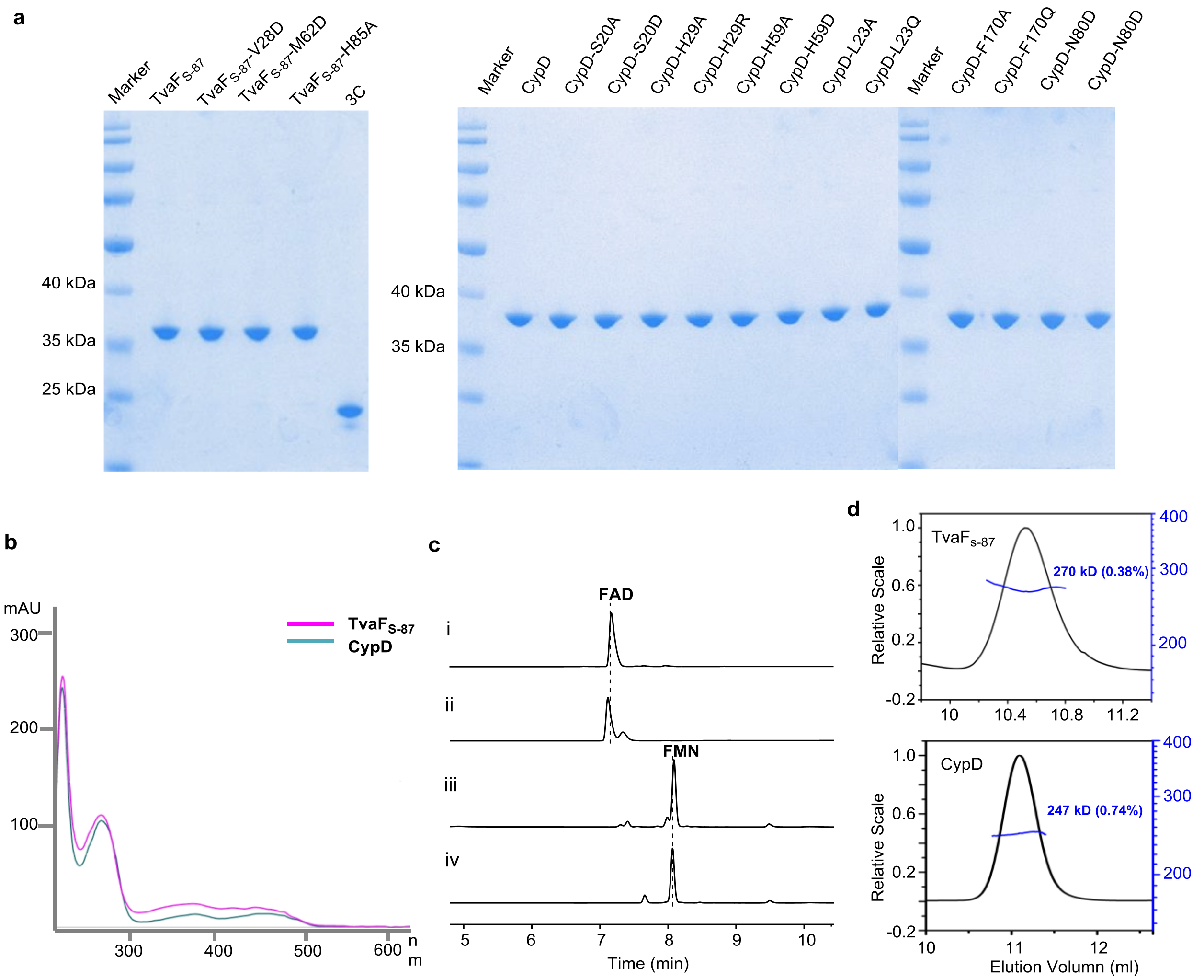

**Supplementary Figure 3.** Summary of LanD-like flavoprotein-catalyzed transformations in this study. Amino acids corresponding to the two target residues that are processed for Avi(Me)Cys formation and their related structures during each transformation process are colored, e.g., corresponding to the C-terminal L-Cys, red; and corresponding the internal L-Thr/L-Ser or L-Cys, blue. √, observed; ×, not observed; and **N.D.**, not detected. (**a**) TvaF$_{S\text{-}87}$. (**b**) CypD. For variation of the internal residue: $X_1$, L-[2,3,3-$D_3$]Ser; $X_2$, acetylated L-Ser; $X_3$, glutamylated L-Ser; $X_4$, phosphorylated L-Ser; $X_5$, Dha; and $X_6$, D-Cys.

| | Peptidyl Substrate | 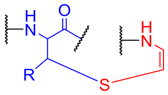 Cyclized Product | 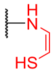 Enethiol Intermediate | 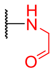 Aldehyde Shunt Product |
|---|---|---|---|---|
| **a** | | | | |
| **1** | SPDEEAQGSVMAAAATVAFHC | ✓ | ✓ | ✓ |
| **1-T8A** | SPDEEAQGSVMAAAAAVAFHC | ✗ | ✓ | ✓ |
| **1-T8S** | SPDEEAQGSVMAAAASVAFHC | ✓ | ✓ | ✓ |
| **1-T8C** | SPDEEAQGSVMAAAACVAFHC | ✗ | ✓ | ✓ |
| **1-C13A** | SPDEEAQGSVMAAAATVAFHA | ✗ | ✗ | ✗ |
| **1-C13S** | SPDEEAQGSVMAAAATVAFHS | ✗ | ✗ | ✗ |
| **1-C13T** | SPDEEAQGSVMAAAATVAFHT | ✗ | ✗ | ✗ |
| **2** | VMAAAATVAFHC | ✓ | ✓ | ✓ |
| **3** | MAAAATVAFHC | ✓ | ✓ | ✓ |
| **4** | AAAATVAFHC | ✗ | ✓ | **N.D.** |
| **5** | AATVAFHC | ✗ | ✓ | **N.D.** |
| **6** | TVAFHC | ✗ | ✓ | **N.D.** |
| **b** | | | | |
| **7** | GSTI CLVC | ✓ | ✓ | ✓ |
| **7-C19A** | GSTI ALVC | ✗ | ✓ | ✓ |
| **7-C19S** | GSTI SLVC | ✓ | ✓ | ✓ |
| **7-C19T** | GSTI TLVC | ✗ | ✓ | ✓ |
| **7-C22A** | GSTI CLVA | ✗ | ✗ | ✗ |
| **7-C22S** | GSTI CLVS | ✗ | ✗ | ✗ |
| **7-C22T** | GSTI CLVT | ✗ | ✗ | ✗ |
| **7-C19S-D$_3$** | GSTI $X_1$LVC | ✓ | ✓ | ✓ |
| **7-C19S-Ac** | GSTI $X_2$LVC | ✓ | ✓ | ✓ |
| **7-C19S-Glu** | GSTI $X_3$LVC | ✗ | ✗ | ✗ |
| **7-C19S-P** | GSTI $X_4$LVC | ✗ | ✗ | ✗ |
| **7-Dha19** | GSTI $X_5$LVC | ✓ | ✗ | ✗ |
| **7-d-C19** | GSTI $X_6$LVC | ✓ | ✓ | ✓ |

**Supplementary Figure 4.** HPLC-MS analysis of enethiol intermediates and associated shunt aldehyde derivatives in the TvaA$_{S-87}$-catalyzed conversions of **1** (the 21-aa C-terminal mimic of the precursor peptide TvaA$_{S-87}$) and its variants. Reactions were conducted at 30°C for 2 hr in the presence of TvaF$_{S-87}$ to convert the substrates **1** (i), **1-T8A** (ii), **1-T8S** (iii), **1-T8C** (iv) and **1-C13A** (v), and in the presence of TvaF$_{S-87}$-V28D (vi), TvaF$_{S-87}$-M62D (vii) and TvaF$_{S-87}$-H85A (viii) to convert **1**. (**b**) For detailed HR-MS and HR-MS/MS data, see **Supplementary Table 4**.

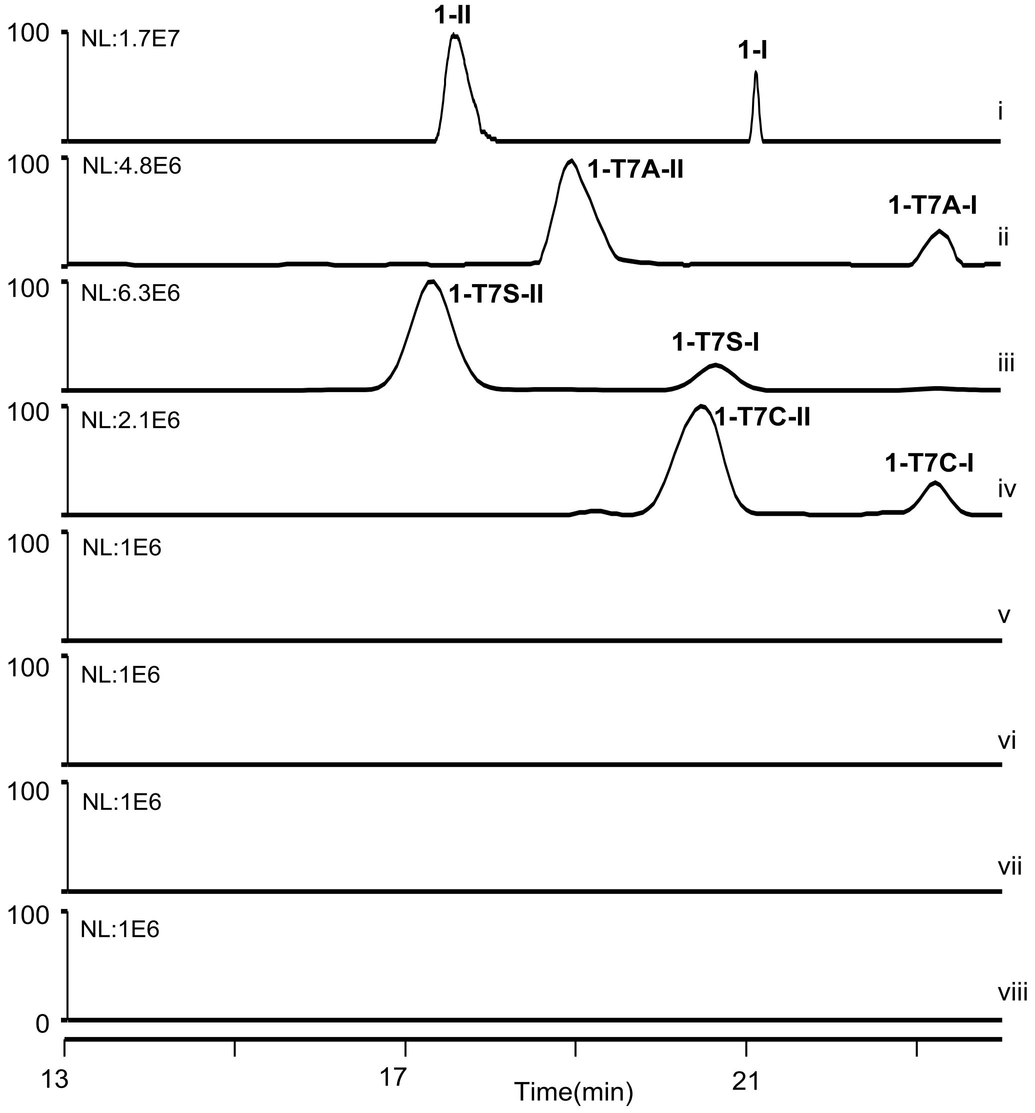

**Supplementary Figure 5.** Characterization of (ene)thiol and aldehyde peptides in the TvaF$_{S-87}$-catalyzed conversion of **1** by chemical derivatization. (**a**) Examination of **1**, **1-I** and **1-III** by HPLC-MS. i, standard **1**; ii, transformation of **1** at 30°C for 2 hr; iii, treatment of **1** with NEM; and iv, treatment of the reaction mixture of iii with NEM. (**b**) Examination of **1-II** by treating the TvaF$_{S-87}$-catalyzed transformation of **1** with 1-(2-Hydrazinyl-2- oxoethyl)pyridin-1-ium chloride (HOPI). (**c**) HR-MS and HR-MS/MS data for the derivatives **1-NEM**, **1-I-NEM** and **1-II-HOPI** of **1**, **1-I** and **1-II**, respectively.

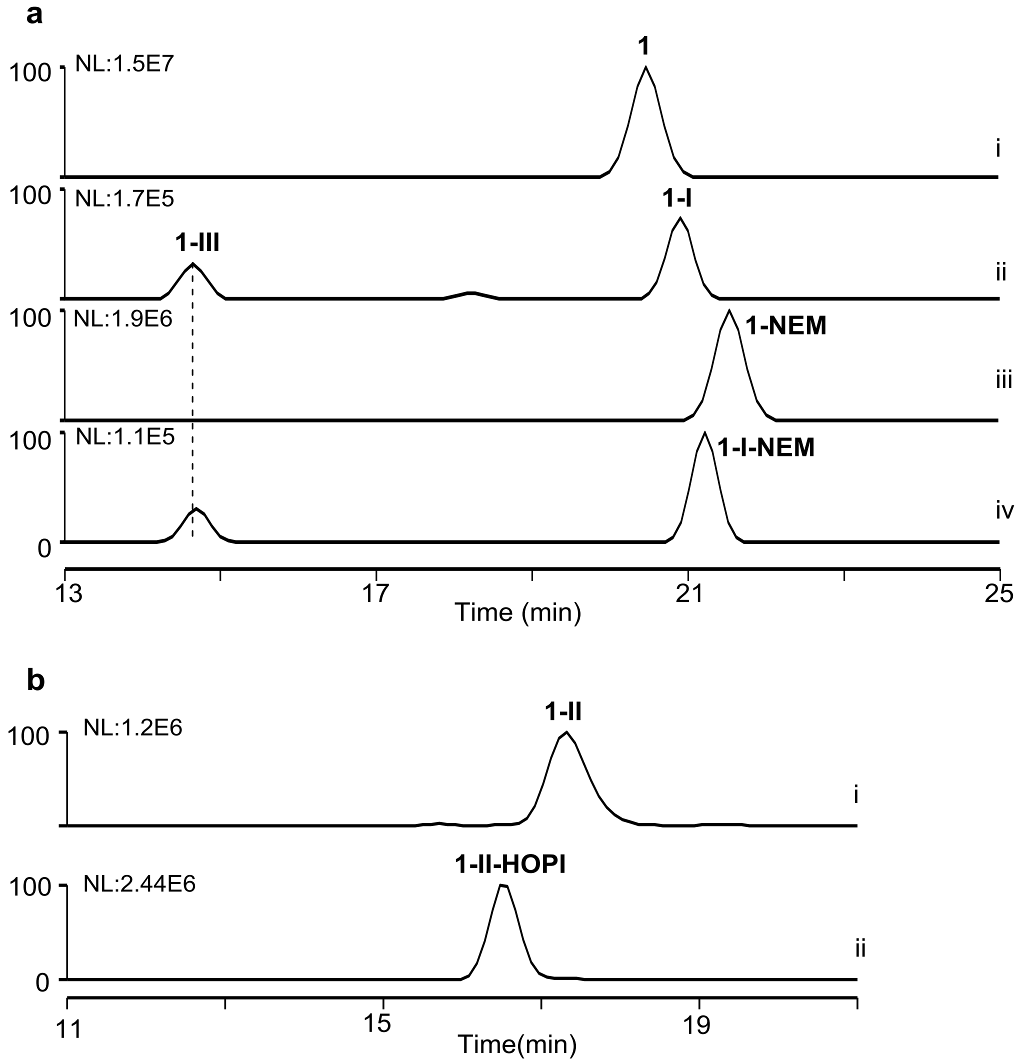

c

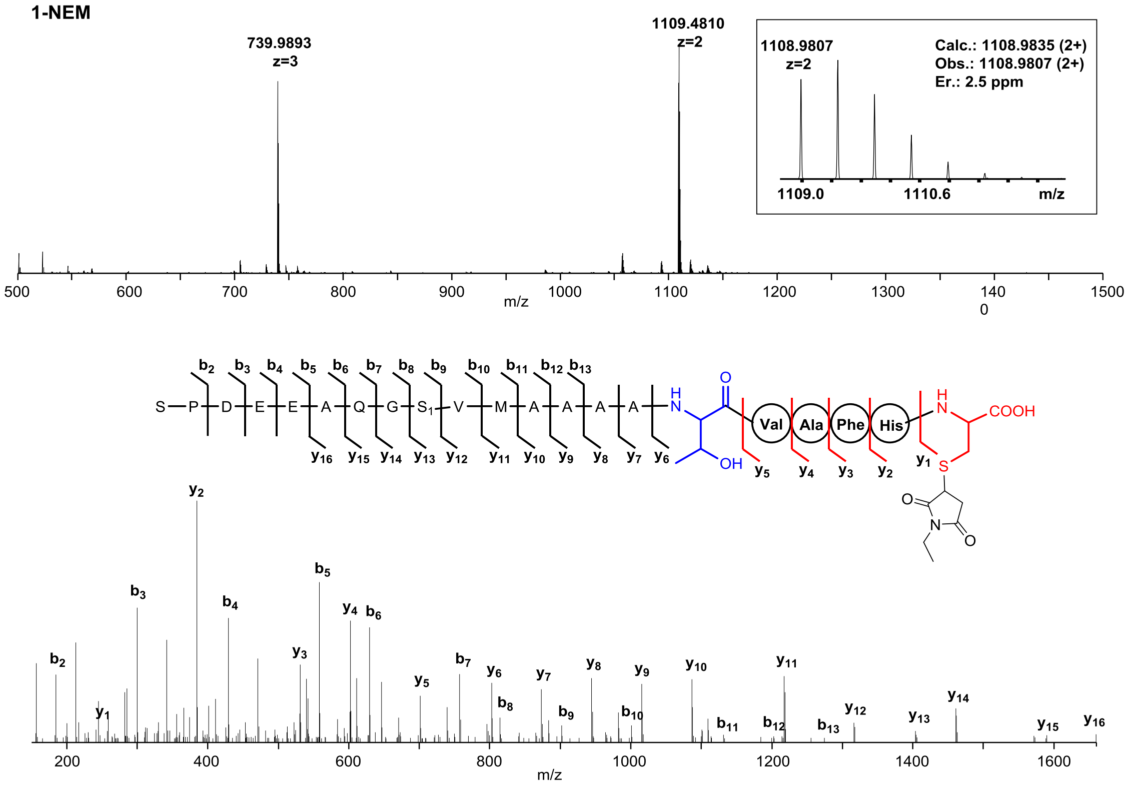


| Ions | Calcd. | Obs. | Er. (ppm) | Ions | Calcd. | Obs. | Er. (ppm) |
|---|---|---|---|---|---|---|---|
| $b_2$ | 185.0926 | 185.0917 | 4.9 | $y_1$ | 247.0752 | 247.0741 | 4.5 |
| $b_3$ | 300.1196 | 300.1183 | 4.3 | $y_2$ | 384.1341 | 384.1327 | 3.6 |
| $b_4$ | 429.1621 | 429.1608 | 3.0 | $y_3$ | 531.2025 | 531.2013 | 2.3 |
| $b_5$ | 558.2047 | 558.2032 | 2.7 | $y_4$ | 602.2396 | 602.2383 | 2.2 |
| $b_6$ | 629.2418 | 629.2400 | 2.9 | $y_5$ | 701.3081 | 701.3065 | 2.3 |
| $b_7$ | 757.3004 | 757.2977 | 3.6 | $y_6$ | 802.3557 | 802.3533 | 3.0 |
| $b_8$ | 814.3219 | 814.3192 | 3.3 | $y_7$ | 873.3928 | 873.3901 | 3.1 |
| $b_9$ | 901.3539 | 901.3534 | 0.6 | $y_8$ | 944.4300 | 944.4268 | 3.4 |
| $b_{10}$ | 1000.4223 | 1000.4199 | 2.4 | $y_9$ | 1015.4671 | 1015.4648 | 2.3 |
| $b_{11}$ | 1131.4628 | 1131.4600 | 2.5 | $y_{10}$ | 1086.5042 | 1086.5013 | 2.7 |
| $b_{12}$ | 1202.4999 | 1202.4928 | 5.9 | $y_{11}$ | 1217.5447 | 1217.5417 | 2.5 |
| $b_{13}$ | 1273.5371 | 1273.5386 | 1.2 | $y_{12}$ | 1316.6131 | 1316.6094 | 2.8 |
| | | | | $y_{13}$ | 1403.6451 | 1403.6471 | 1.4 |
| | | | | $y_{14}$ | 1460.6666 | 1460.6624 | 2.9 |

| | | | | | | | |
|---|---|---|---|---|---|---|---|
| | | | | $y_{15}$ | 1588.7252 | 1588.7162 | 5.7 |
| | | | | $y_{16}$ | 1659.7623 | 1659.7634 | 0.7 |

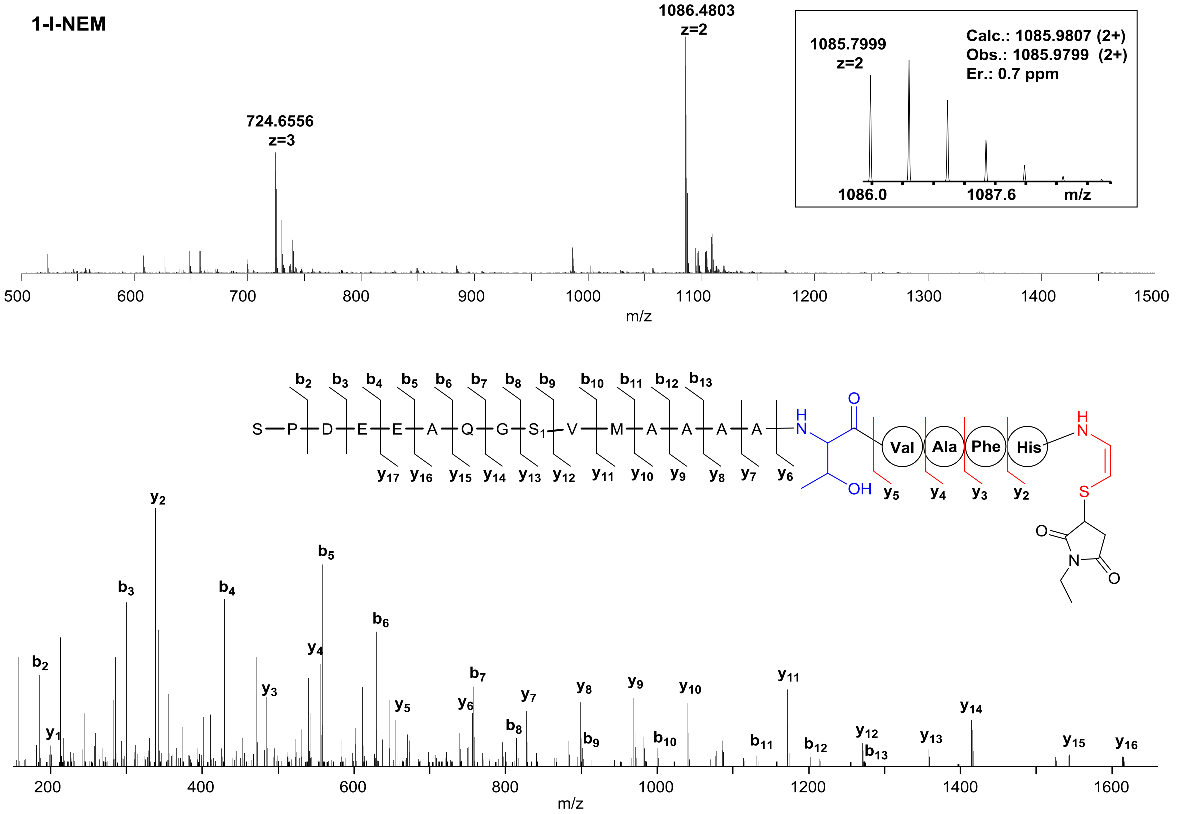


| Ions | Calcd. | Obs. | Er. (ppm) | Ions | Calcd. | Obs. | Er. (ppm) |
|---|---|---|---|---|---|---|---|
| $b_2$ | 185.0926 | 185.0919 | 3.8 | $y_1$ | 201.0697 | 201.0690 | 3.5 |
| $b_3$ | 300.1196 | 300.1186 | 3.3 | $y_2$ | 338.1286 | 338.1276 | 3.0 |
| $b_4$ | 429.1621 | 429.1611 | 2.3 | $y_3$ | 485.1970 | 485.1957 | 2.7 |
| $b_5$ | 558.2047 | 558.2033 | 2.5 | $y_4$ | 556.2342 | 556.2328 | 2.5 |
| $b_6$ | 629.2418 | 629.2401 | 2.7 | $y_5$ | 655.3026 | 655.3019 | 1.1 |
| $b_7$ | 757.3004 | 757.2966 | 5.0 | $y_6$ | 756.3503 | 756.3481 | 2.9 |
| $b_8$ | 814.3219 | 814.3212 | 0.9 | $y_7$ | 827.3874 | 827.3857 | 2.1 |
| $b_9$ | 901.3539 | 901.3505 | 3.8 | $y_8$ | 898.4245 | 898.4221 | 2.7 |
| $b_{10}$ | 1000.4223 | 1000.4212 | 1.1 | $y_9$ | 969.4616 | 969.4600 | 1.7 |
| $b_{11}$ | 1131.4628 | 1131.4642 | 1.2 | $y_{10}$ | 1040.4987 | 1040.4977 | 1.0 |
| $b_{12}$ | 1202.4999 | 1202.4954 | 3.7 | $y_{11}$ | 1171.5392 | 1171.5377 | 1.3 |

| | | | | | | | |
|---|---|---|---|---|---|---|---|
| $b_{13}$ | 1273.5371 | 1273.5284 | 6.8 | $y_{12}$ | 1270.6076 | 1270.6068 | 0.6 |
| | | | | $y_{13}$ | 1357.6396 | 1357.6368 | 2.1 |
| | | | | $y_{14}$ | 1414.6611 | 1414.6575 | 2.5 |
| | | | | $y_{15}$ | 1542.7197 | 1542.7157 | 2.6 |
| | | | | $y_{16}$ | 1613.7568 | 1613.7555 | 0.8 |

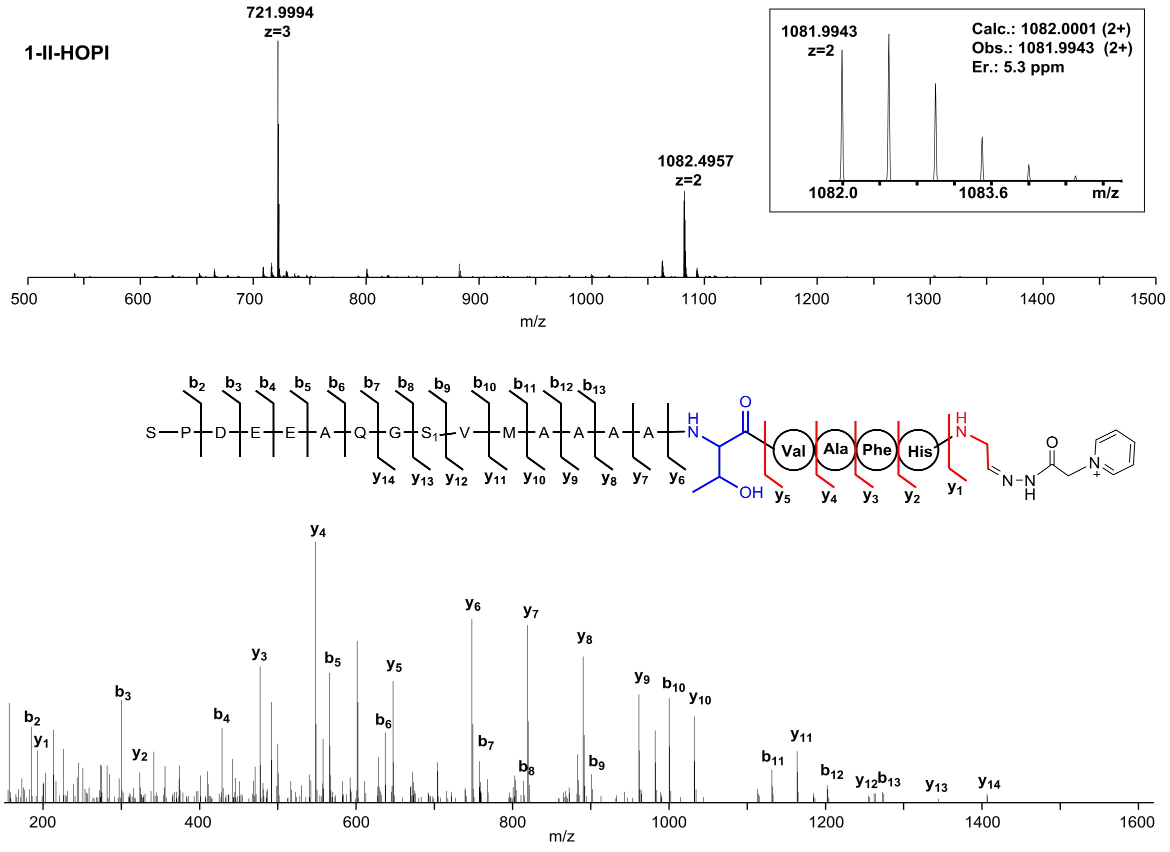


| Ions | Calcd. | Obs. | Er. (ppm) | Ions | Calcd. | Obs. | Er. (ppm) |
|---|---|---|---|---|---|---|---|
| $b_2$ | 185.0926 | 185.0915 | 5.9 | $y_1$ | 193.1084 | 193.1079 | 2.6 |
| $b_3$ | 300.1196 | 300.1182 | 4.7 | $y_2$ | 330.1673 | 330.1669 | 1.2 |
| $b_4$ | 429.1621 | 429.1604 | 3.9 | $y_3$ | 477.2357 | 477.2344 | 2.7 |
| $b_5$ | 558.2047 | 558.2009 | 6.8 | $y_4$ | 548.2728 | 548.2713 | 2.7 |
| $b_6$ | 629.2418 | 629.2360 | 9.2 | $y_5$ | 647.3412 | 647.3391 | 3.2 |
| $b_7$ | 757.3004 | 757.2967 | 4.9 | $y_6$ | 748.3889 | 748.3865 | 3.2 |
| $b_8$ | 814.3219 | 814.3193 | 3.2 | $y_7$ | 819.4260 | 819.4235 | 3.1 |

| | | | | | | | |
|---|---|---|---|---|---|---|---|
| $b_9$ | 901.3539 | 901.3504 | 3.8 | $y_8$ | 890.4631 | 890.4605 | 2.9 |
| $b_{10}$ | 1000.4223 | 1000.4188 | 3.5 | $y_9$ | 961.5002 | 961.4975 | 2.8 |
| $b_{11}$ | 1131.4628 | 1131.4590 | 3.4 | $y_{10}$ | 1032.5373 | 1032.5341 | 3.1 |
| $b_{12}$ | 1202.4999 | 1202.4945 | 4.5 | $y_{11}$ | 1163.5778 | 1163.5741 | 3.2 |
| $b_{13}$ | 1273.5371 | 1273.5334 | 2.9 | $y_{12}$ | 1262.6462 | 1262.6433 | 2.3 |
| | | | | $y_{13}$ | 1349.6783 | 1349.6676 | 7.9 |
| | | | | $y_{14}$ | 1406.6997 | 1406.6971 | 1.8 |

**Supplementary Figure 6.** HPLC-MS analysis of enethiol intermediates and associated shunt aldehyde derivatives in the CypD-catalyzed conversions of **7** (the 8-aa C-terminal mimic of the precursor peptide CypA) and its variants. (**a**) Conversions of **7** (i), **7-C19A** (ii), **7-C19T** (iii), **7-C19S** (iv), **7-C22A** (v) and **7-C22S** (vi). Reactions were conducted at 30°C for 2 hr in 60 μL of the reaction mixture that contained 100 μM synthetic peptide and 30 μM CypD. (**b**) Conversions of **7-Dha19** (i), **7** (ii), **7-d-C19** (iii), **7-C19S** (iv), **7-C19S-Ac** (v), **7-C19S-P** (vi) and **7-C19S-Glu** (vii). Reactions were conducted at 30°C for 2 hr in 60 μL of the reaction mixture that contained 30 μM synthetic peptide and 10 μM CypD. For detailed HR-MS and HR-MS/MS data, see **Supplementary Table 4**.

**a**

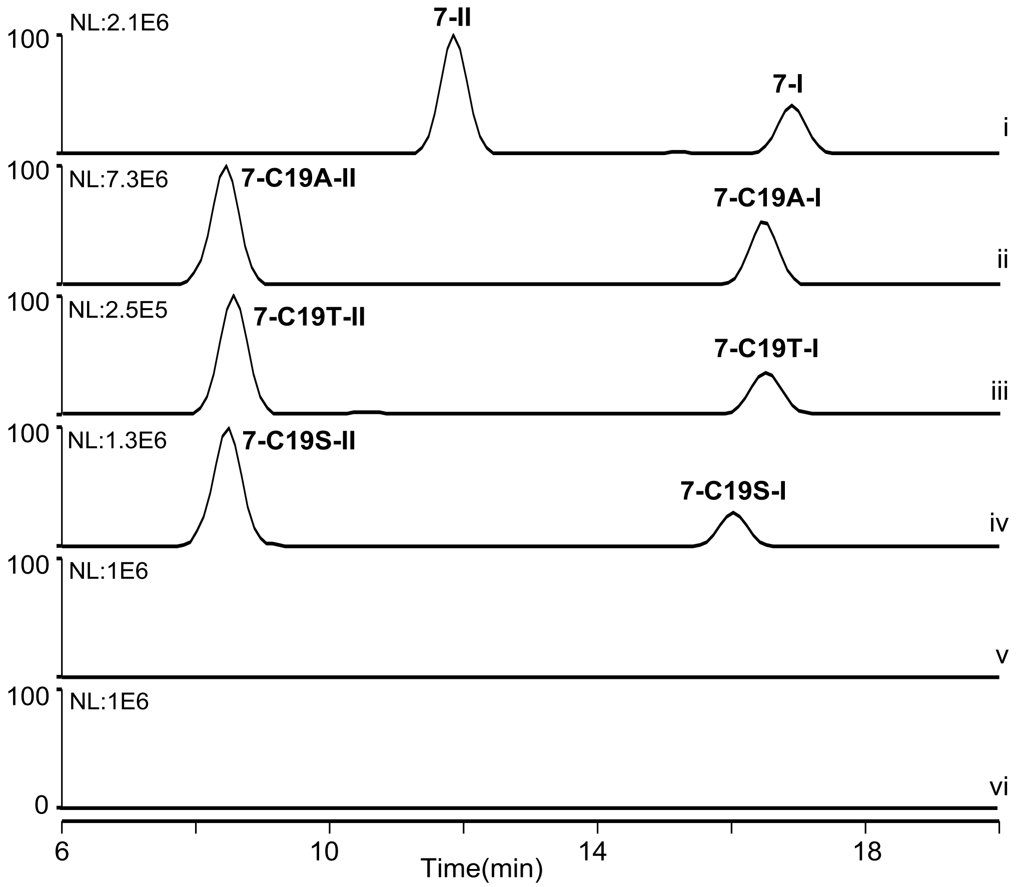

b

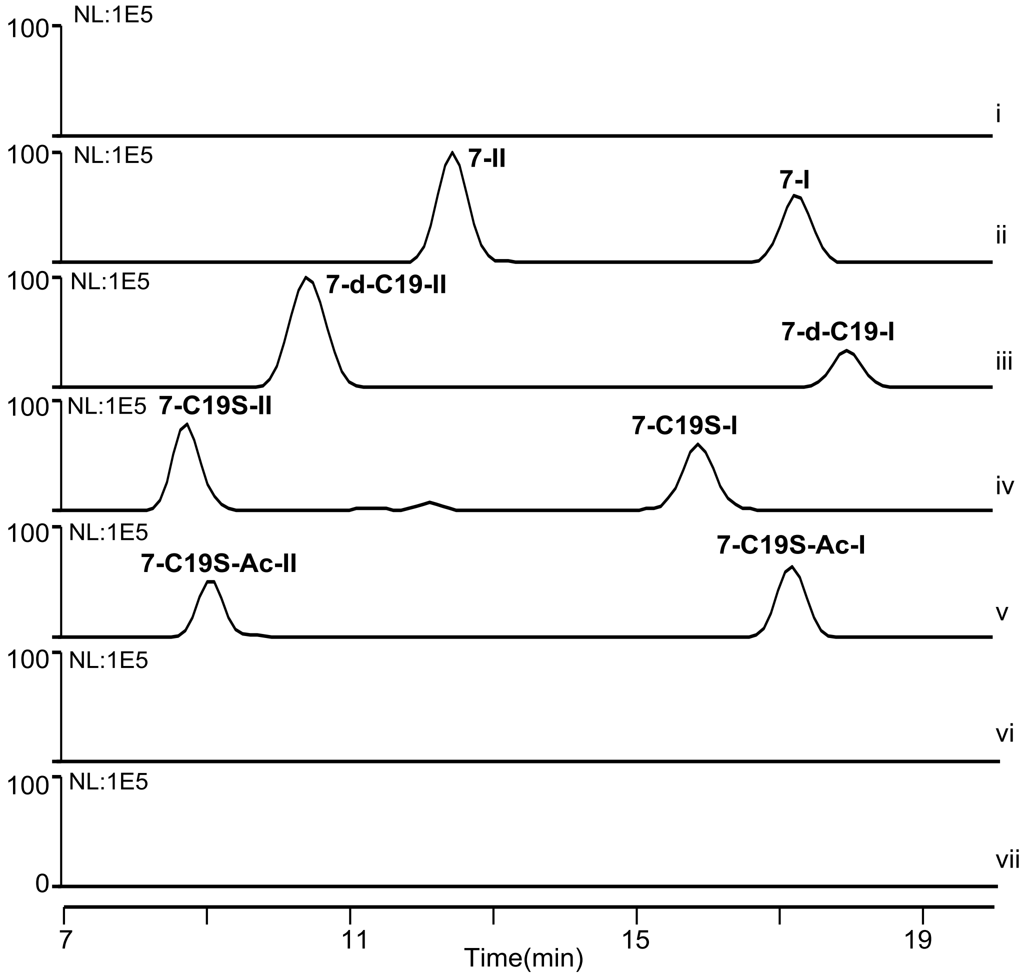

NL:1E5
100
i
NL:1E5
100
7-II
7-I
ii
100
NL:1E5
7-d-C19-II
7-d-C19-I
iii
100
NL:1E5
7-C19S-II
7-C19S-I
iv
100
NL:1E5
7-C19S-Ac-II
7-C19S-Ac-I
v
100
NL:1E5
vi
100
NL:1E5
vii
0
7
11
15
19
Time(min)

**Supplementary Figure 7.** Characterization of (ene)thiol peptides in the CypD-catalyzed conversion of **7** by chemical derivation. (**a**) Examination of **7**, **7-I** and **7-III** by HPLC-MS. i, standard **7**; ii, transformation of **7** into **7-I** and **7-III** at 30°C for 2 hr; iii, treatment of **7** with NEM; and iv, treatment of the reaction mixture of iii with NEM. (**c**) HR-MS and HR-MS/MS data for the derivatives **7-NEM** and **7-I-NEM** of **7** and **7-I**, respectively.

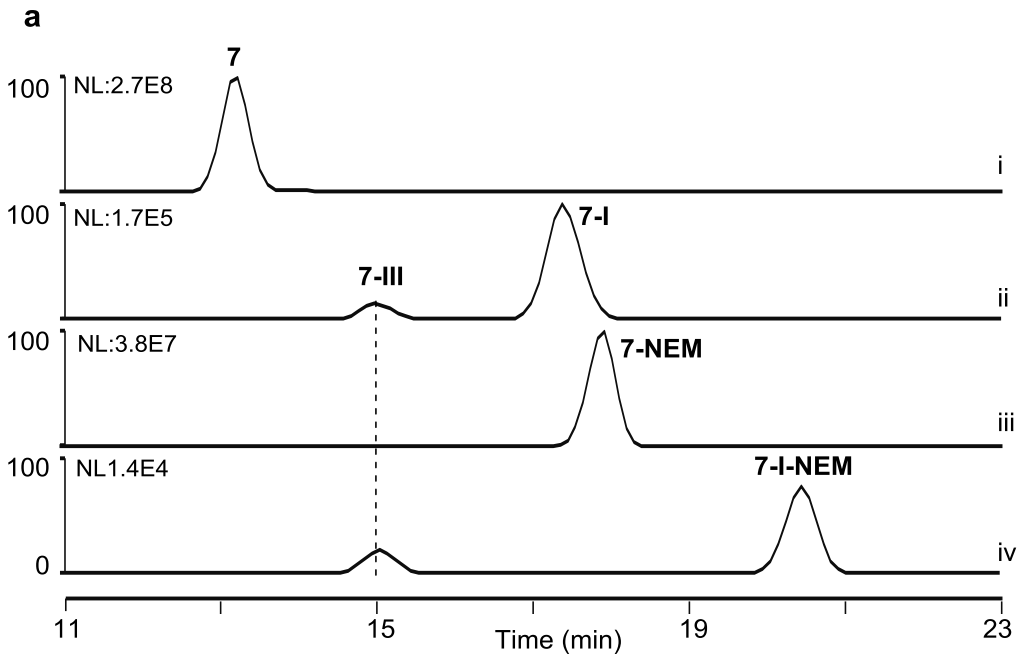

**b**

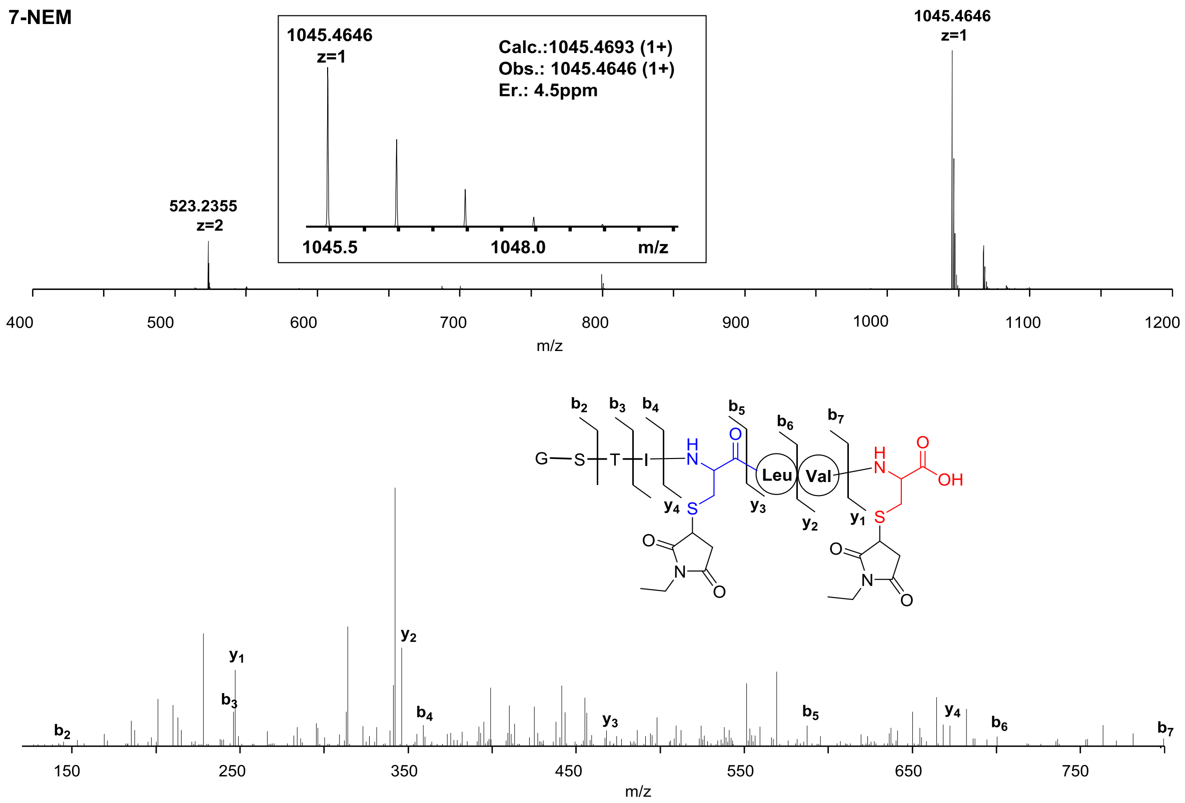


| Ions | Calcd. | Obs. | Er. (ppm) |
|---|---|---|---|
| $b_2$ | 145.0613 | 145.0605 | 5.5 |
| $b_3$ | 246.1090 | 246.1088 | 0.8 |
| $b_4$ | 359.1931 | 359.1942 | 3.1 |
| $b_5$ | 587.2499 | 587.2477 | 3.7 |
| $b_6$ | 700.3340 | 700.3318 | 3.1 |
| $b_7$ | 799.4024 | 799.3995 | 3.6 |
| $y_1$ | 247.0752 | 247.0741 | 4.5 |
| $y_2$ | 346.1436 | 346.1423 | 3.8 |
| $y_3$ | 459.2277 | 459.2276 | 0.2 |
| $y_4$ | 687.2845 | 687.2841 | 0.6 |

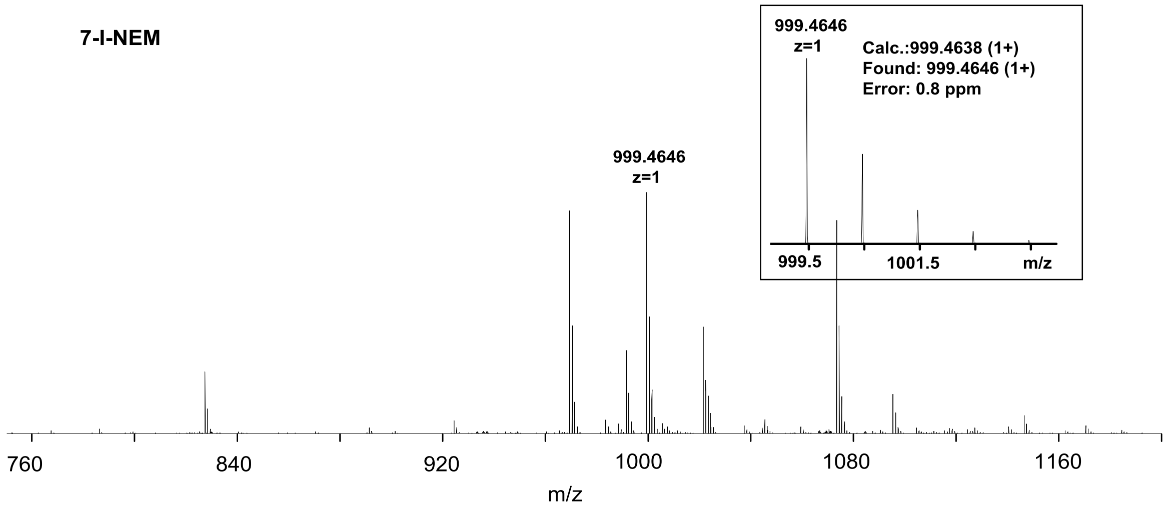


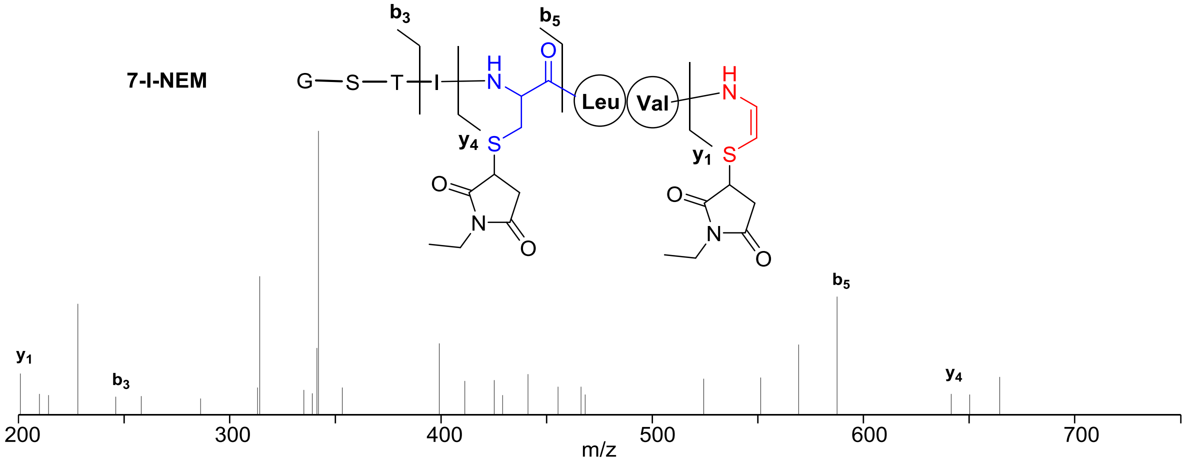


| Ions | Calcd. | Obs. | Er. (ppm) |
|---|---|---|---|
| $b_3$ | 246.1090 | 246.1070 | 8.1 |
| $b_5$ | 587.2499 | 587.2490 | 1.5 |
| $y_1$ | 201.0697 | 201.0686 | 5.4 |
| $y_4$ | 641.2791 | 641.2791 | 0.0 |

**Supplementary Figure 8.** 2D-TOCSY f2-slices at related f1 experiments (600 MHz, DMSO-$d_6$) for residue identification in **7-III**.

Chemical shift of -NH of the L-Ser residue (8.50 ppm).

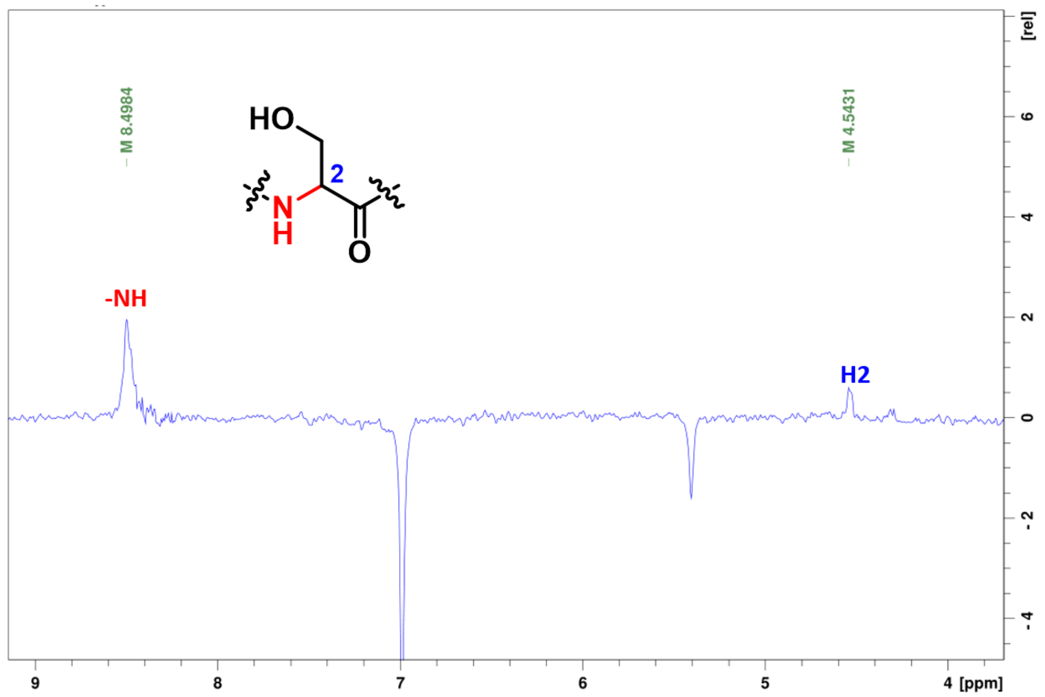


Chemical shift of -NH of L-Thr residue (8.04 ppm).

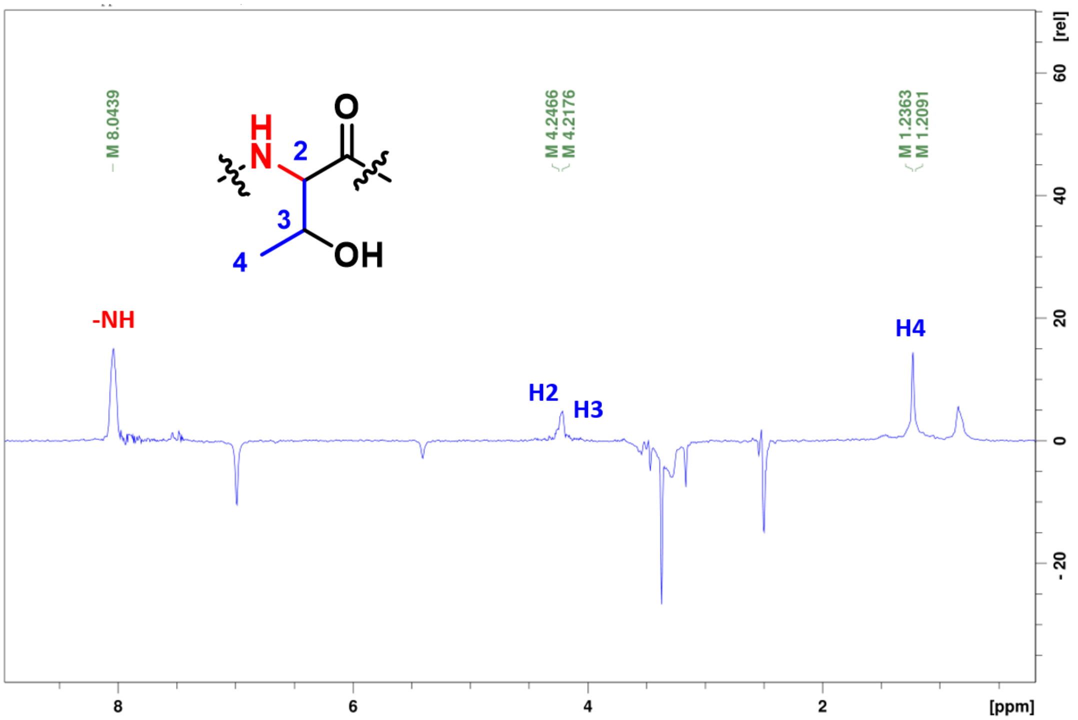

Chemical shift of H2 of L-Ile residue (4.10 ppm).

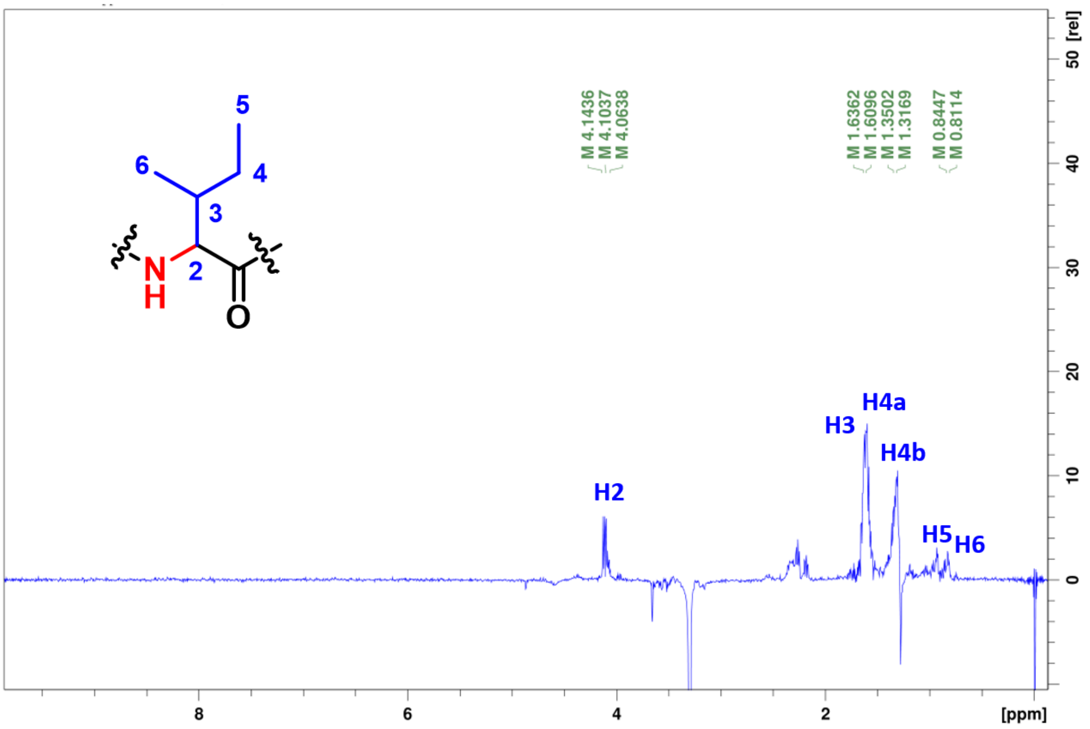


Chemical shift of NH of L-Leu residue (8.21 ppm).

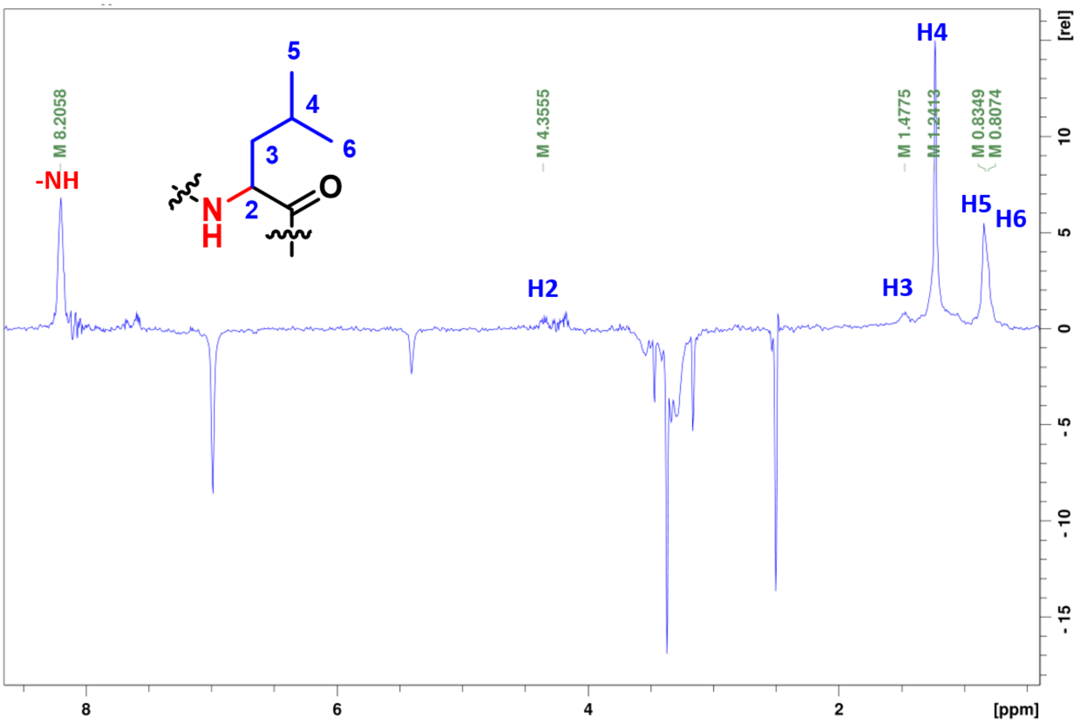


Chemical shift of NH of L-Val residue (7.00 ppm).

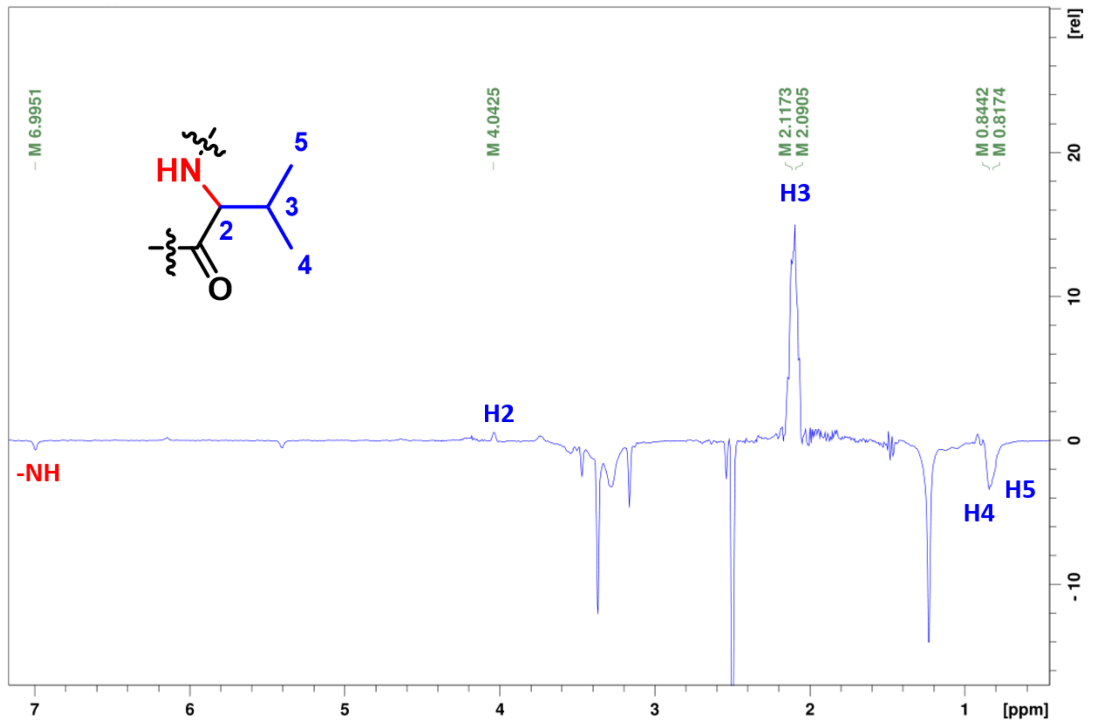


Chemical shift of NH of L-Cys2 residue (7.16 ppm).

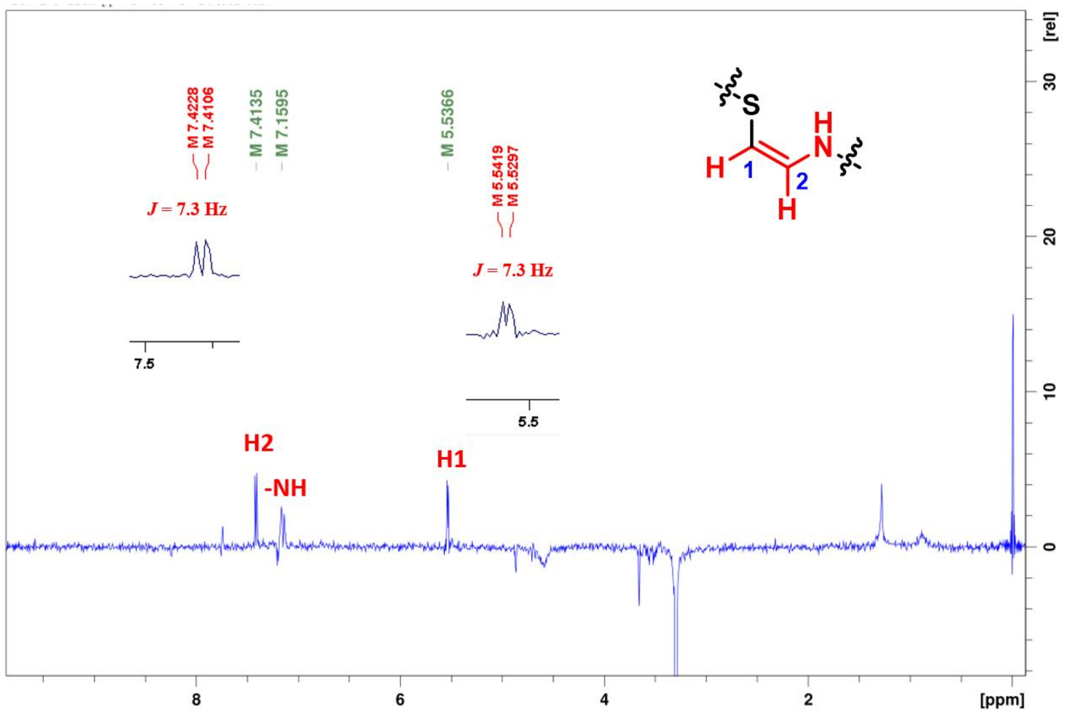

**Supplementary Figure 9.** Overall structural comparison of the monomers of TvaF$_{S-87}$, CypD, MrsD and the EpiD complex (shown by ribbon diagram). (**a**) TvaF$_{S-87}$ and CypD. (**b**) TvaF$_{S-87}$ EpiD. (**c**) CypD and EpiD. (**d**) TvaF$_{S-87}$ MrsD. (**e**) CypD and MrsD. (**f**) Two different TvaF$_{S-87}$ monomers (i.e., monomer 1 and monomer 2) found in TvaF$_{S-87}$ dodecamer. (**g**) The monomer of the EpiD H67N complex with a peptide substrate mimic (PDB ID: 1G5Q) and the TvaF$_{S-87}$ monomer 2.

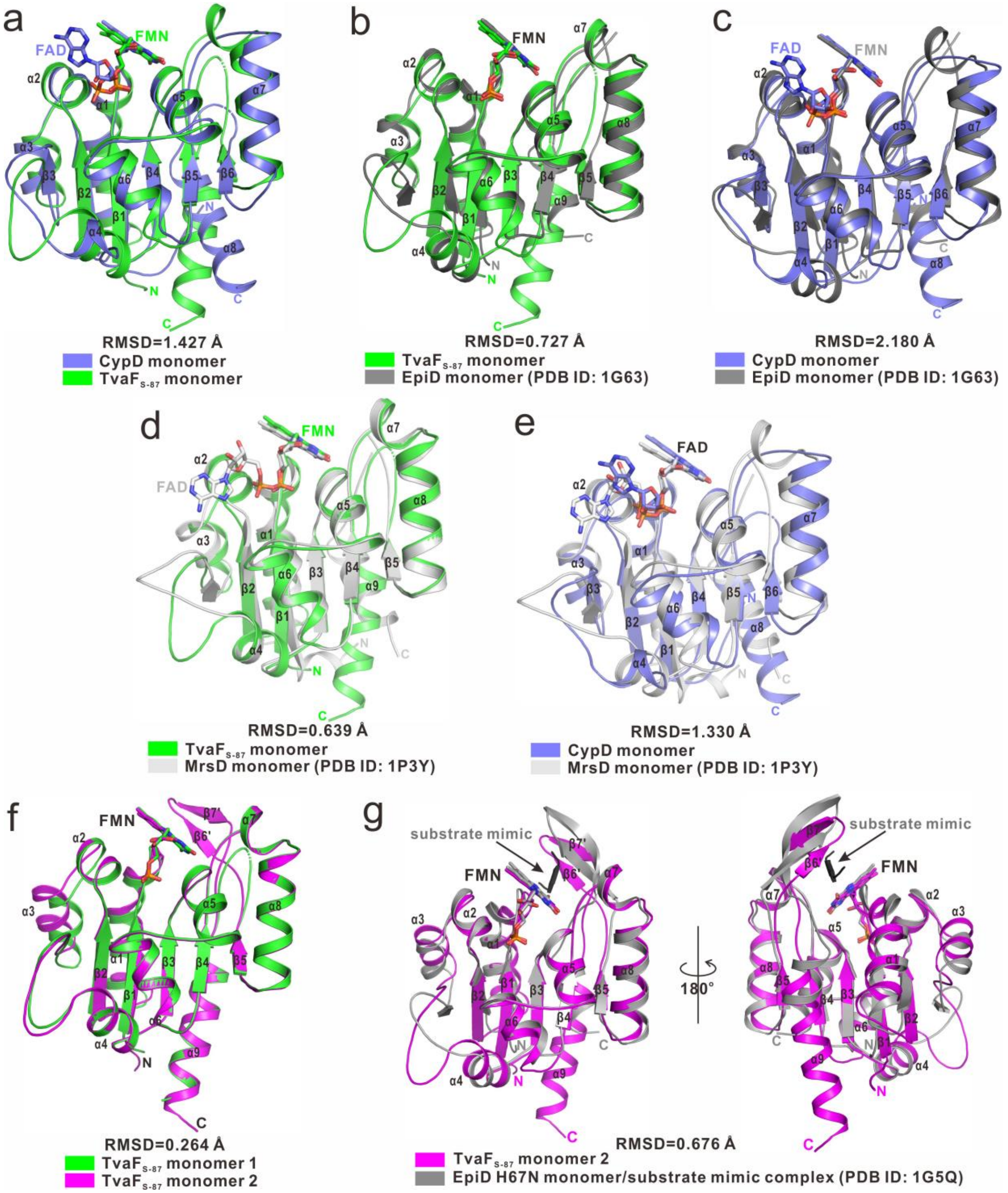

**Supplementary Figure 10.** FMN binding interface in the TvaF$_{S-87}$ structure. (**a**) The ribbon-stick model showing the FMN cofactors buried in the interfaces between two TvaF$_{S-87}$ monomers in a trimeric subunit. The three TvaF$_{S-87}$ monomers are shown in the ribbon model and colored in magenta, green, and orange, respectively, while the bound FMN cofactors are shown in the stick model. (**b**) The enlarged stereo view of the ribbon-stick-ball representation showing the detailed interactions between a FMN cofactor and two TvaF$_{S-87}$ monomers. The hydrogen bonds involved in the interaction are shown as dotted lines.

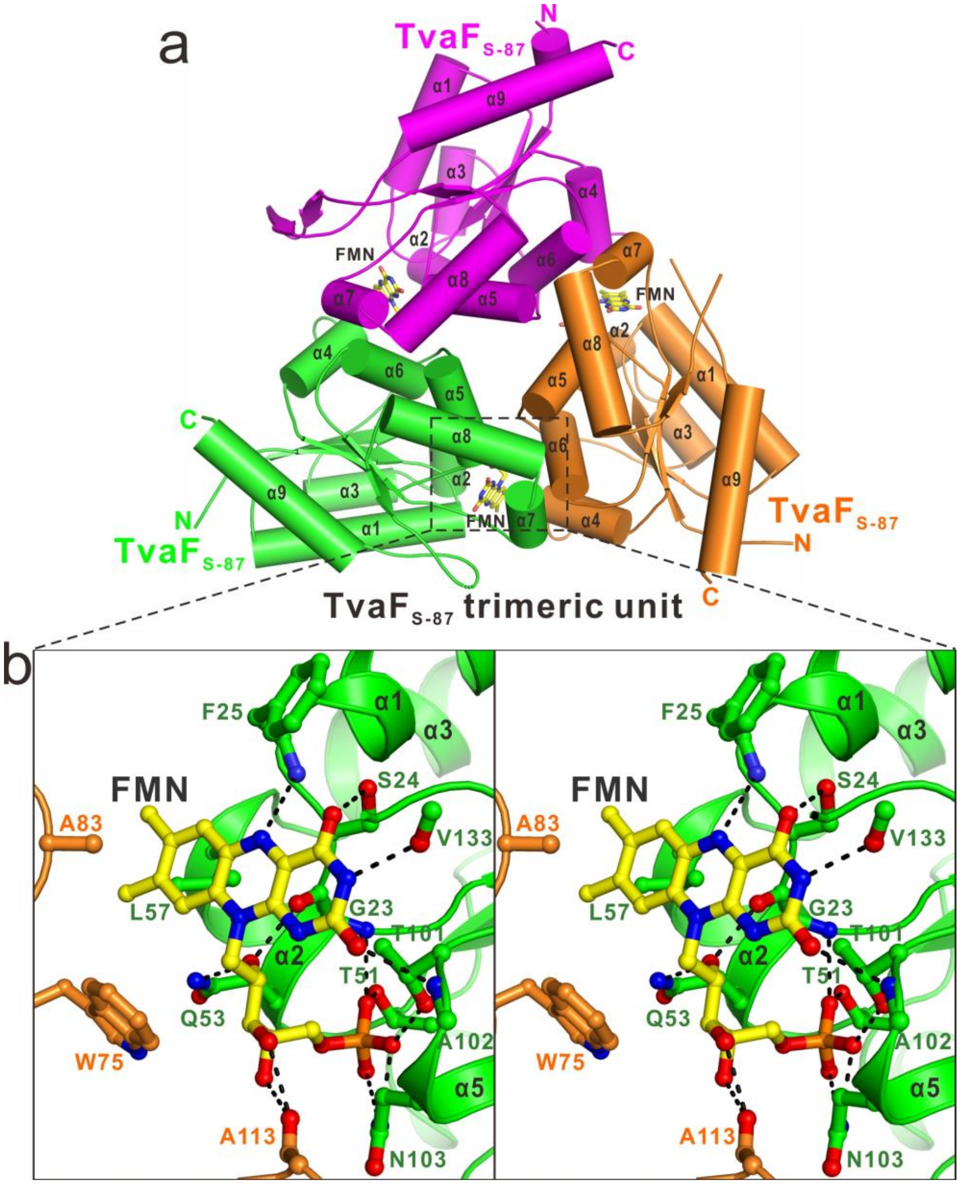

**Supplementary Figure 11.** FAD binding interface in the CypD structure. (**a**) The ribbon-stick model showing the FAD cofactors buried in the interfaces between two CypD monomers in a trimeric unit. The three CypD monmers are shown in the ribbon model and colored in cyan, slate blue, and hotpink, respectively, while the bound FAD cofactors are shown in the stick model. (**b**) The enlarged stereo view of the ribbon-stick-ball representation showing the detailed interactions between a FAD cofactor and two CypD monomers. The hydrogen bonds involved in the interaction are shown as dotted lines

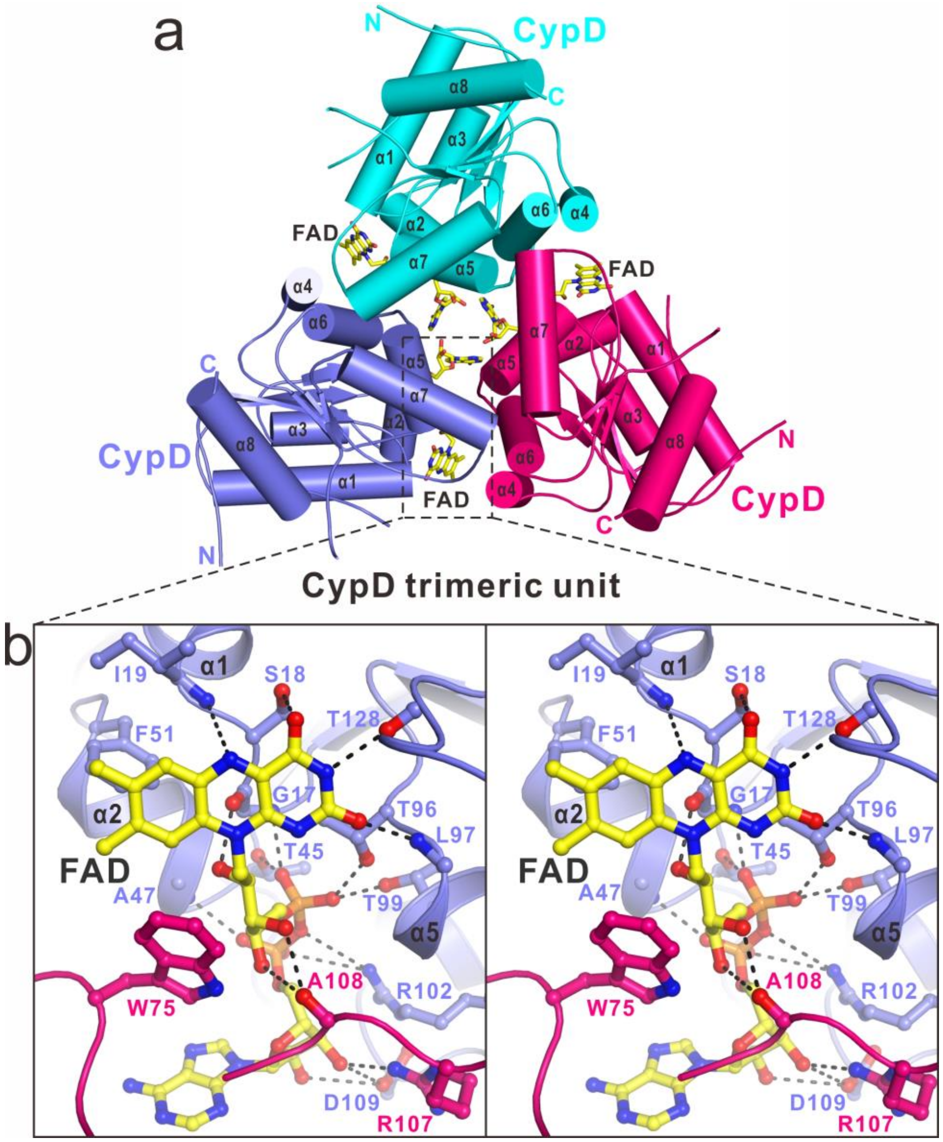

**Supplementary Figure 12.** Structure-based sequence alignment of CypD, EpiD and TvaF$_{S-87}$. The conserved residues are highlighted by colors using software Jalview2.8.1 (http://www.jalview.org/). The interface residues of TvaF$_{S-87}$ and CypD that are critical for the interactions with their bound cofactors are highlighted with red stars (polar interactions) or black triangles (hydrophobic interactions), while the interface residues of TvaF$_{S-87}$ and CypD that are crucial for the interaction between two trimeric subunits are labeled with black stars (polar interactions) or red triangles (hydrophobic interactions). Furthermore, the CypD residues involved in the interaction with **ISLVS** (the pentapeptide sequence of **8**) are labeled with red (polar interactions) or black (hydrophobic interactions) gears.

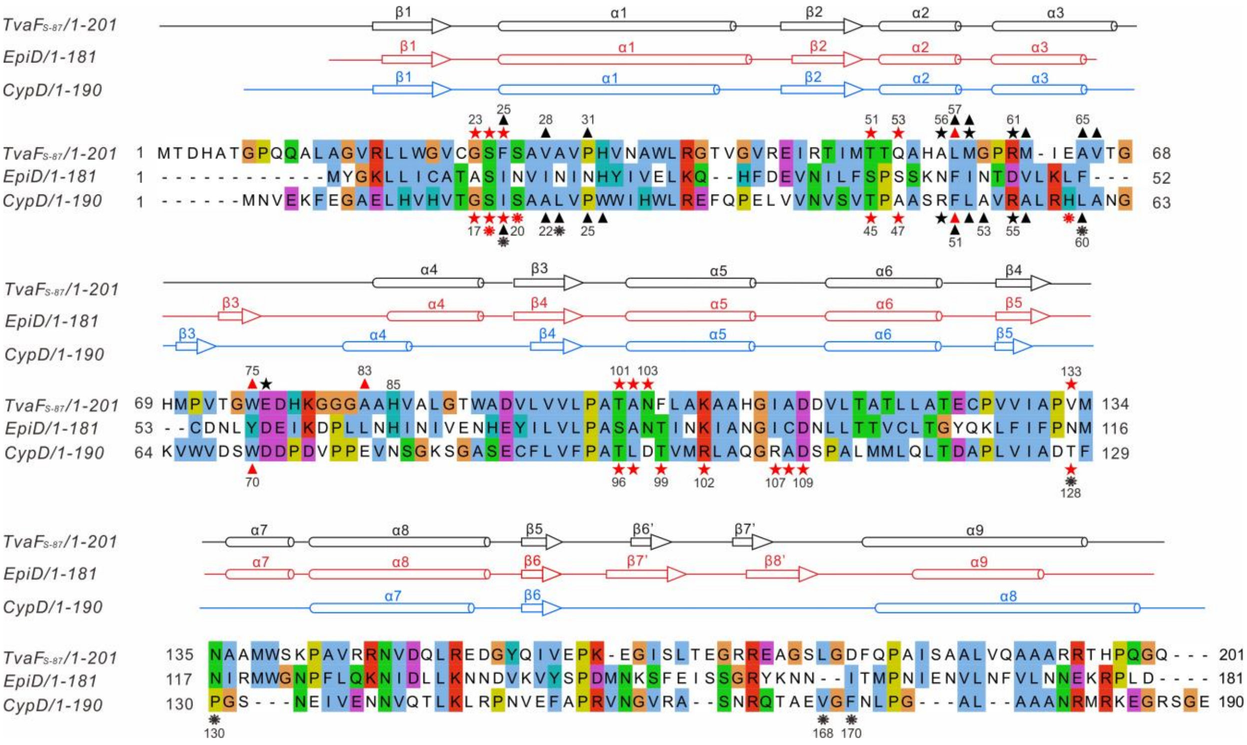

**Supplementary Figure 13.** Structural and biochemical analyses of the binding interface between two trimeric units of the TvaF$_{S-87}$ dodecamer. (**a**) The ribbon-stick model showing the structural packing between two trimeric units. The bound FMN cofactors are shown in the stick model, and the six monomers forming two trimeric subunits are shown in the ribbon model and labeled with different colors. (**b**) The enlarged stereo view of the ribbon-stick representation showing the binding interface between two trimeric units. The hydrogen bonds involved in the interaction are shown as dotted lines. (**c**) Overlay plot of the static light scattering data of wild type TvaF$_{S-87}$, and the TvaF$_{S-87}$-M62D and TvaF$_{S-87}$-V28D mutants. Clearly, the wide type TvaF$_{S-87}$ forms a stable dodecamer, while the two mutants exclusively form a stable trimer.

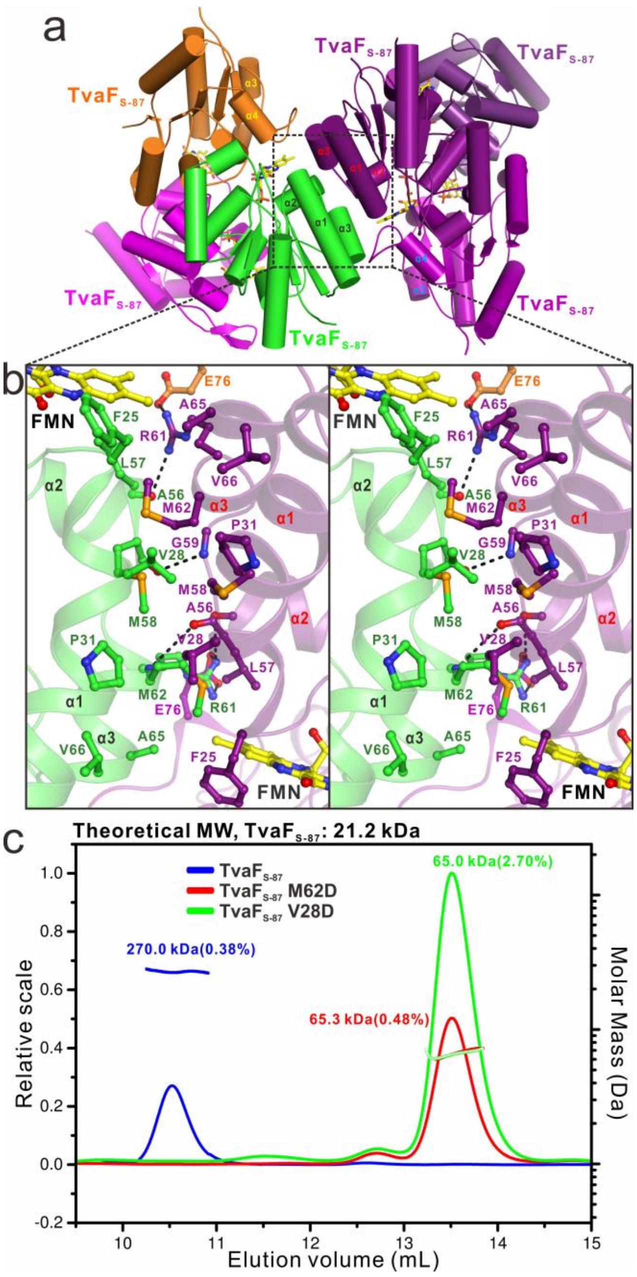

**Supplementary Figure 14.** Structural analysis of the binding interface between two trimeric subunits of the CypD dodecamer. (**a**) The ribbon-stick model showing the structural packing between two trimeric units. In this drawing, The bound FAD cofactors are shown in the stick model, and the six monomers forming two trimeric bunits are shown in the ribbon model and labeled with different colors. (**b**) The enlarged stereo view of the ribbon-stick representation showing the binding interface between two trimeric CypD units. The hydrogen bonds involved in the interaction are shown as dotted lines.

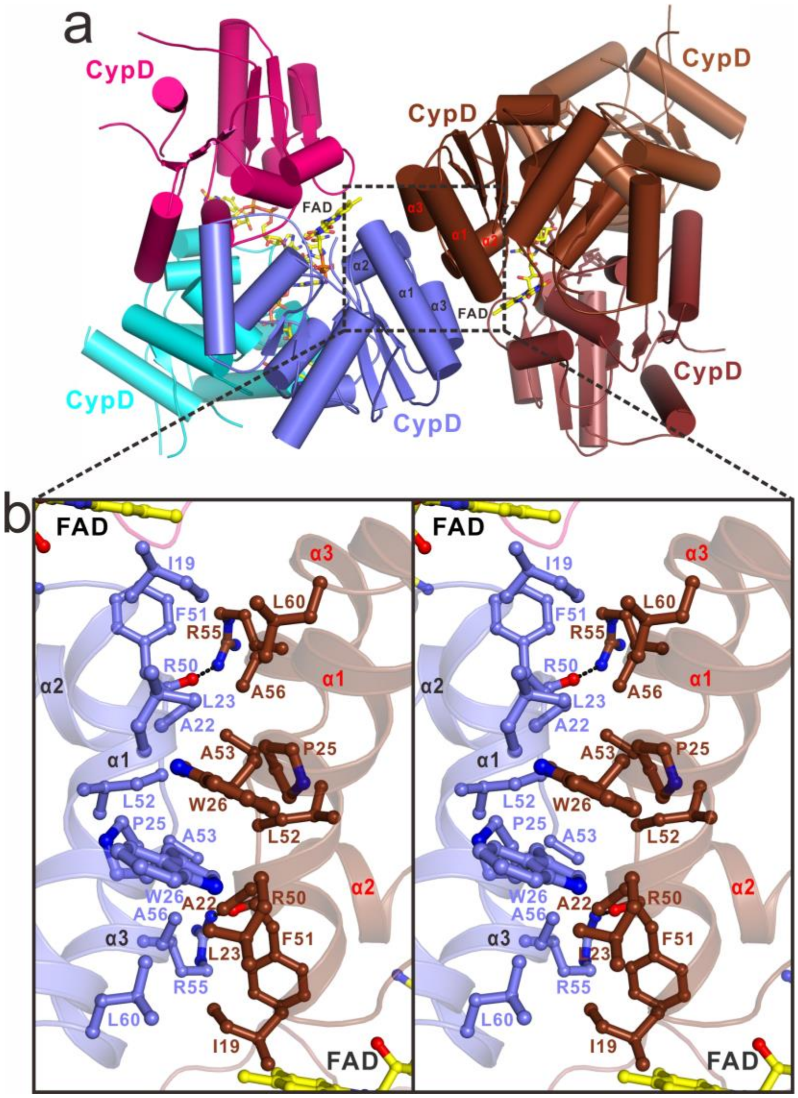

**Supplementary Figure 15.** Structural analysis of the CypD monomer in the CypD complex with the peptide substrate **8**. (**a**) Comparison of the CypD monomer in the complex (purple blue) and in its apo-form (slate blue), respectively. (**b**) The $F_O$-$F_C$ map of the pentapeptide **IS$_{19}$LVS$_{22}$** sequence of **8** in the complex structure. The electron density map is calculated by omitting this sequence from the final PDB file of the complex and contoured at 2.0σ. The CypD is shown in the cartoon mode, while the bound **IS$_{19}$LVS$_{22}$** sequence and FAD are shown in the stick-ball mode.

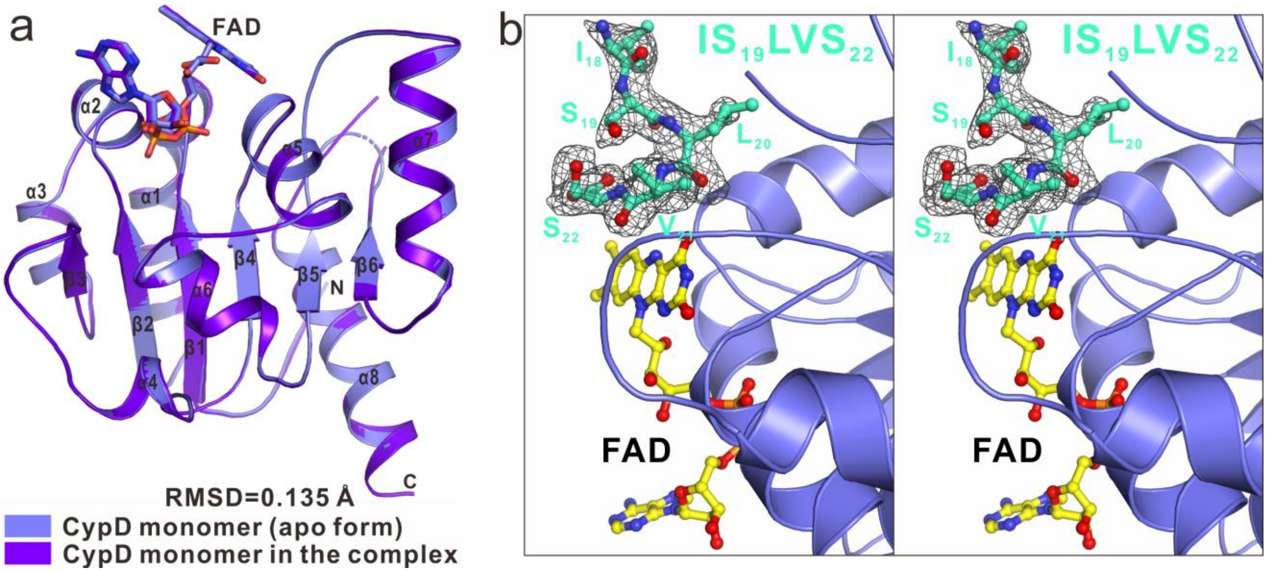

**Supplementary Figure 16.** Structural analysis of the substrate-binding pockets of CypD dodecamer. (**a**) The combined surface representation and the stick-ball model showing the substrate-binding pockets of CypD. (**b**) The combined surface charge potential representation and the stick-ball model showing that the solvent-exposed and highly charged pocket formed between two trimeric units is close to a bound FAD cofactor.

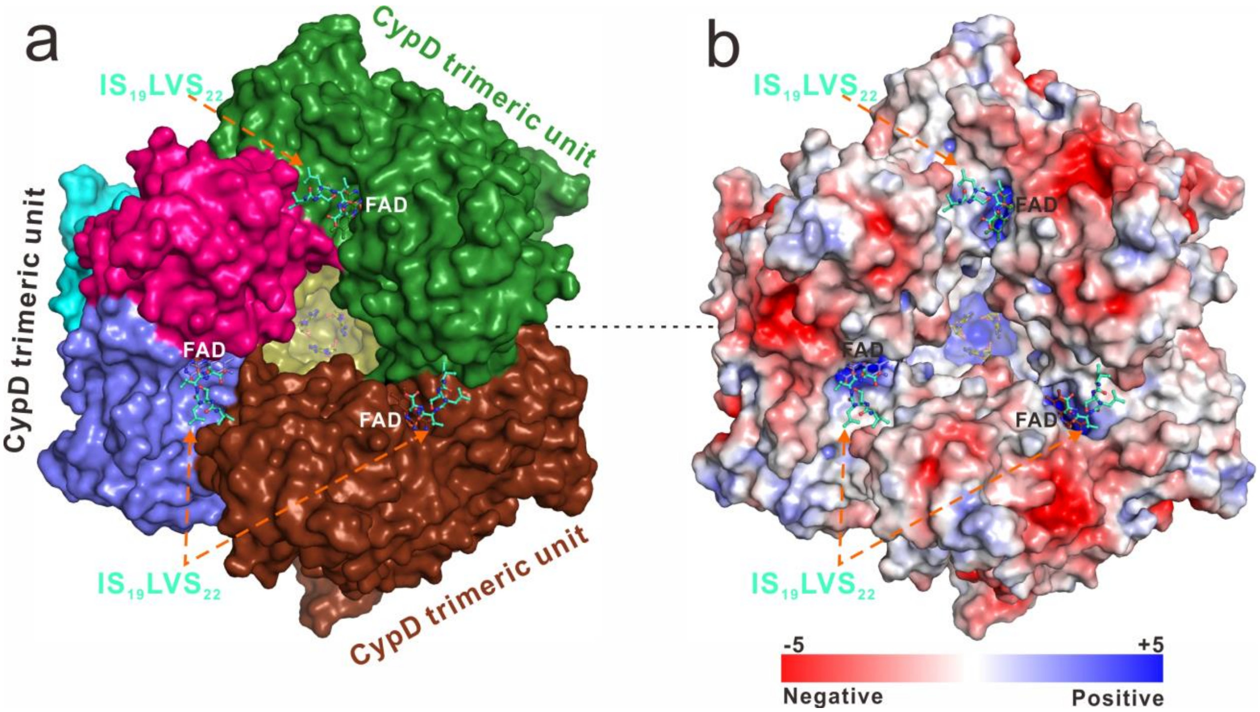

**Supplementary Figure 17.** Structural comparison of the CypD complex and the EpiD H67N mutant complex. (**a**) The superimposition of the CypD complex with the peptide substrate **8** (purple blue and green cyan) and the EpiD H67N mutant complex with a pentapeptide substrate mimic (grey and black). The ribbon-stick model reveals the overlapping of the two putative substrate binding pockets. (**b**) The combined surface representation and the stick-ball model showing the overlaps of the two substrate peptides in the CypD/**8** complex and the EpiD H67N mutant complex.

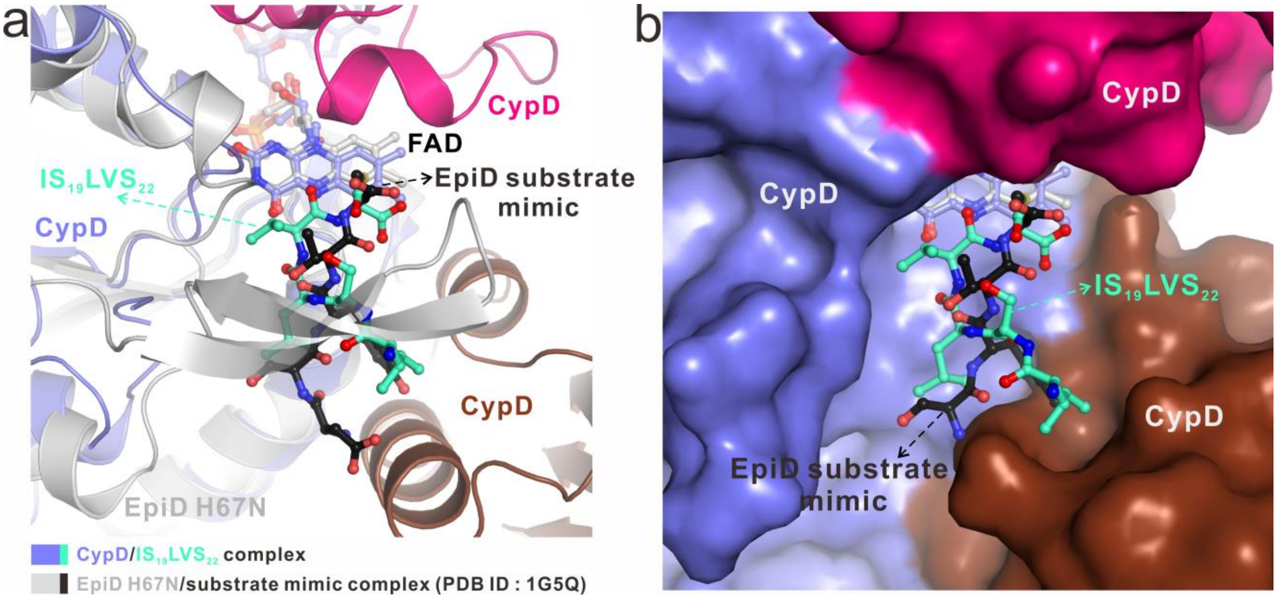

**Supplementary Figure 18.** *In vitro* assays of CypD activity using the substrates, **7** (i), **7-C19S** (ii) and **7-C19S-D$_3$** (containing the deuterium-labeled L-[2,3,3-D$_3$]Ser residue, iii). Conversions were conducted at 30°C for 4 hr in 60 μL of the reaction mixture that contained 100 μM synthetic peptide and 30 μM N-terminally TRX-tagged CypD along with 50 mM Tris-HCl (pH 8.5), 10 mM TCEP, 5 μM FAD and 1 μM 3C. The reaction mixture for **7-C19S-D$_3$** conversion was concentrated ten times for HR -MS analysis.

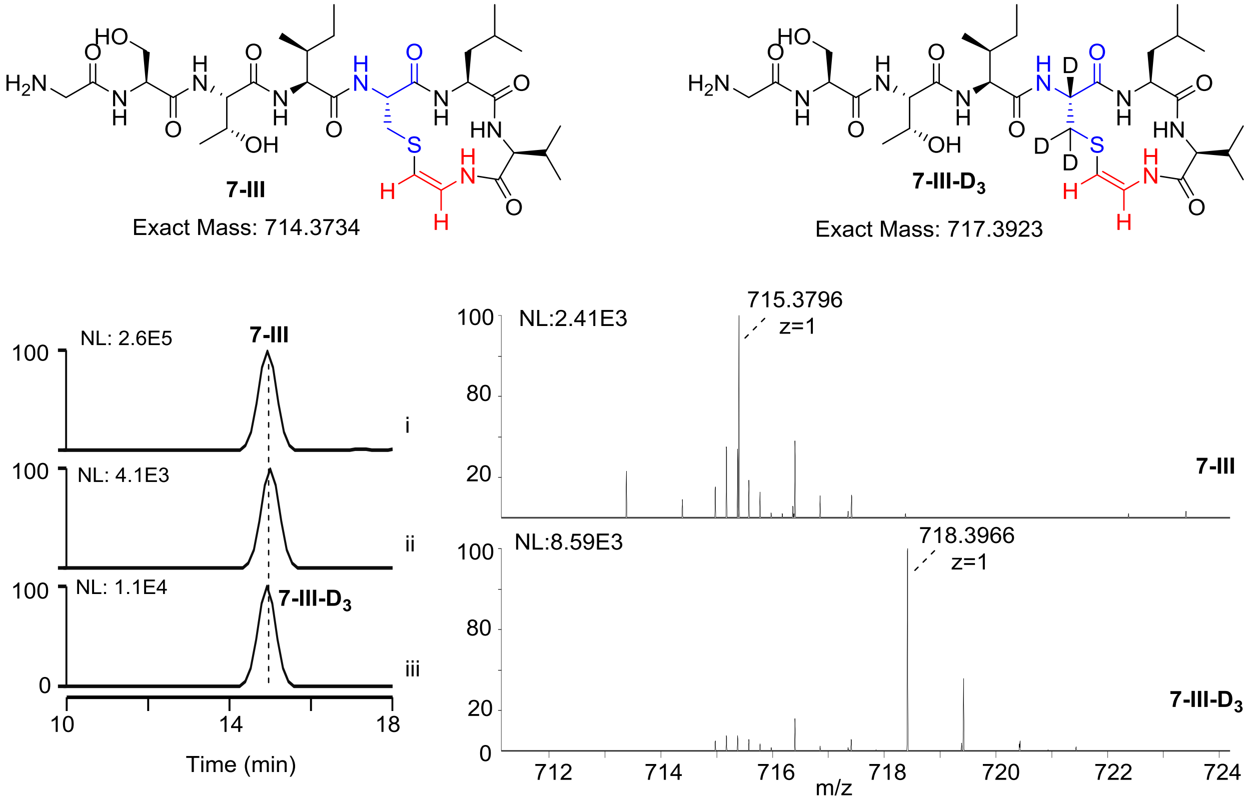

**Supplementary Figure 19.** Characterization of Dha-containing peptides in the CypD-catalyzed conversion of **7-Dha** by chemical derivation. (**a**) Examination of **7-Dha** and its derivatives by HPLC-MS. i, standard **7-Dha**; ii, treatment of **7-Dha** with DTT at 30°C for 2 hr; iii, transformation of **7-Dha** into **7-III** and **7-IV** at 30°C for 2 hr; and iv, treatment of the reaction mixture of iii with DTT. (**b**) HR-MS and HR-MS/MS data for the derivative **7-Dha-DTT**.

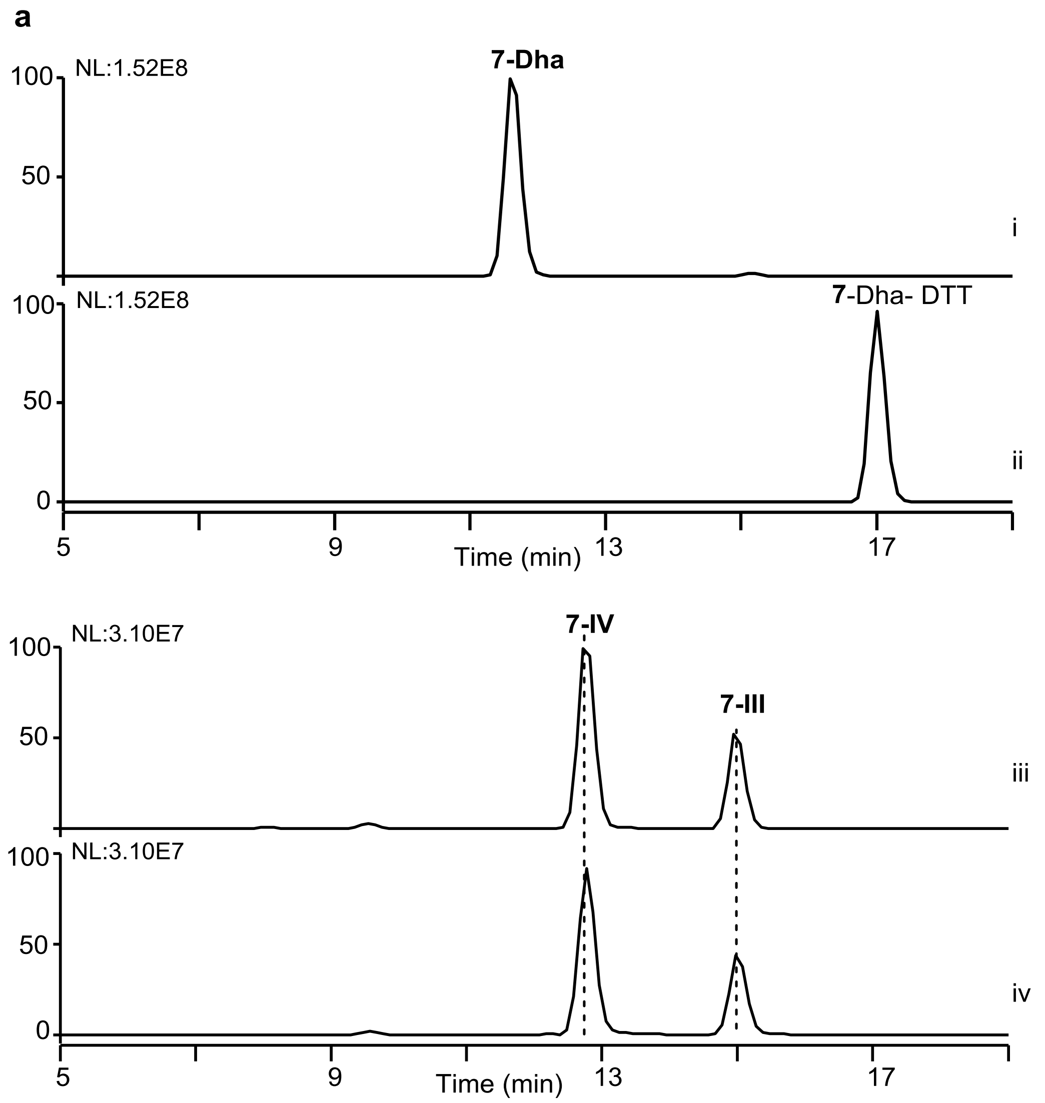

**b**

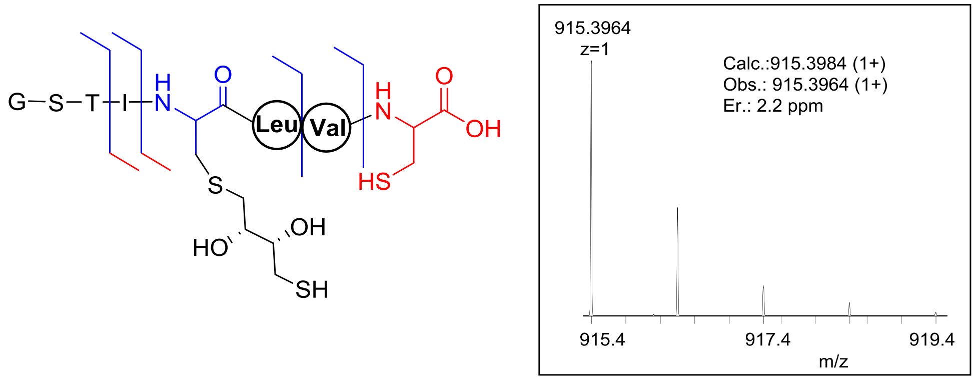


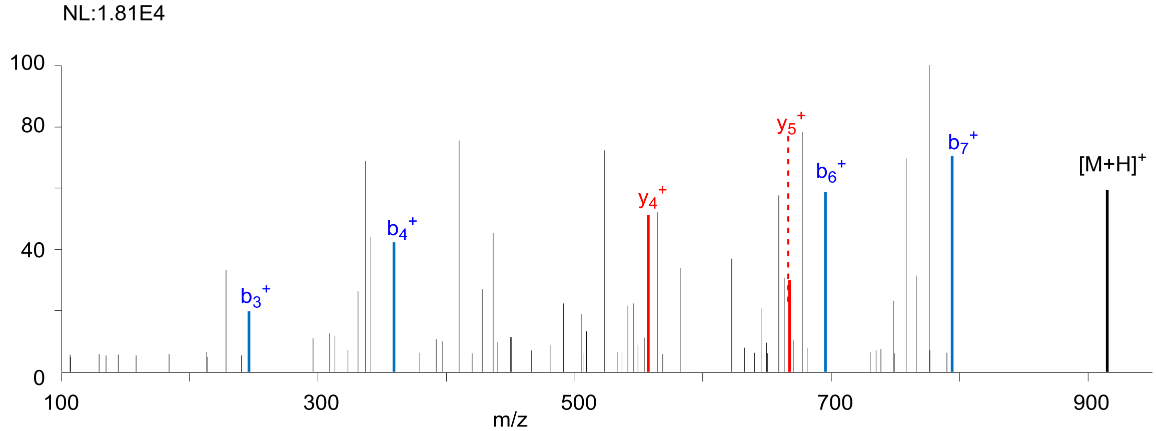


| | Calcd. | Obs. | Er.(ppm) |
|---|---|---|---|
| $b_3^+$ | 246.1090 | 246.1079 | 4.5 |
| $b_4^+$ | 359.1931 | 359.1927 | 1.1 |
| $b_6^+$ | 695.3093 | 695.3108 | 2.2 |
| $b_7^+$ | 794.3787 | 794.3792 | 0.6 |
| $y_4^+$ | 557.2137 | 557.2122 | 2.7 |
| $y_5^+$ | 670.2978 | 670.2968 | 1.5 |

**Supplementary Figure 20.** Comparison of the enzymatic activities of CypD with its variants based on examination of the production of **7-I**, **7-II** and **7-III**. The activity has been quantified by the intensities in HR-MS analysis, in which the activity of wild type CypD to produce **7-1** and **7-II** was normalized to 100%. Assays were performed in triplicates, and the standard deviations are indicated by the error bars. Conversions were conducted at 30°C for 4 hr in 60 μL of the reaction mixture that contained 100 μM **7** and 30 μM N-terminally TRX-tagged CypD, along with 50 mM Tris-HCl (pH 8.5), 10 mM TCEP, 5 μM FAD, 1 μM 3C.

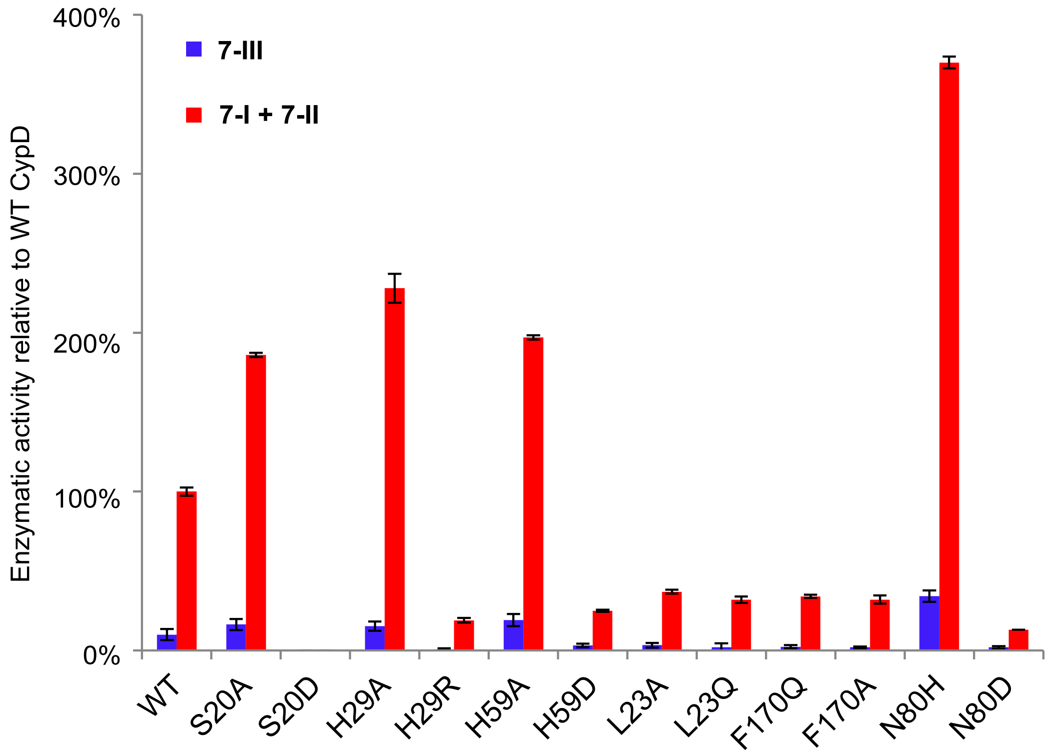

**Supplementary Tables**

**Supplementary Table 1.** Related bacterial strains and plasmids used in this study.

| Strains/Plasmids | Characteristic(s) | Sources/References |
|---|---|---|
| ***Streptomyces* sp.** | | |
| **NRRL S-87** | Wild type strain, TVA-YJ-2-producing strain for PCR | NRRL |
| ***Escherichia coli*** | | |
| DH5α | Host for general cloning | Transgen |
| BL21 (DE3) | Host for protein expression | Transgen |
| JZ101 | BL21 (DE3) derivative, containing pZL 1001 for producing TRX-tagged $TvaF_{S-87}$ | This study |
| JZ102 | BL21 (DE3) derivative, containing pZL 1003for producing TRX-tagged CypD | This study |
| JZ103 | BL21 (DE3) derivative, containing pZL 1002 for producing 3C | This study |
| JZ104 | BL21 (DE3) derivative, containing pZL 1004 for producing TRX-tagged $TvaF_{S-87}$- V28D | This study |
| JZ105 | BL21 (DE3) derivative, containing pZL1005 for producing TRX-tagged $TvaF_{S-87}$-M62D | This study |
| JZ106 | BL21 (DE3) derivative, containing pZL 1006 for producing TRX-tagged $TvaF_{S-87}$- H85A | This study |
| JZ107 | BL21 (DE3) derivative, containing pZL 1007 for producing TRX-tagged CypD-S20A | This study |
| JZ108 | BL21 (DE3) derivative, containing pZL 1008 for producing TRX-tagged CypD-S20D | This study |
| JZ109 | BL21 (DE3) derivative, containing pZL 1009 for producing TRX-tagged CypD-L23A | This study |
| JZ110 | BL21 (DE3) derivative, containing pZL 1010 for producing TRX-tagged CypD-L23Q | This study |
| JZ111 | BL21 (DE3) derivative, containing pZL 1011 for producing TRX-tagged CypD-F170A | This study |
| JZ112 | BL21 (DE3) derivative, containing pZL 1012 for producing TRX-tagged CypD-F170Q | This study |

| | | |
|---|---|---|
| JZ113 | BL21 (DE3) derivative, containing pZL 1013 for producing TRX-tagged CypD-H59A | This study |
| JZ114 | BL21 (DE3) derivative, containing pZL 1014 for producing TRX-tagged CypD-H59D | This study |
| JZ115 | BL21 (DE3) derivative, containing pZL 1015 for producing TRX-tagged CypD-H29R | This study |
| JZ116 | BL21 (DE3) derivative, containing pZL 1016 for producing TRX-tagged CypD-H29A | This study |
| JZ117 | BL21 (DE3) derivative, containing pZL 1017 for producing TRX-tagged CypD-N80H | This study |
| JZ118 | BL21 (DE3) derivative, containing pZL 1018 for producing TRX-tagged CypD- N80D | This study |
| **Plasmids** | *E. coli* subcloning vector | |
| pET28a(+) | Protein expression vector used in *E.coli*, encoding N-terminal His-tag, kanamycin resistance | Novagen |
| pRSFDeut-1 | Protein expression vector used in *E.coli*, encoding N-terminal His-tag, kanamycin resistance | Novagen |
| pZL1001 | pRSFDeut-1 derivative, containing *trx* and a 606 bp PCR product that encodes $tvaF_{S\text{-}87}$ | This study |
| pZL1002 | pET28a(+) derivative, containing a 552 bp synthesized gene that encodes *3c* | This study |
| pZL1003 | pRSFDeut-1 derivative, containing *trx* and a 573 bp synthesized gene that encodes *cypD* | This study |
| pZL1004 | pZL1001 derivative for V28D mutated $tvaF_{S\text{-}87}$ | This study |
| pZL1005 | pZL1001 derivative for M62D mutated $tvaF_{S\text{-}87}$ | This study |
| pZL1006 | pZL1001 derivative for H85A mutated $tvaF_{S\text{-}87}$ | This study |
| pZL1007 | pZL1003 derivative for S20A mutated *cypD* | This study |
| pZL1008 | pZL1003 derivative for S20D mutated *cypD* | This study |
| pZL1009 | pZL1003 derivative for L23A mutated *cypD* | This study |
| pZL1010 | pZL1003 derivative for L23Q mutated *cypD* | This study |
| pZL1011 | pZL1003 derivative for F170A mutated *cypD* | This study |
| pZL1012 | pZL1003 derivative for F170Q mutated *cypD* | This study |
| pZL1013 | pZL1003 derivative for H59D mutated *cypD* | This study |
| pZL1014 | pZL1003 derivative for H59A mutated *cypD* | This study |
| pZL1015 | pZL1003 derivative for H29R mutated *cypD* | This study |

| | | |
|---|---|---|
| pZL1016 | pZL1003 derivative for H29A mutated *cypD* | This study |
| pZL1017 | pZL1003 derivative for N80H mutated *cypD* | This study |
| pZL1018 | pZL1003 derivative for N80D mutated *cypD* | This study |

**Supplementary Table 2.** Primers used in this study.

| **Primers** | **Sequence (restriction sites are underlined)** |
|---|---|
| *tvaF*$_{S\text{-}87}$ -for | AAAAAGGATCCATGACCGACCACGCCACCG |
| *tvaF*$_{S\text{-}87}$ -rev | AAAAAGAATTCTCACTGTCCTTGTGGGTG |
| *tvaF*$_{S\text{-}87}$-V28D-for | TCCGCTGACGCGGTCCCCCACGTGAAC |
| *tvaF*$_{S\text{-}87}$-V28D-rev | GACCGGTCCAGCGGAGAAGGAGCCGCA |
| *tvaF*$_{S\text{-}87}$-M62D-for | CCGCGCGACATCGAGGCTGTCACCGGC |
| *tvaF*$_{S\text{-}87}$-M62D-rev | CTCGATGTCGCGCGGCCCCATGAGGGC |
| *tvaF*$_{S\text{-}87}$-H85A-for | GCCGCCGCAGTCGCCCTCGGCACCTGG |
| *tvaF*$_{S\text{-}87}$-H85A-rev | GGCGACTGCGGCGGCGCCTCCGCCCTTG |
| *cypD*-S20A-for | AGCATCGCAGCGGCGCTGGTTCCGTGG |
| *cypD*-S20A-rev | CGCCGCTGCGATGCTGCCGGTAACGTG |
| *cypD*-S20D-for | AGCATCGACGCGGCGCTGGTTCCGTGG |
| *cypD*-S20D-rev | CGCCGCGTCGATGCTGCCGGTAACGTG |
| *cypD*-L23A-for | GCGGCGGCAGTTCCGTGGTGGATTCAC |
| *cypD*-L23A-rev | CGGAACTGCCGCCGCGCTGATGCTGCC |
| *cypD*-L23Q-for | GCGGCGCAAGTTCCGTGGTGGATTCAC |
| *cypD*-L23Q-rev | CGGAACTTGCGCCGCGCTGATGCTGCC |
| *cypD*-F170A-for | GTGGGTGCAAACCTGCCGGGTGCGCTG |
| *cypD*-F170A-rev | CAGGTTTGCACCCACTTCCGCGGTTTG |
| *cypD*-F170Q-for | GTGGGTCAAAACCTGCCGGGTGCGCTG |
| *cypD*-F170Q-rev | CAGGTTTTGACCCACTTCCGCGGTTTG |
| *cypD*-H59D-for | CTGCGTGACCTGGCGAACGGCAAAGTG |
| *cypD*-H59D-rev | CGCCAGGTCACGCAGCGCACGCACCGC |
| *cypD*-H59A-for | CTGCGTGCACTGGCGAACGGCAAAGTG |
| *cypD*-H59A-rev | CGCCAGTGCACGCAGCGCACGCACCGC |
| *cypD*-H29R-for | TGGATTAGATGGCTGCGTGAGTTCCAG |
| *cypD*-H29R-rev | CAGCCATCTAATCCACCACGGAACCAG |
| *cypD*-H29A-for | TGGATTGCATGGCTGCGTGAGTTCCAG |
| *cypD*-H29A-rev | CAGCCATGCAATCCACCACGGAACCAG |
| *cypD*-N80H-for | GAAGTTCACAGCGGTAAAAGCGGCGCG |
| *cypD*-N80H-rev | ACCGCTGTGAACTTCCGGCGGCACATC |
| *cypD*-N80D-for | GAAGTTGACAGCGGTAAAAGCGGCGCG |

| | |
|---|---|
| *CypD*N80D-rev | ACCGCTGTCAACTTCCGGCGGCACATC |

**Supplementary Table 3.** Gene sequences of *tvaF*$_{S\text{-}87}$, *cypD* and *3c*.

| Name | Sequence |
|---|---|
| *tvaF*$_{S\text{-}87}$ | ATGACCGACCACGCCACCGGCCCGCAGCAGGCACTGGCGGGGGTTCGGT<br>TGTTGTGGGGCGTGTGCGGCTCCTTCTCCGCTGTTGCGGTCCCCCACGTG<br>AACGCATGGCTGCGTGGCACCGTCGGGGTCCGGGAGATCCGCACCATCA<br>TGACCACGCAGGCACACGCCCTCATGGGGCCGCGCATGATCGAGGCTGT<br>CACCGGCCATATGCCGGTGACCGGCTGGGAGGACCACAAGGGCGGAGG<br>CGCCGCCCACGTCGCCCTCGGCACCTGGGCGGACGTGCTGGTGGTCTTGC<br>CGGCCACCGCCAATTTCCTGGCCAAGGCAGCACACGGCATCGCGGACGA<br>CGTCCTGACAGCCACACTGCTCGCCACCGAGTGCCCCGTGGTGATCGCCC<br>CCGTCATGAACGCGGCCATGTGGTCCAAACCCGCCGTACGCCGCAACGT<br>CGATCAGCTCCGCGAAGACGGCTACCAGATCGTCGAGCCGAAGGAGGGC<br>ATCTCCCTCACCGAGGGCCGACGGGAAGCCGGTTCACTCGGCGACTTCC<br>AGCCGGCAATCTCCGCCGCTCTGGTCCAGGCAGCTGCACGACGCACCCA<br>CCCACAAGGACAGTGA |
| *cypD* | ATGAACGTGGAGAAGTTCGAGGGTGCGGAGCTGCACGTGCACGTTACCG<br>GCAGCATCAGCGCGGCGCTGGTTCCGTGGTGGATTCACTGGCTGCGTGA<br>GTTCCAGCCGGAGCTGGTTGTGAACGTGAGCGTTACCCCGGCGGCGAGC<br>CGTTTTCTGGCGGTGCGTGCGCTGCGTCACCTGGCGAACGGCAAAGTGTG<br>GGTTGACAGCTGGGACGATCCGGATGTGCCGCCGGAAGTTAACAGCGGT<br>AAAAGCGGCGCGAGCGAGTGCTTCCTGGTGTTTCCGGCGACCCTGGACA<br>CCGTTATGCGTCTGGCGCAGGGTCGTGCGGATAGCCCGGCGCTGATGAT<br>GCTGCAACTGACCGACGCGCCGCTGGTTATCGCGGATACCTTTCCGGGCA<br>GCAACGAAATTGTGGAGAACAACGTTCAGACCCTGAAACTGCGTCCGAA<br>CGTGGAGTTCGCGCCGCGTGTGAACGGTGTTCGTGCGAGCAACCGTCAA<br>ACCGCGGAAGTGGGTTTTAACCTGCCGGGTGCGCTGGCGGCGGCGAACC<br>GTATGCGTAAAGAAGGTCGTAGCGGCGAGTAA |
| *3c* | ATGGCTAGCATGACTGGTGGACAGCAAATGGGTCGCGGATCCGGACCTA<br>ACACTGAGTTTGCTCTGTCTCTGCTGCGTAAAAACATCATGACTATCACC<br>ACTAGTAAAGGCGAGTTCACTGGTCTGGGTATTCACGATCGTGTTTGTGT<br>TATTCCTACTCATGCTCAGCCGGGTGACGATGTTCTGGTAAACGGTCAAA<br>AAATTCGTGTTAAGGATAAATACAAACTGGTTGACCCGGAAAACATCAA<br>TCTAGAACTGACCGTACTGACTCTGGATCGTAATGAAAAGTTCCGTGACA<br>TCCGTGGTTTTATTTCTGAAGACCTGGAAGGTGTCGACGCAACCCTGGTT<br>GTACATAGCAATAACTTTACTAACACTATTCTGGAGGTTGGTCCGGTAAC<br>TATGGCTGGTCTGATCAACCTGTCTAGCACTCCGACCAACCGCATGATTC<br>GTTACGACTACGCAACTAAAACTGGTCAGTGTGGTGGTGTTCTGTGCGCA<br>ACCGGTAAGATCTTTGGCATCCATGTAGGCGGTAACGGTCGTCAGGGTTT<br>CTCTGCACAACTGAAGAAGCAATACTTTGTAGAGAAGCAGTAA |

**Supplementary Table 4.** HR-MS and HR-MS/MS Data collection of all compounds analyzed in this study. _ _, not observed; and N.D., not detected.

| **Peptidyl mimics (PM)** | **Calcd.** | **Obs.** | **Er. ppm** |
|---|---|---|---|
| **1** 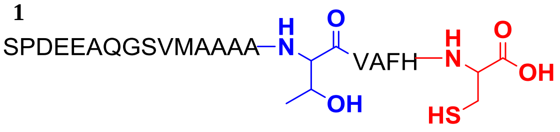  | 1046.4596 $[M+2H]^{2+}$ | 1046.4606 $[M+2H]^{2+}$ | 1.0 |
| **1-I** 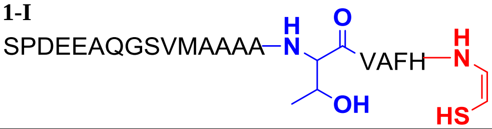  | 1023.4569 $[M+2H]^{2+}$ | 1023.4562 $[M+2H^{+}]^{2+}$ | 0.7 |
| **1-III** 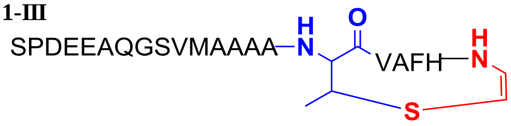  | 1014.4516 $[M+2H]^{2+}$ | 1014.4581 $[M+2H]^{2+}$ | 6.4 |
| **1-II** 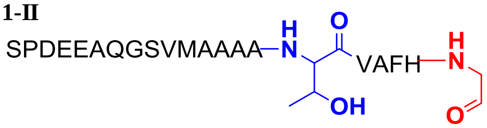  | 1015.4684 $[M+2H]^{2+}$ | 1015.4662 $[M+2H]^{2+}$ | 2.2 |
| **1-T8A** 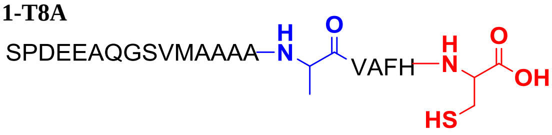  | 1031.4544 $[M+2H]^{2+}$ | 1031.4557 $[M+2H]^{2+}$ | 1.3 |
| **1-T8A-I** 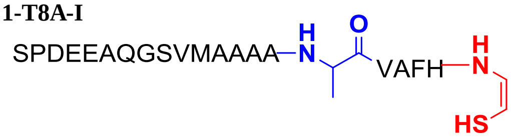  | 1008.4517 $[M+2H]^{2+}$ | 1008.4541 $[M+2H]^{2+}$ | 2.4 |
| **1-T8A-II** 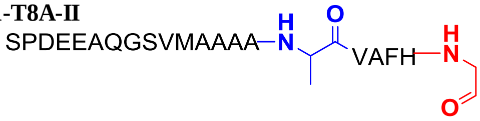  | 1000.4631 $[M+2H]^{2+}$ | 1000.4646 $[M+2H]^{2+}$ | 1.5 |
| **1-T8S** 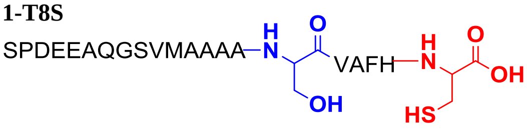  | 1039.4519 $[M+2H]^{2+}$ | 1039.4493 $[M+2H]^{2+}$ | 2.5 |
| **1-T8S-I** 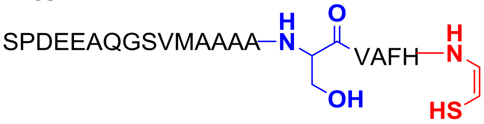  | 1016.4491 $[M+2H]^{2+}$ | 1016.4470 $[M+2H]^{2+}$ | 2.1 |

| | | | |
|---|---|---|---|
| **1-T8S-III** 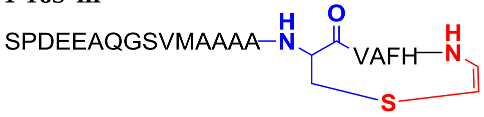  | 1007.4438 $[M+2H]^{2+}$ | 1007.4490 $[M+2H]^{2+}$ | 5.2 |
| **1-T8S-II** 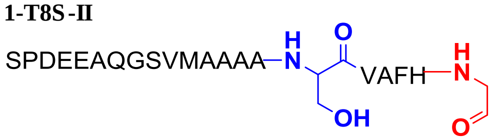  | 1008.4605 $[M+2H]^{2+}$ | 1008.4579 $[M+2H]^{2+}$ | 2.6 |
| **1-T8C** 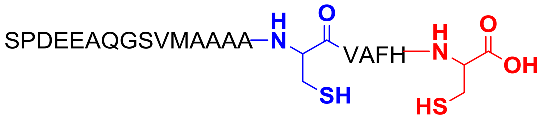  | 1047.4404 $[M+2H]^{2+}$ | 1047.4406 $[M+2H]^{2+}$ | 0.2 |
| **1-T8C-I** 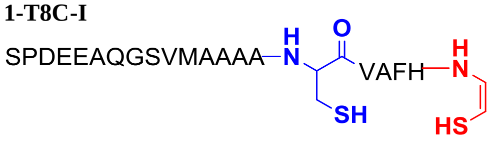  | 1024.4377 $[M+2H]^{2+}$ | 1024.4386 $[M+2H]^{2+}$ | 0.9 |
| **1-T8C-III** 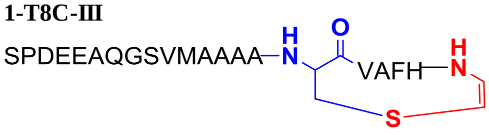  | 1007.4438 $[M+2H]^{2+}$ | – – | – – |
| **1-T8C-II** 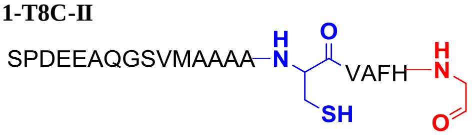  | 1016.4492 $[M+2H]^{2+}$ | 1016.4494 $[M+2H]^{2+}$ | 0.2 |
| **1-C13A** 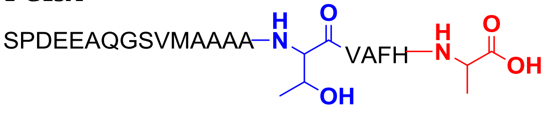  | 1030.4737 $[M+2H]^{2+}$ | 1030.4764 $[M+2H]^{2+}$ | 2.7 |
| **1-C13A-I** 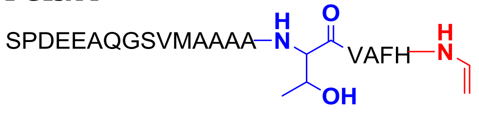  | 1007.4709 $[M+2H]^{2+}$ | – – | – – |
| **1-C13S** 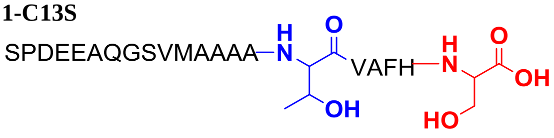  | 1038.4711 $[M+2H]^{2+}$ | 1038.4697 $[M+2H]^{2+}$ | 1.4 |
| **1-C13S-I** 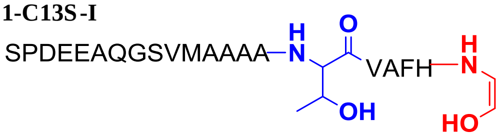  | 1015.4684 $[M+2H]^{2+}$ | – – | – – |

| | | | |
|---|---|---|---|
| **1-C13T** 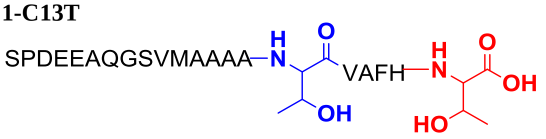 | 1045.4790 $[M+2H]^{2+}$ | 1045.4780 $[M+2H]^{2+}$ | 1.0 |
| **1-C13T-I** 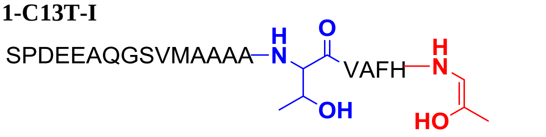 | 1022.4762 $[M+2H]^{2+}$ | – – | – – |
| **1-NEM** 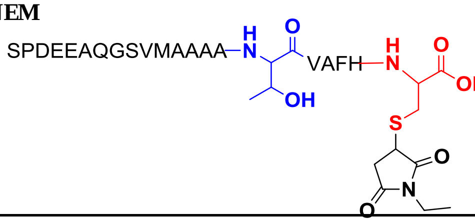 | 1108.9835 $[M+2H]^{2+}$ | 1108.9807 $[M+2H]^{2+}$ | 2.5 |
| **1-I-NEM** 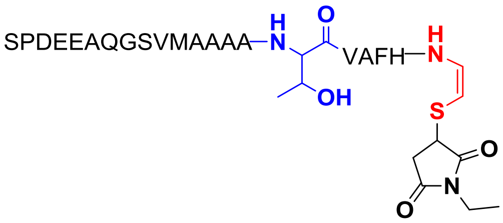 | 1085.9807 $[M+2H]^{2+}$ | 1085.9799 $[M+2H]^{2+}$ | 0.7 |
| **1-II-HOPI** 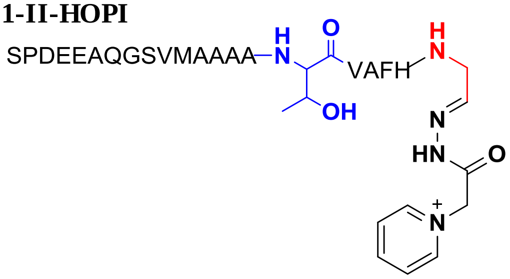 | 1082.0001 $[M+2H]^{2+}$ | 1081.9943 $[M+2H]^{2+}$ | 5.3 |
| **2** 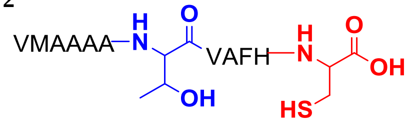 | 596.2866 $[M+2H]^{2+}$ | 596.2846 $[M+2H]^{2+}$ | 3.4 |
| **2-I** 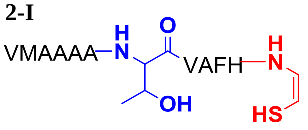 | 573.2839 $[M+2H]^{2+}$ | 573.2809 $[M+2H]^{2+}$ | 5.2 |
| **2-III** 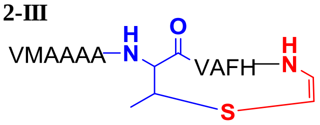 | 564.2786 $[M+2H]^{2+}$ | 564.2836 $[M+2H]^{2+}$ | 8.9 |

| | | | |
|---|---|---|---|
| **2-II** 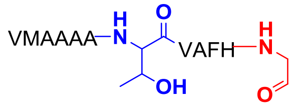 | 565.2953 [M+2H]$^{2+}$ | 565.2923 [M+2H]$^{2+}$ | 5.7 |
| **3** 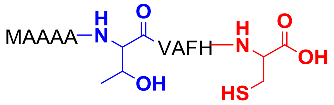 | 546.7524 [M+2H]$^{2+}$ | 546.7499 [M+2H]$^{2+}$ | 4.6 |
| **3-I** 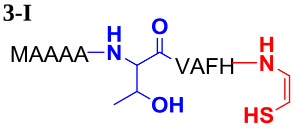 | 523.7497 [M+2H]$^{2+}$ | 523.7488 [M+2H]$^{2+}$ | 1.7 |
| **3-III** 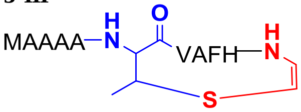 | 514.7444 [M+2H]$^{2+}$ | 514.7419 [M+2H]$^{2+}$ | 4.9 |
| **3-II** 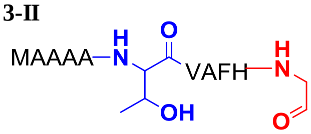 | 515.7611 [M+2H]$^{2+}$ | 515.7584 [M+2H]$^{2+}$ | 5.2 |
| **4** 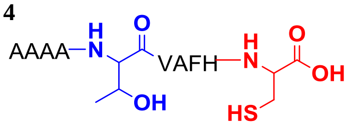 | 961.4566 [M+H]$^{+}$ | 961.4536 [M+H]$^{+}$ | 3.1 |
| **4-I** 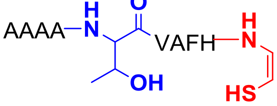 | 915.4511 [M+H]$^{+}$ | 915.4510 [M+H]$^{+}$ | 0.1 |
| **4-III** 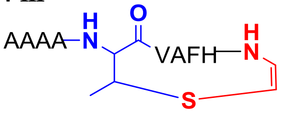 | 897.4405 [M+H]$^{+}$ | – – | – – |
| **4-II** 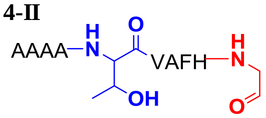 | 899.4739 [M+H]$^{+}$ | N.D. | N.D. |
| **5** 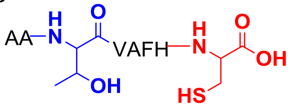 | 819.3823 [M+H]$^{+}$ | 819.3790 [M+H]$^{+}$ | 4.0 |

| | | | |
|---|---|---|---|
| **5-I** 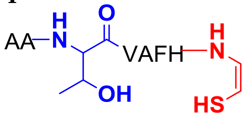 | 773.3769<br>$[M+H]^+$ | 773.3810<br>$[M+H]^+$ | 5.3 |
| **5-III** 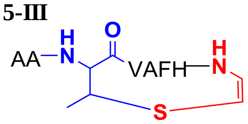 | 755.3663<br>$[M+H]^+$ | – – | – – |
| **5-II** 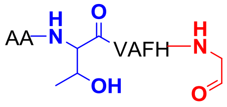 | 757.3997<br>$[M+H]^+$ | N.D. | N.D. |
| **6** 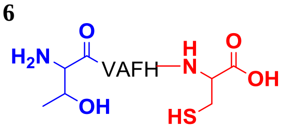 | 677.3081<br>$[M+H]^+$ | 677.3058<br>$[M+H]^+$ | 3.4 |
| **6-I** 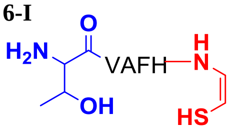 | 631.3026<br>$[M+H]^+$ | 631.3067<br>$[M+H]^+$ | 6.5 |
| **6-III** 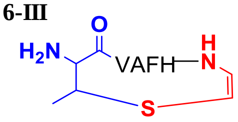 | 613.2921<br>$[M+H]^+$ | – – | – – |
| **6-II** 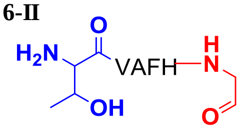 | 615.3255<br>$[M+H]^+$ | N.D. | N.D. |
| **7** 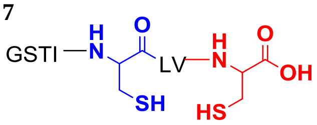 | 795.3739<br>$[M+H]^+$ | 795.3726<br>$[M+H]^+$ | 1.6 |
| **7-I** 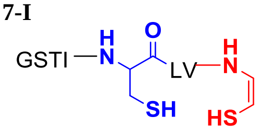 | 749.3684<br>$[M+H]^+$ | 749.3668<br>$[M+H]^+$ | 2.1 |
| **7-III/IV** 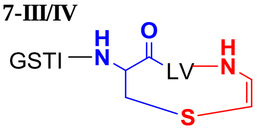 | 715.3808<br>$[M+H]^+$ | 715.3805<br>$[M+H]^+$ | 0.4 |
| **7-II** 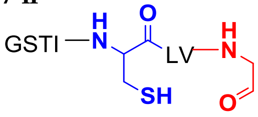 | 733.3913<br>$[M+H]^+$ | 733.3898<br>$[M+H]^+$ | 2.1 |

| | | | |
|---|---|---|---|
| **7-NEM** 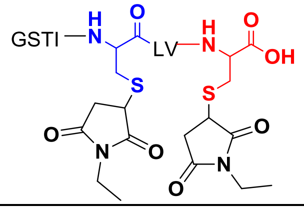 | 1045.4693 [M+H]+ | 1045.4646 [M+H]+ | 4.5 |
| **7-I-NEM** 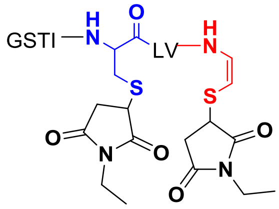 | 999.4638 [M+H]+ | 999.4646 [M+H]+ | 0.8 |
| **7-C19S** 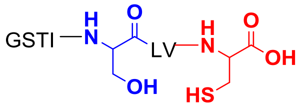 | 779.3968 [M+H]+ | 779.3934 [M+H]+ | 4.4 |
| **7-C19S-I** 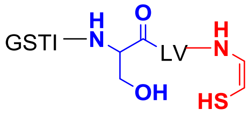 | 733.3913 [M+H]+ | 733.3911 [M+H]+ | 0.3 |
| **7-C19S-III** 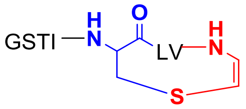 | 715.3808 [M+H]+ | 715.3802 [M+H]+ | 0.8 |
| **7-C19S-II** 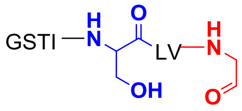 | 717.4141 [M+H]+ | 717.4122 [M+H]+ | 2.7 |
| **7-C19T** 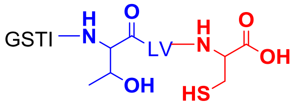 | 793.4124 [M+H]+ | 793.4112 [M+H]+ | 1.5 |
| **7-C19T-I** 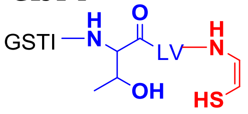 | 747.4069 [M+H]+ | 747.4070 [M+H]+ | 0.1 |
| **7-C19T-III** 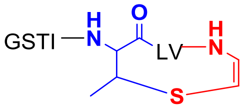 | 729.3965 [M+H]+ | – – | – – |

| | | | |
|---|---|---|---|
| **7-C19T-II** 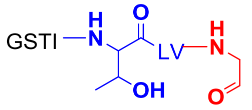 | 731.4298 [M+H]$^+$ | 731.4286 [M+H]$^+$ | 1.6 |
| **7-C19A** 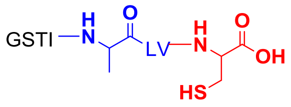 | 763.4019 [M+H]$^+$ | 763.4012 [M+H]$^+$ | 0.9 |
| **7-C19A-I** 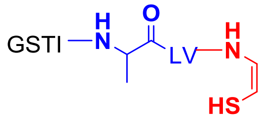 | 717.3964 [M+H]$^+$ | 717.3962 [M+H]$^+$ | 0.3 |
| **7-C19A-II** 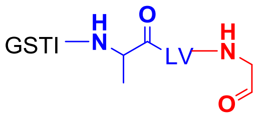 | 701.4192 [M+H]$^+$ | 701.4181 [M+H]$^+$ | 1.6 |
| **7-C22A** 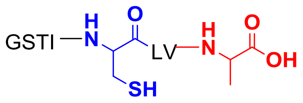 | 763.4019 [M+H]$^+$ | 763.4013 [M+H]$^+$ | 0.8 |
| **7- C22A-I** 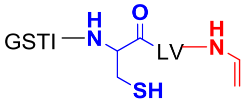 | 717.3964 [M+H]$^+$ | – – | – – |
| **7-C22S** 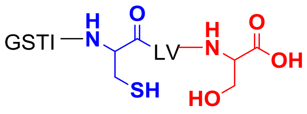 | 779.3968 [M+H]$^+$ | 779.3947 [M+H]$^+$ | 2.7 |
| **7- C22S-I** 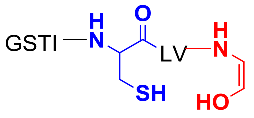 | 733.3913 [M+H]$^+$ | – – | – – |
| **7-C22T** 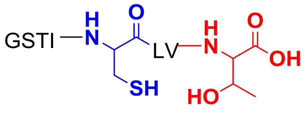 | 793.4124 [M+H]$^+$ | 793.4112 [M+H]$^+$ | 1.5 |
| **7- C22T-I** 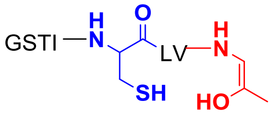 | 747.4069 [M+H]$^+$ | – – | – – |

| | | | |
|---|---|---|---|
| **7-C19S-Ac**<br>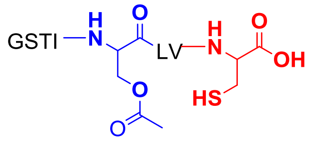 | 821.4073<br>$[M+H]^+$ | 821.4020<br>$[M+H]^+$ | 6.5 |
| **7-C19S-Ac-I**<br>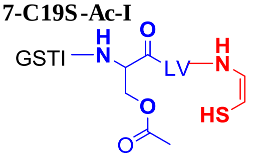 | 775.4019<br>$[M+H]^+$ | 775.3988<br>$[M+H]^+$ | 4.0 |
| **7-C19S-Ac-II**<br>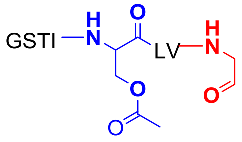 | 759.4247<br>$[M+H]^+$ | 759.4232<br>$[M+H]^+$ | 2.0 |
| **7-C19S-P**<br>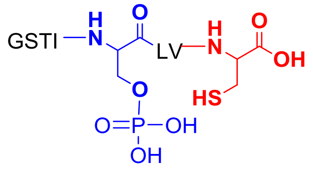 | 859.3631<br>$[M+H]^+$ | 859.3580<br>$[M+H]^+$ | 5.9 |
| **7-C19S-P-I**<br>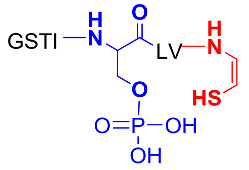 | 813.3576<br>$[M+H]^+$ | – – | – – |
| **7-C19S-P-II**<br>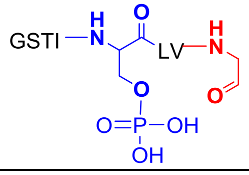 | 797.3805<br>$[M+H]^+$ | – – | – – |
| **7-C19S-Glu**<br>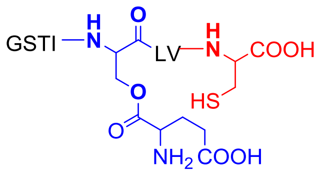 | 454.7233<br>$[M+2H]^{2+}$ | 454.7211<br>$[M+2H]^{2+}$ | 4.9 |
| **7-C19S-Glu-I**<br>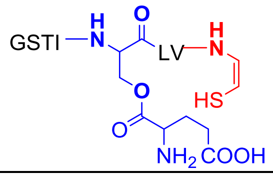 | 431.7206<br>$[M+2H]^{2+}$ | – – | – – |

| | | | |
|---|---|---|---|
| **7-C19S-Glu-II** 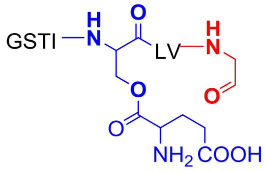 | 423.7320 $[M+2H]^{2+}$ | – – | – – |
| **7-Dha19** 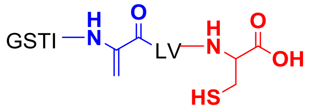 | 761.3862 $[M+H]^{+}$ | 761.3836 $[M+H]^{+}$ | 3.4 |
| **7-Dha19-I** 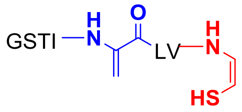 | 715.3808 $[M+H]^{+}$ | – – | – – |
| **7-Dha19-II** 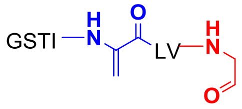 | 699.4036 $[M+H]^{+}$ | – – | – – |
| **7- d-C19** 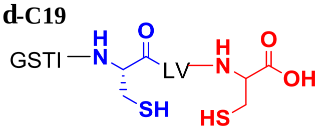 | 795.3739 $[M+H]^{+}$ | 795.3711 $[M+H]^{+}$ | 3.5 |
| **7-C19S-$D_3$** 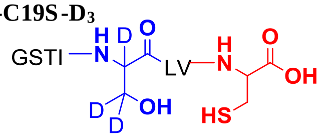 | 782.4156 $[M+H]^{+}$ | 782.4136 $[M+H]^{+}$ | 2.6 |
| **7-C19S-$D_3$-I** 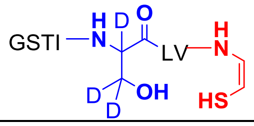 | 736.4101 $[M+H]^{+}$ | 736.4086 $[M+H]^{+}$ | 2.0 |
| **7-C19S-$D_3$-II** 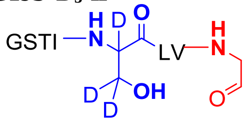 | 720.4330 $[M+H]^{+}$ | 720.4305 $[M+H]^{+}$ | 3.5 |

1

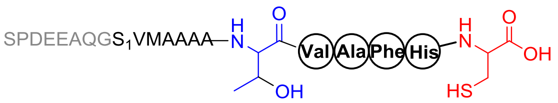


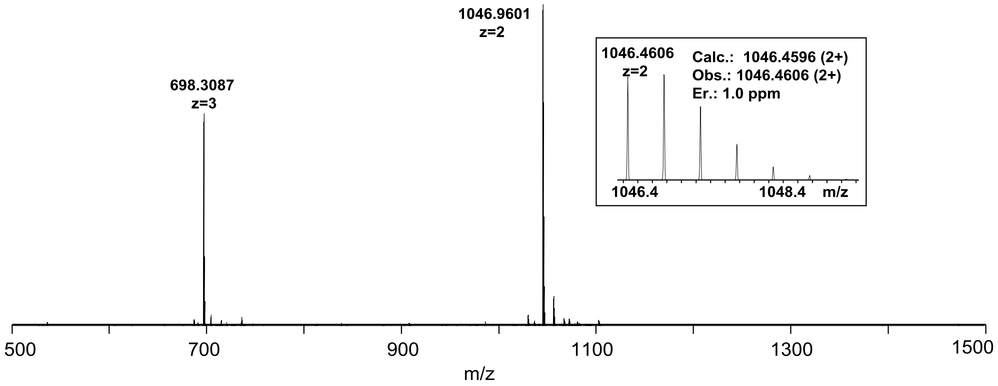


1

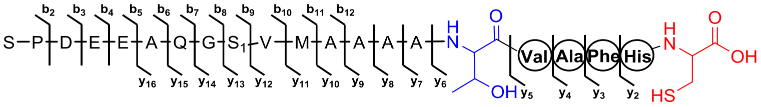


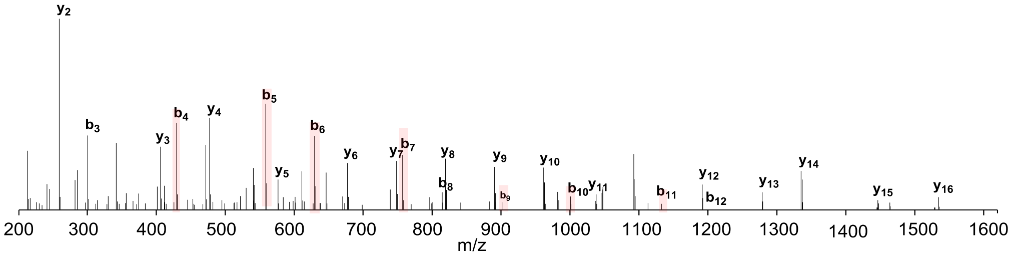


| Ions | Calcd. | Obs. | Er. (ppm) | Ions | Calcd. | Obs. | Er. (ppm) |
|---|---|---|---|---|---|---|---|
| **$b_2$** | 185.0926 | 185.0921 | 2.7 | **$y_2$** | 259.0864 | 259.0860 | 1.5 |
| **$b_3$** | 300.1196 | 300.1190 | 2.0 | **$y_3$** | 406.1548 | 406.1544 | 1.0 |
| **$b_4$** | 429.1621 | 429.1617 | 0.9 | **$y_4$** | 477.1920 | 477.1917 | 0.6 |
| **$b_5$** | 558.2047 | 558.2045 | 0.4 | **$y_5$** | 576.2604 | 576.2601 | 0.5 |
| **$b_6$** | 629.2418 | 629.2414 | 0.6 | **$y_6$** | 677.3081 | 677.3079 | 0.3 |
| **$b_7$** | 757.3004 | 757.3000 | 0.5 | **$y_7$** | 748.3452 | 748.3447 | 0.7 |
| **$b_8$** | 814.3219 | 814.3214 | 0.6 | **$y_8$** | 819.3823 | 819.3820 | 0.4 |
| **$b_9$** | 901.3539 | 901.3528 | 1.2 | **$y_9$** | 890.4194 | 890.4192 | 0.2 |
| **$b_{10}$** | 1000.4223 | 1000.4218 | 0.5 | **$y_{10}$** | 961.4565 | 961.4564 | 0.1 |
| **$b_{11}$** | 1131.4628 | 1131.4623 | 0.4 | **$y_{11}$** | 1092.4970 | 1092.4971 | 0.1 |
| **$b_{12}$** | 1202.4999 | 1202.4978 | 1.7 | **$y_{12}$** | 1191.5654 | 1191.5647 | 0.3 |
| | | | | **$y_{13}$** | 1278.5974 | 1278.5996 | 1.7 |
| | | | | **$y_{14}$** | 1335.6189 | 1335.6188 | 0.1 |
| | | | | **$y_{15}$** | 1463.6775 | 1463.6702 | 5.0 |
| | | | | **$y_{16}$** | 1534.7146 | 1534.7172 | 1.7 |

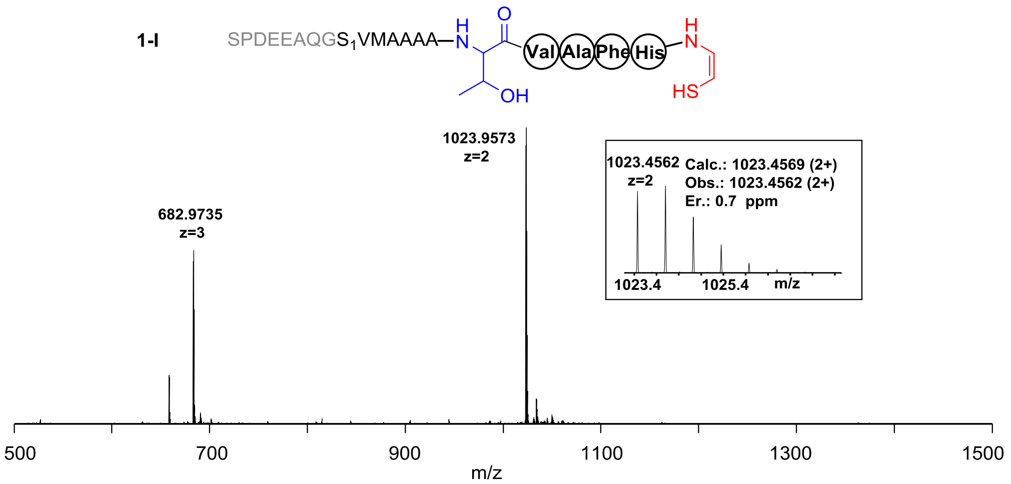


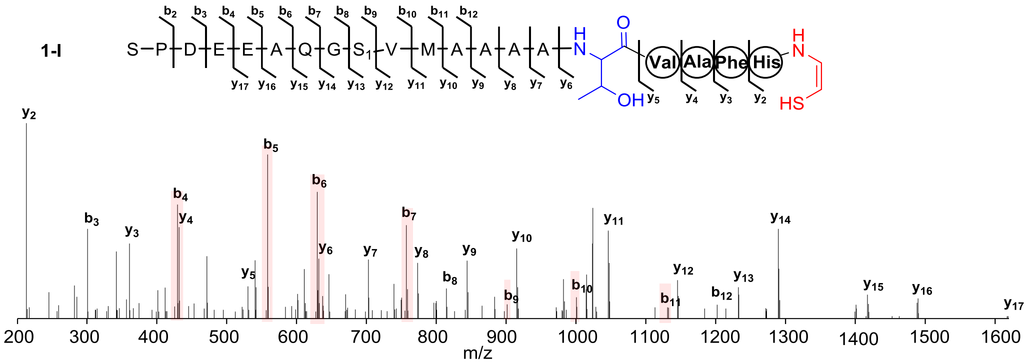


| Ions | Calcd. | Obs. | Er. (ppm) | Ions | Calcd. | Obs. | Er. (ppm) |
|---|---|---|---|---|---|---|---|
| $b_2$ | 185.0926 | 185.0921 | 2.7 | $y_2$ | 213.0810 | 213.0807 | 1.4 |
| $b_3$ | 300.1196 | 300.1189 | 2.3 | $y_3$ | 360.1494 | 360.1489 | 1.4 |
| $b_4$ | 429.1621 | 429.1618 | 0.7 | $y_4$ | 431.1865 | 431.1861 | 0.9 |
| $b_5$ | 558.2047 | 558.2045 | 0.4 | $y_5$ | 530.2549 | 530.2548 | 0.2 |
| $b_6$ | 629.2418 | 629.2413 | 0.8 | $y_6$ | 631.3026 | 631.3021 | 0.8 |
| $b_7$ | 757.3004 | 757.3003 | 0.1 | $y_7$ | 702.3397 | 702.3392 | 0.7 |
| $b_8$ | 814.3219 | 814.3215 | 0.5 | $y_8$ | 773.3768 | 773.3760 | 1.0 |
| $b_9$ | 901.3539 | 901.3558 | 2.1 | $y_9$ | 844.4139 | 844.4139 | 0.0 |
| $b_{10}$ | 1000.4223 | 1000.4229 | 0.6 | $y_{10}$ | 915.4510 | 915.4505 | 0.5 |
| $b_{11}$ | 1131.4628 | 1131.4623 | 0.4 | $y_{11}$ | 1046.4915 | 1046.4917 | 0.2 |
| $b_{12}$ | 1202.4999 | 1202.5009 | 0.8 | $y_{12}$ | 1145.5599 | 1145.5603 | 0.3 |
| | | | | $y_{13}$ | 1232.5920 | 1232.5917 | 0.2 |
| | | | | $y_{14}$ | 1289.6134 | 1289.6141 | 0.5 |
| | | | | $y_{15}$ | 1417.6720 | 1417.6727 | 0.5 |
| | | | | $y_{16}$ | 1488.7091 | 1488.7134 | 2.9 |
| | | | | $y_{17}$ | 1617.7517 | 1617.7543 | 1.6 |

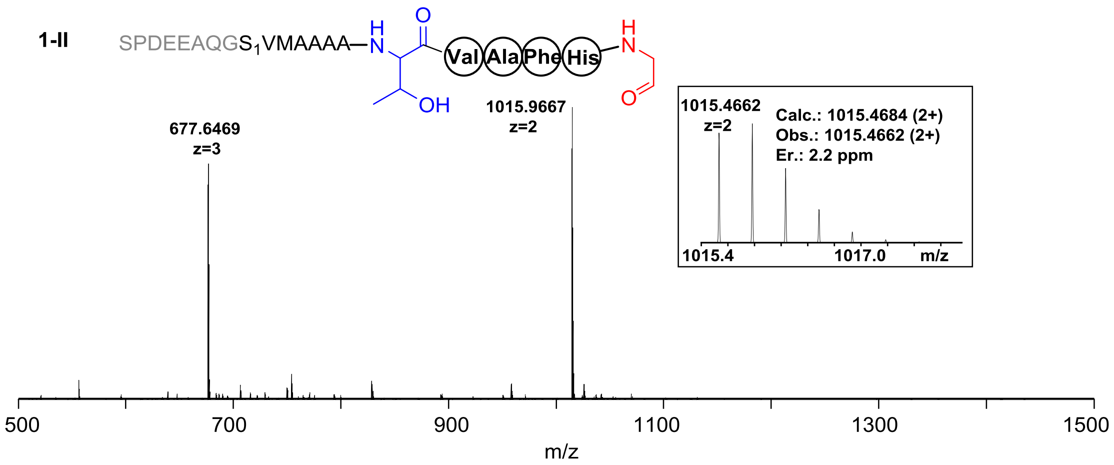


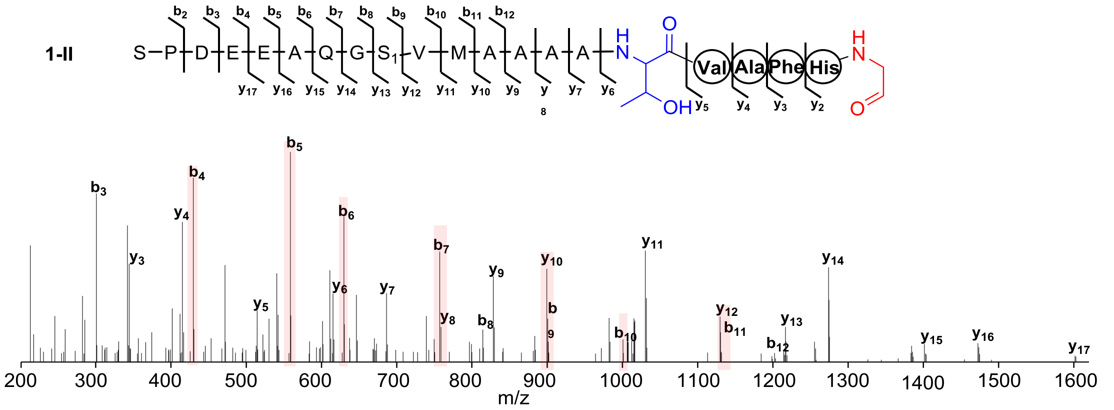


| Ions | Calcd. | Obs. | Er. (ppm) | Ions | Calcd. | Obs. | Er. (ppm) |
|---|---|---|---|---|---|---|---|
| $b_2$ | 185.0926 | 185.0921 | 2.7 | $y_2$ | 197.1038 | 197.1034 | 2.0 |
| $b_3$ | 300.1196 | 300.1190 | 2.0 | $y_3$ | 344.1722 | 344.1717 | 1.5 |
| $b_4$ | 429.1621 | 429.1617 | 0.9 | $y_4$ | 415.2093 | 415.2089 | 1.0 |
| $b_5$ | 558.2047 | 558.2043 | 0.7 | $y_5$ | 514.2777 | 514.2776 | 0.2 |
| $b_6$ | 629.2418 | 629.2415 | 0.5 | $y_6$ | 615.3254 | 615.3251 | 0.5 |
| $b_7$ | 757.3004 | 757.3001 | 0.4 | $y_7$ | 686.3625 | 686.3622 | 0.4 |
| $b_8$ | 814.3219 | 814.3215 | 0.5 | $y_8$ | 757.3996 | 757.3990 | 0.8 |
| $b_9$ | 901.3539 | 901.3535 | 0.4 | $y_9$ | 828.4368 | 828.4363 | 0.6 |
| $b_{10}$ | 1000.4223 | 1000.4216 | 0.7 | $y_{10}$ | 899.4739 | 899.4739 | 0.0 |
| $b_{11}$ | 1131.4628 | 1131.4618 | 0.9 | $y_{11}$ | 1030.5144 | 1030.5140 | 0.4 |
| $b_{12}$ | 1202.4999 | 1202.5031 | 2.7 | $y_{12}$ | 1129.5828 | 1129.5831 | 0.3 |
| | | | | $y_{13}$ | 1216.6148 | 1216.6149 | 0.1 |
| | | | | $y_{14}$ | 1273.6363 | 1273.6371 | 0.6 |
| | | | | $y_{15}$ | 1401.6948 | 1401.6958 | 0.7 |
| | | | | $y_{16}$ | 1472.7320 | 1472.7324 | 0.3 |
| | | | | $y_{17}$ | 1601.7746 | 1601.7646 | 6.2 |

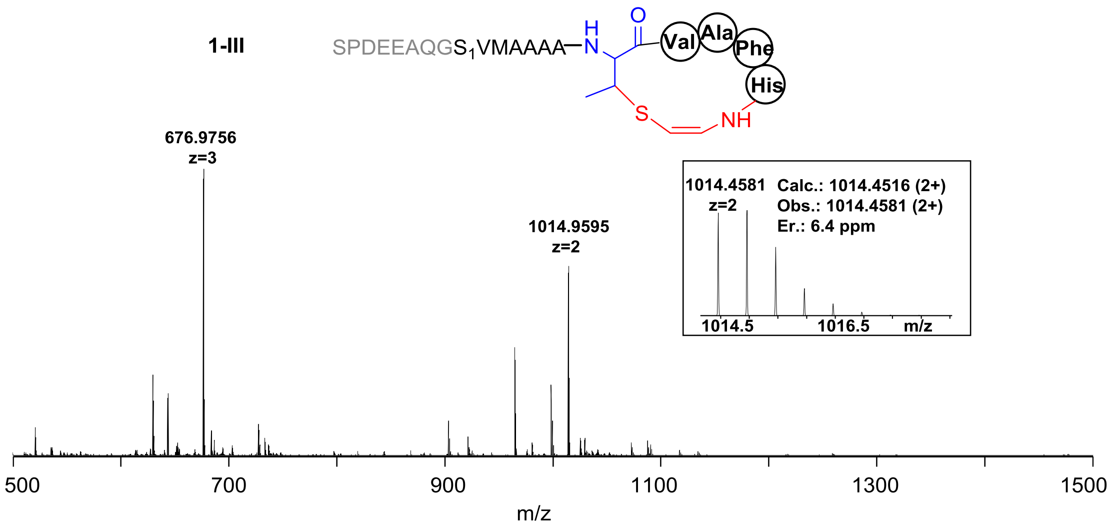


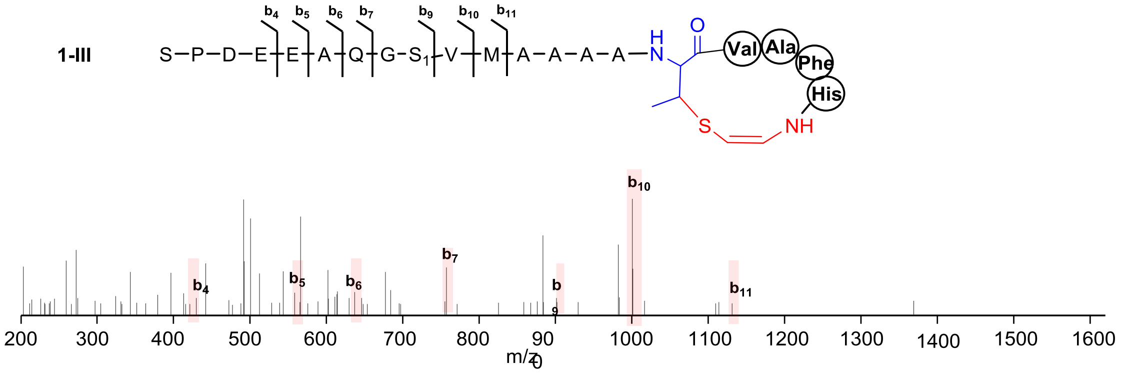


| Ions | Calcd. | Obs. | Er. (ppm) |
|---|---|---|---|
| **$b_4$** | 429.1621 | 429.1612 | 2.1 |
| **$b_5$** | 558.2047 | 558.2034 | 2.3 |
| **$b_6$** | 629.2418 | 629.2401 | 2.7 |
| **$b_7$** | 757.3004 | 757.2979 | 3.3 |
| **$b_9$** | 901.3539 | 901.3539 | 0.0 |
| **$b_{10}$** | 1000.4223 | 1000.4206 | 1.7 |
| **$b_{11}$** | 1131.4628 | 1131.4624 | 0.4 |

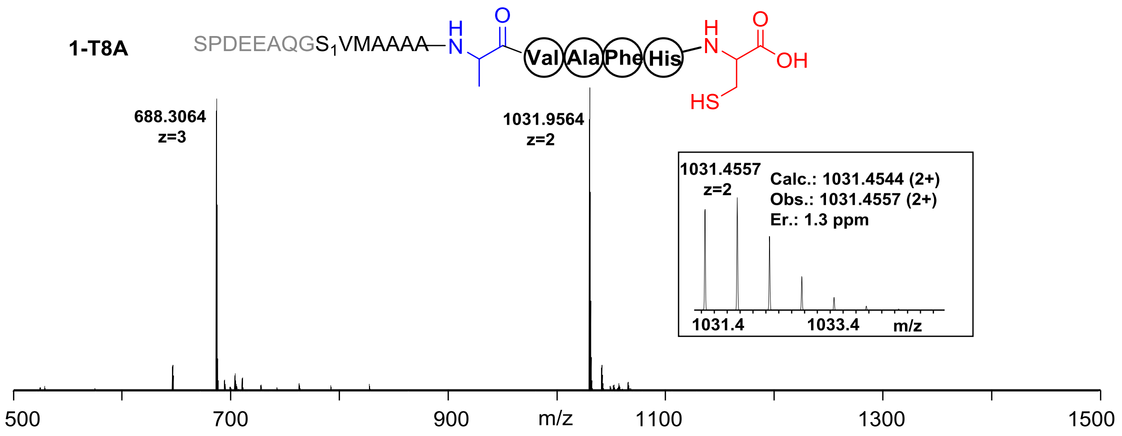


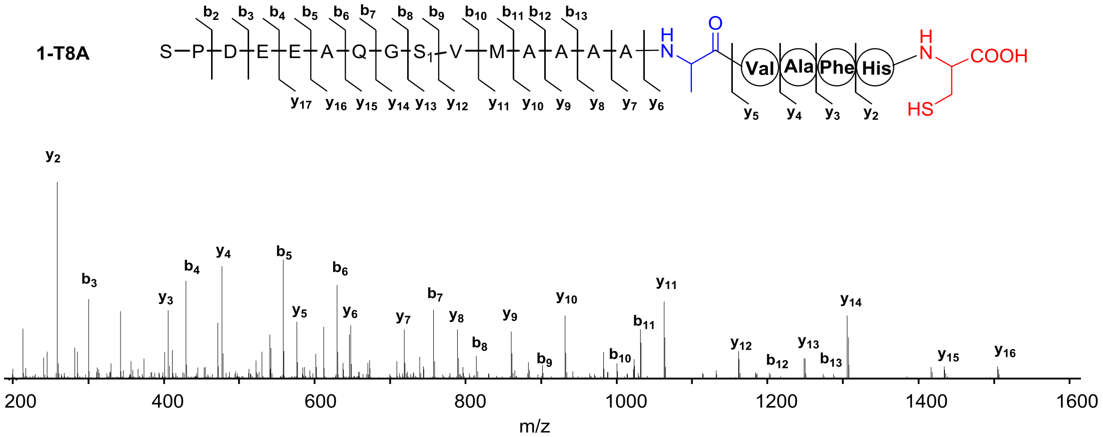


| Ions | Calcd. | Obs. | Er. (ppm) | Ions | Calcd. | Obs. | Er. (ppm) |
|---|---|---|---|---|---|---|---|
| **b2** | 185.0926 | 185.0925 | 0.5 | **y2** | 259.0864 | 259.0865 | 0.4 |
| **b3** | 300.1196 | 300.1196 | 0.0 | **y3** | 406.1548 | 406.1553 | 1.2 |
| **b4** | 429.1621 | 429.1627 | 1.4 | **y4** | 477.1920 | 477.1927 | 1.5 |
| **b5** | 558.2047 | 558.2057 | 1.8 | **y5** | 576.2604 | 576.2614 | 1.7 |
| **b6** | 629.2418 | 629.2428 | 1.6 | **y6** | 647.2975 | 647.2982 | 1.1 |
| **b7** | 757.3004 | 757.3016 | 1.6 | **y7** | 718.3346 | 718.3358 | 1.7 |
| **b8** | 814.3219 | 814.3230 | 1.4 | **y8** | 789.3717 | 789.3732 | 1.9 |
| **b9** | 901.3539 | 901.3554 | 1.7 | **y9** | 860.4088 | 860.4099 | 1.3 |
| **b10** | 1000.4223 | 1000.4249 | 2.6 | **y10** | 931.4459 | 931.4471 | 1.3 |
| **b11** | 1131.4628 | 1131.4663 | 3.1 | **y11** | 1062.4864 | 1062.4883 | 1.8 |
| **b12** | 1202.4999 | 1202.4990 | 0.7 | **y12** | 1161.5548 | 1161.5579 | 2.7 |
| **b13** | 1273.5371 | 1273.5366 | 0.4 | **y13** | 1248.5869 | 1248.5912 | 3.4 |
| | | | | **y14** | 1305.6083 | 1305.6116 | 2.5 |
| | | | | **y15** | 1433.6669 | 1433.6674 | 0.3 |
| | | | | **y16** | 1504.7040 | 1504.7078 | 2.5 |
| | | | | **y17** | 1633.7466 | 1633.7512 | 2.8 |

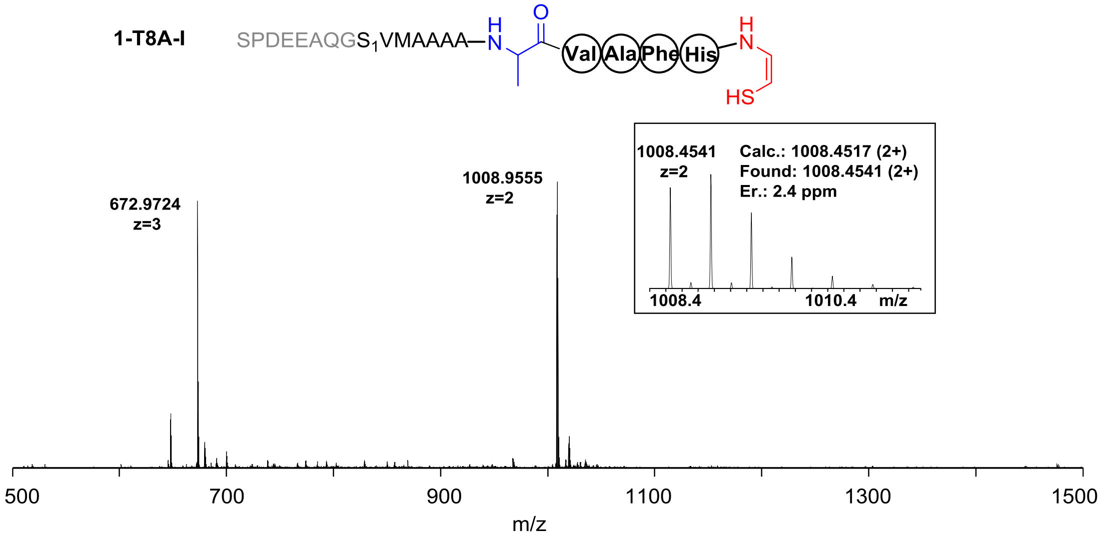


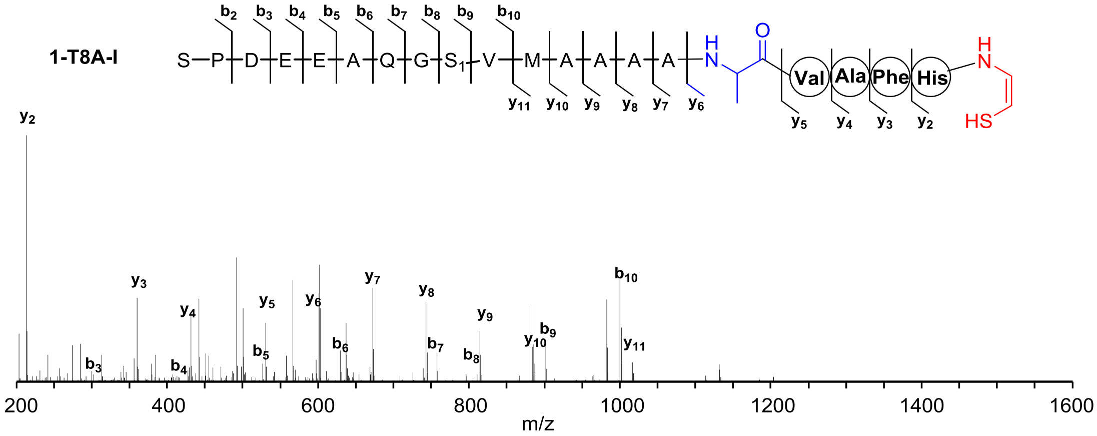


| Ions | Calcd. | Obs. | Er. (ppm) | Ions | Calcd. | Obs. | Er. (ppm) |
|---|---|---|---|---|---|---|---|
| $b_2$ | 185.0926 | 185.0924 | 1.1 | $y_2$ | 213.0810 | 213.0809 | 0.5 |
| $b_3$ | 300.1196 | 300.1194 | 0.7 | $y_3$ | 360.1494 | 360.1495 | 0.3 |
| $b_4$ | 429.1621 | 429.1620 | 0.2 | $y_4$ | 431.1865 | 431.1869 | 0.9 |
| $b_5$ | 558.2047 | 558.2054 | 1.3 | $y_5$ | 530.2549 | 530.2557 | 1.5 |
| $b_6$ | 629.2418 | 629.2429 | 1.7 | $y_6$ | 601.2920 | 601.2928 | 1.3 |
| $b_7$ | 757.3004 | 757.3011 | 0.9 | $y_7$ | 672.3291 | 672.3300 | 1.3 |
| $b_8$ | 814.3219 | 814.3228 | 1.1 | $y_8$ | 743.3662 | 743.3671 | 1.2 |
| $b_9$ | 901.3539 | 901.3548 | 1.0 | $y_9$ | 814.4034 | 814.4041 | 0.9 |
| $b_{10}$ | 1000.4223 | 1000.4239 | 1.6 | $y_{10}$ | 885.4405 | 885.4415 | 1.1 |
| | | | | $y_{11}$ | 1016.4810 | 1016.4821 | 1.1 |

**1-T8A-II**

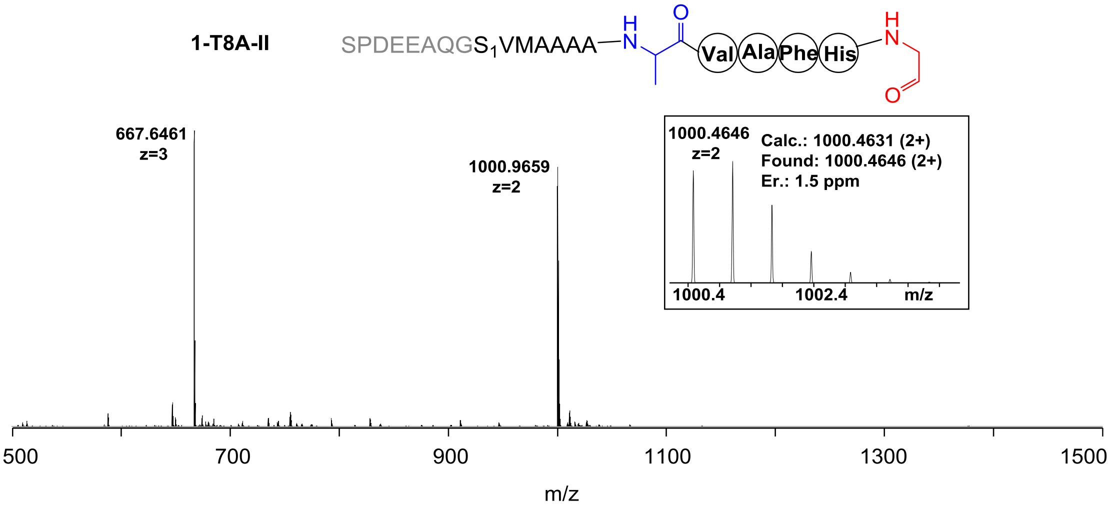
SPDEEAQGS1VMAAAA
H
N
O
Val
Ala
Phe
His
H
N
O
667.6461
z=3
1000.9659
z=2
1000.4646
z=2
Calc.: 1000.4631 (2+)
Found: 1000.4646 (2+)
Er.: 1.5 ppm
1000.4
1002.4
m/z
500
700
900
1100
1300
1500
m/z

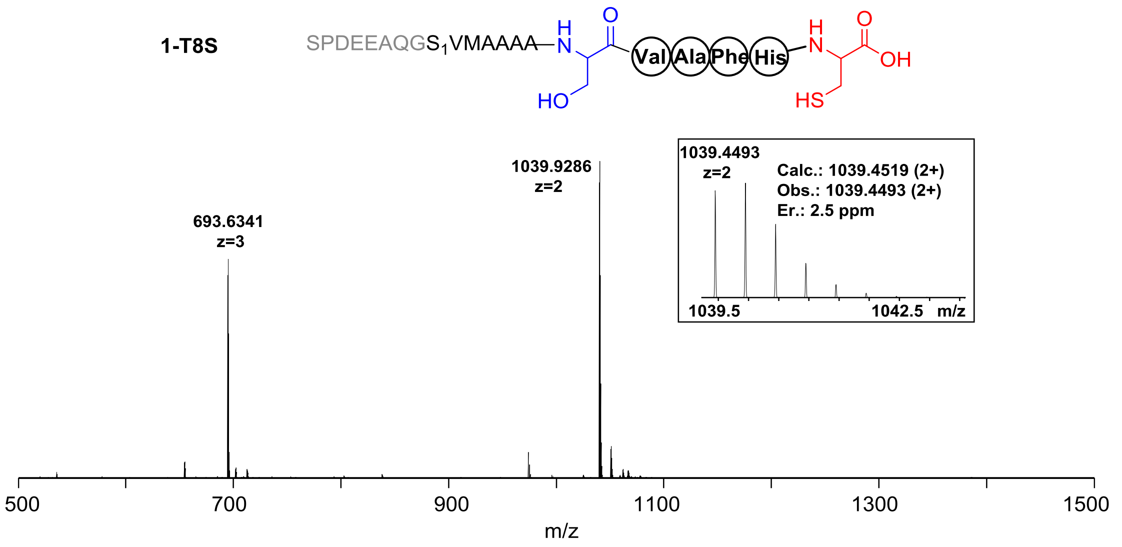


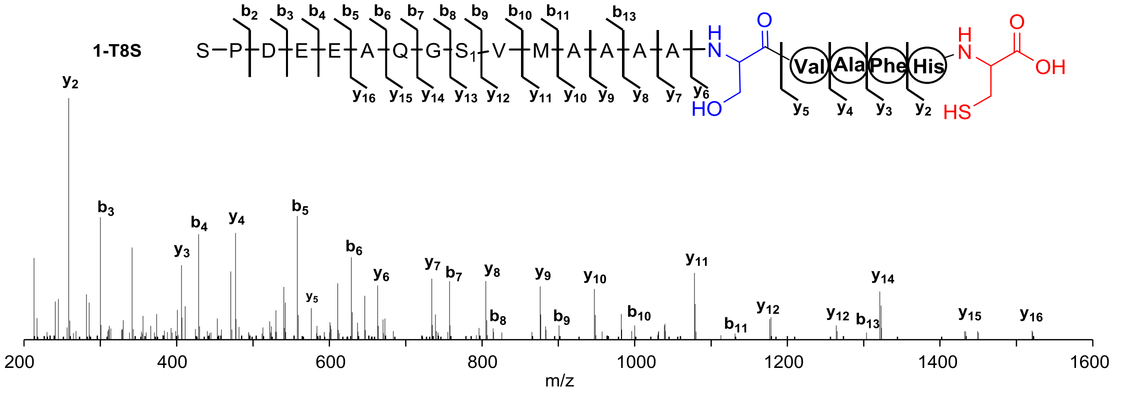


| Ions | Calcd. | Obs. | Er. (ppm) | Ions | Calcd. | Obs. | Er. (ppm) |
|---|---|---|---|---|---|---|---|
| $b_2$ | 185.0926 | 185.0918 | 4.3 | $y_2$ | 259.0864 | 259.0855 | 3.5 |
| $b_3$ | 300.1196 | 300.1185 | 3.7 | $y_3$ | 406.1548 | 406.1536 | 3.0 |
| $b_4$ | 429.1621 | 429.1608 | 3.0 | $y_4$ | 477.1920 | 477.1907 | 2.7 |
| $b_5$ | 558.2047 | 558.2032 | 2.7 | $y_5$ | 576.2604 | 576.2590 | 2.4 |
| $b_6$ | 629.2418 | 629.2401 | 2.7 | $y_6$ | 663.2924 | 663.2906 | 2.7 |
| $b_7$ | 757.3004 | 757.2986 | 2.4 | $y_7$ | 734.3295 | 734.3281 | 1.9 |
| $b_8$ | 814.3219 | 814.3184 | 4.3 | $y_8$ | 805.3666 | 805.3644 | 2.7 |
| $b_9$ | 901.3539 | 901.3513 | 2.9 | $y_9$ | 876.4037 | 876.4011 | 3.0 |
| $b_{10}$ | 1000.4223 | 1000.4188 | 3.5 | $y_{10}$ | 947.4409 | 947.4387 | 2.3 |
| $b_{11}$ | 1131.4628 | 1131.4606 | 1.9 | $y_{11}$ | 1078.4813 | 1078.4789 | 2.2 |
| $b_{13}$ | 1273.5371 | 1273.5305 | 5.2 | $y_{12}$ | 1177.5498 | 1177.5486 | 1.0 |
| | | | | $y_{13}$ | 1264.5818 | 1264.5795 | 1.8 |
| | | | | $y_{14}$ | 1321.6033 | 1321.6012 | 1.6 |
| | | | | $y_{15}$ | 1449.6618 | 1449.6553 | 4.5 |
| | | | | $y_{16}$ | 1520.6989 | 1520.6921 | 4.5 |

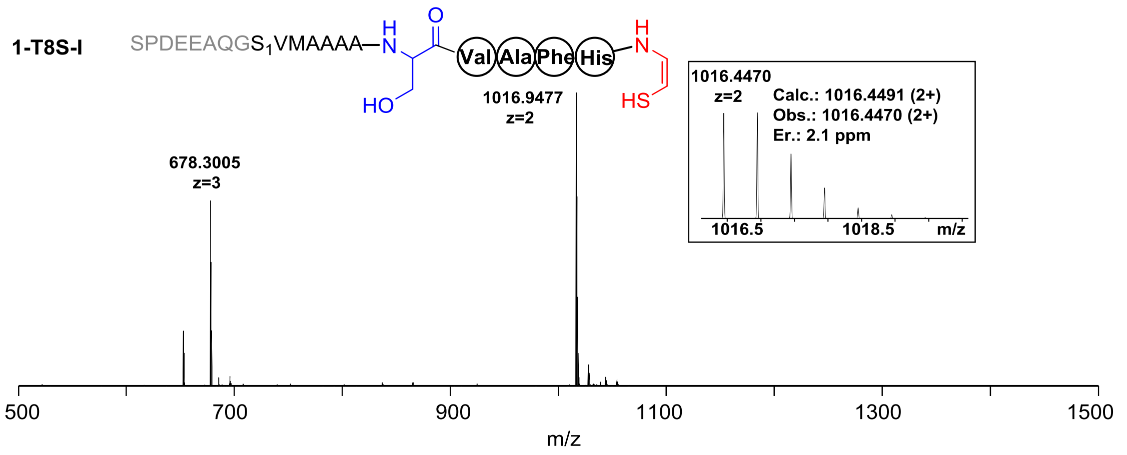


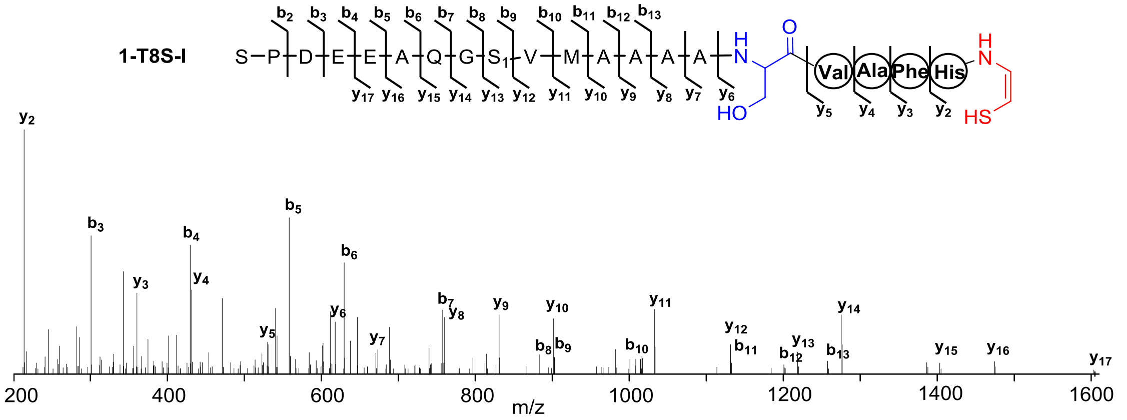


| Ions | Calcd. | Obs. | Er. (ppm) | Ions | Calcd. | Obs. | Er. (ppm) |
|---|---|---|---|---|---|---|---|
| $b_2$ | 185.0926 | 185.0917 | 4.9 | $y_2$ | 213.0810 | 213.0800 | 4.7 |
| $b_3$ | 300.1196 | 300.1182 | 4.7 | $y_3$ | 360.1494 | 360.1481 | 3.6 |
| $b_4$ | 429.1621 | 429.1607 | 3.3 | $y_4$ | 431.1865 | 431.1850 | 3.5 |
| $b_5$ | 558.2047 | 558.2030 | 3.0 | $y_5$ | 530.2549 | 530.2532 | 3.2 |
| $b_6$ | 629.2418 | 629.2397 | 3.3 | $y_6$ | 617.2869 | 617.2850 | 3.1 |
| $b_7$ | 757.3004 | 757.2979 | 3.3 | $y_7$ | 688.3240 | 688.3217 | 3.3 |
| $b_8$ | 814.3219 | 814.3190 | 3.6 | $y_8$ | 759.3612 | 759.3586 | 3.4 |
| $b_9$ | 901.3539 | 901.3532 | 0.8 | $y_9$ | 830.3983 | 830.3956 | 3.3 |
| $b_{10}$ | 1000.4223 | 1000.4202 | 2.1 | $y_{10}$ | 901.4354 | 901.4319 | 3.9 |
| $b_{11}$ | 1131.4628 | 1131.4576 | 4.6 | $y_{11}$ | 1032.4759 | 1032.4729 | 2.9 |
| $b_{12}$ | 1202.4999 | 1202.4965 | 2.8 | $y_{12}$ | 1131.5443 | 1131.5414 | 2.6 |
| $b_{13}$ | 1273.5371 | 1273.5314 | 4.8 | $y_{13}$ | 1218.5763 | 1218.5739 | 2.0 |
| | | | | $y_{14}$ | 1275.5978 | 1275.5944 | 2.7 |
| | | | | $y_{15}$ | 1403.6564 | 1403.6528 | 2.6 |
| | | | | $y_{16}$ | 1474.6935 | 1474.6876 | 4.0 |
| | | | | $y_{17}$ | 1603.7361 | 1603.7247 | 7.1 |

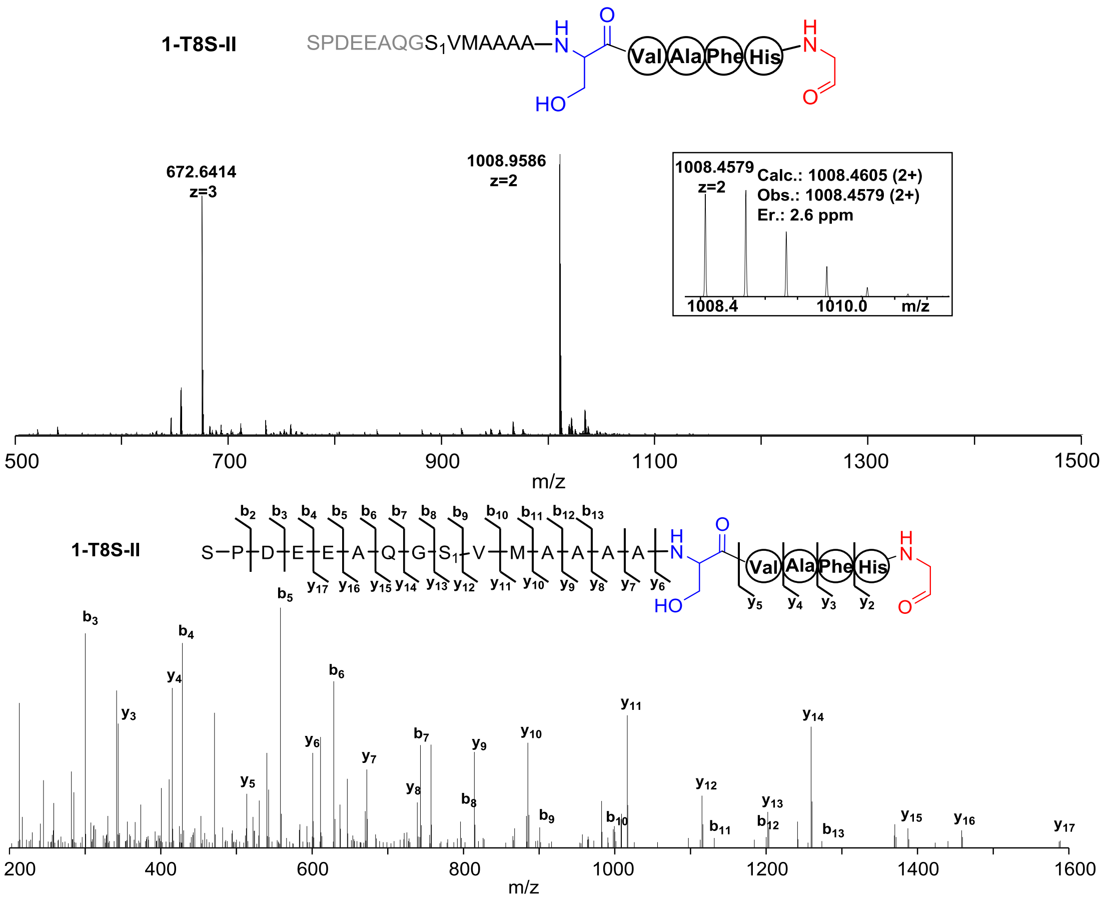


| Ions | Calcd. | Obs. | Er. (ppm) | Ions | Calcd. | Obs. | Er. (ppm) |
|---|---|---|---|---|---|---|---|
| $b_2$ | 185.0926 | 185.0917 | 4.9 | $y_2$ | 197.1038 | 197.1029 | 4.6 |
| $b_3$ | 300.1196 | 300.1183 | 4.3 | $y_3$ | 344.1722 | 344.1709 | 3.8 |
| $b_4$ | 429.1621 | 429.1607 | 3.3 | $y_4$ | 415.2093 | 415.2079 | 3.4 |
| $b_5$ | 558.2047 | 558.2031 | 2.9 | $y_5$ | 514.2777 | 514.2762 | 2.9 |
| $b_6$ | 629.2418 | 629.2399 | 3.0 | $y_6$ | 601.3098 | 601.3080 | 3.0 |
| $b_7$ | 757.3004 | 757.2979 | 3.3 | $y_7$ | 672.3469 | 672.3448 | 3.1 |
| $b_8$ | 814.3219 | 814.3194 | 3.1 | $y_8$ | 743.3840 | 743.3818 | 3.0 |
| $b_9$ | 901.3539 | 901.3522 | 1.9 | $y_9$ | 814.4211 | 814.4185 | 3.2 |
| $b_{10}$ | 1000.4223 | 1000.4153 | 7.0 | $y_{10}$ | 885.4582 | 885.4559 | 2.6 |
| $b_{11}$ | 1131.4628 | 1131.4601 | 2.4 | $y_{11}$ | 1016.4987 | 1016.4959 | 2.6 |
| $b_{12}$ | 1202.4999 | 1202.4944 | 4.6 | $y_{12}$ | 1115.5671 | 1115.5648 | 2.1 |
| $b_{13}$ | 1273.5371 | 1273.5309 | 4.9 | $y_{13}$ | 1202.5992 | 1202.5963 | 2.4 |
| | | | | $y_{14}$ | 1259.6206 | 1259.6176 | 2.4 |
| | | | | $y_{15}$ | 1387.6792 | 1387.6765 | 1.9 |
| | | | | $y_{16}$ | 1458.7163 | 1458.7126 | 2.5 |
| | | | | $y_{17}$ | 1587.7589 | 1587.7516 | 4.6 |

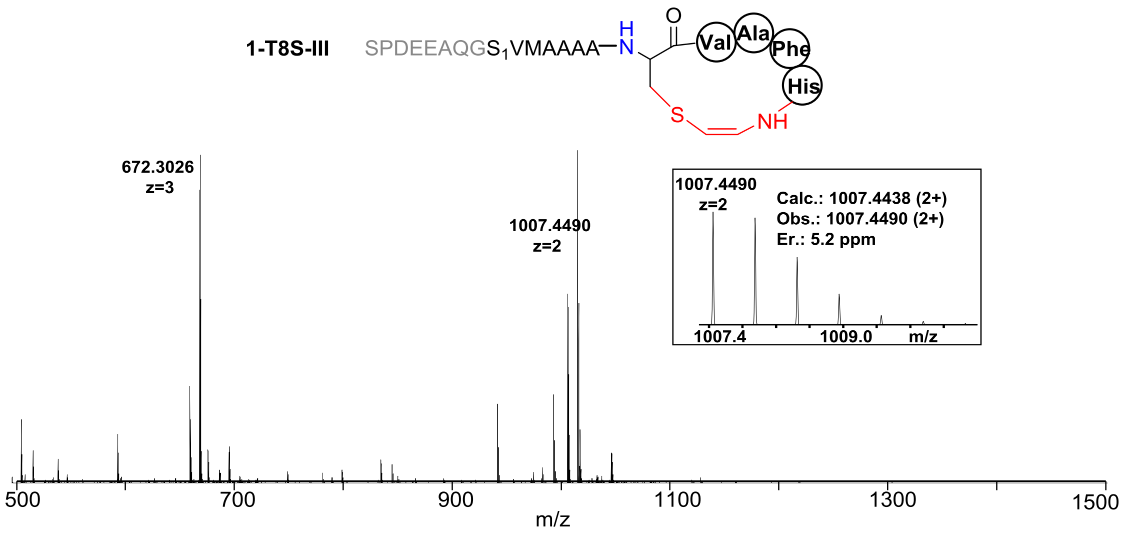


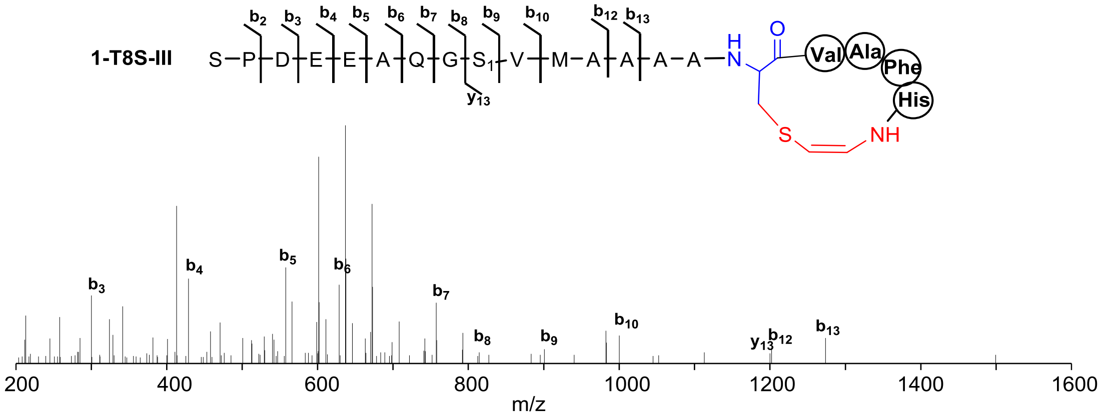


| Ions | Calcd. | Obs. | Er. (ppm) |
|---|---|---|---|
| **$b_2$** | 185.0926 | 185.0917 | 4.9 |
| **$b_3$** | 300.1196 | 300.1182 | 4.7 |
| **$b_4$** | 429.1621 | 429.1604 | 3.9 |
| **$b_5$** | 558.2047 | 558.2034 | 2.3 |
| **$b_6$** | 629.2418 | 629.2399 | 3.0 |
| **$b_7$** | 757.3004 | 757.2980 | 3.2 |
| **$b_8$** | 814.3219 | 814.3208 | 1.4 |
| **$b_9$** | 901.3539 | 901.3511 | 3.1 |
| **$b_{10}$** | 1000.4223 | 1000.4196 | 2.7 |
| **$b_{12}$** | 1202.4999 | 1202.4965 | 2.8 |
| **$b_{13}$** | 1273.5371 | 1273.5314 | 4.5 |
| **$y_{13}$** | 1200.5657 | 1200.5712 | 4.6 |

**1-T8C**

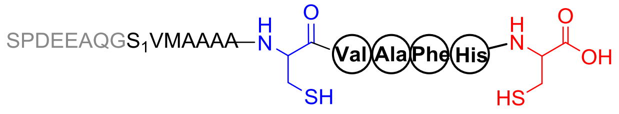
SPDEEAQGS1VMAAAA
Val
Ala
Phe
His
SH
HS
OH

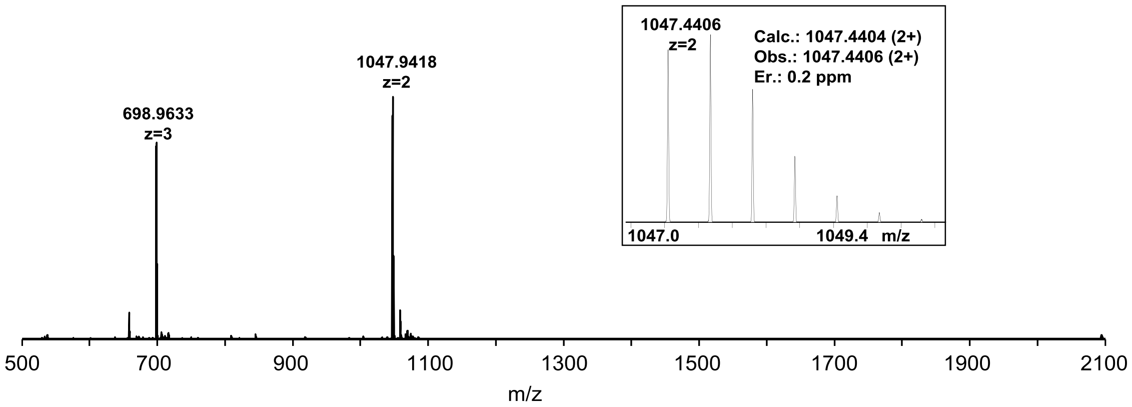
1047.9418
z=2
698.9633
z=3
1047.4406
z=2
Calc.: 1047.4404 (2+)
Obs.: 1047.4406 (2+)
Er.: 0.2 ppm
1047.0
1049.4 m/z
m/z

**1-T8C-II**

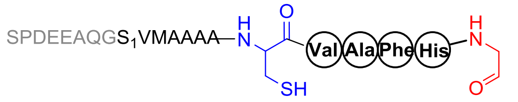
SPDEEAQGS1VMAAAA
Val
Ala
Phe
His
SH

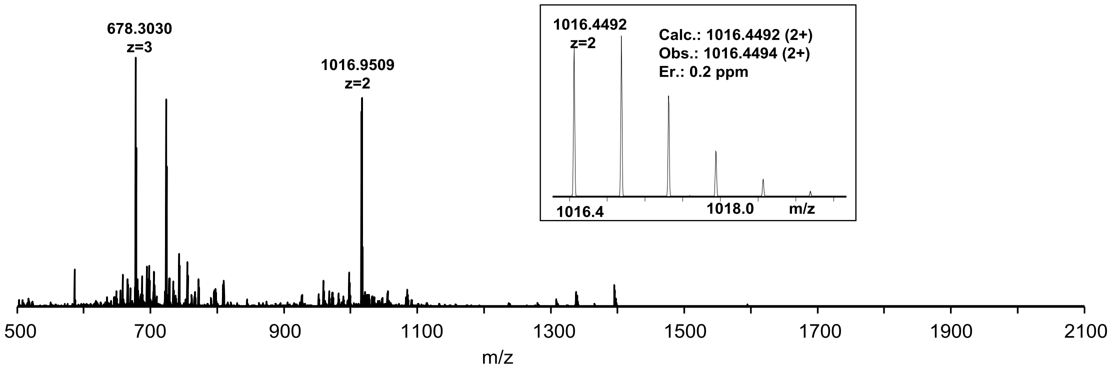
678.3030
z=3
1016.9509
z=2
1016.4492
z=2
Calc.: 1016.4492 (2+)
Obs.: 1016.4494 (2+)
Er.: 0.2 ppm
1016.4
1018.0
m/z
m/z

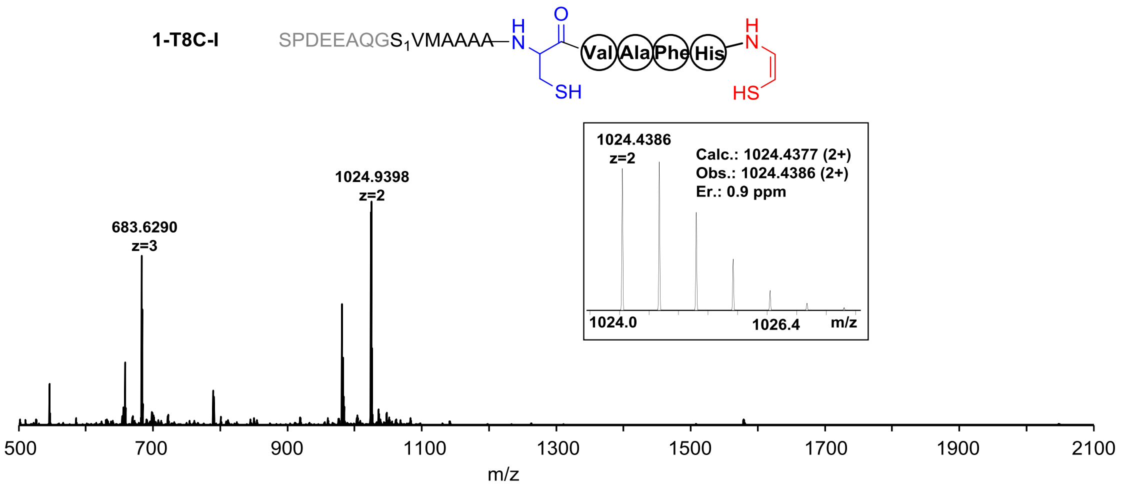


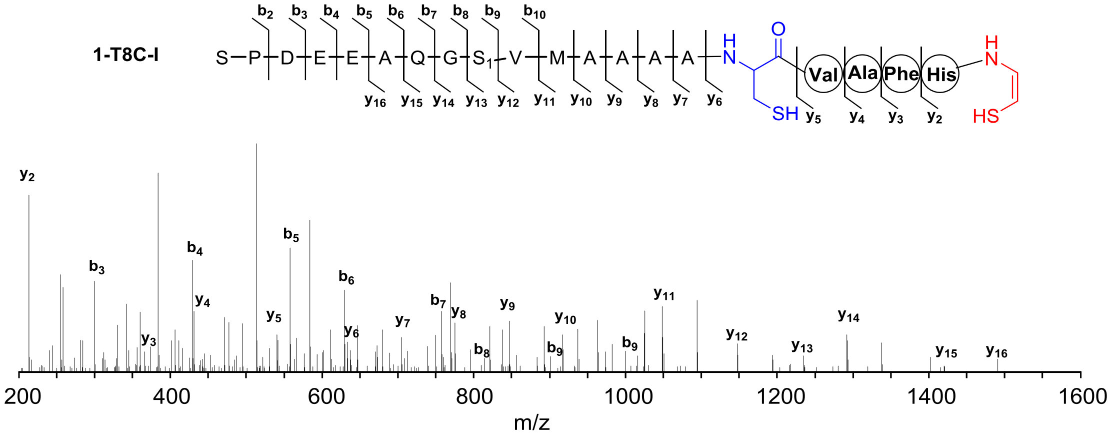


| Ions | Calcd. | Obs. | Er. (ppm) | Ions | Calcd. | Obs. | Er. (ppm) |
|---|---|---|---|---|---|---|---|
| $b_2$ | 185.0926 | 185.0924 | 1.1 | $y_2$ | 213.0810 | 213.0808 | 0.9 |
| $b_3$ | 300.1196 | 300.1195 | 0.3 | $y_3$ | 360.1494 | 360.1494 | 0.0 |
| $b_4$ | 429.1621 | 429.1626 | 1.2 | $y_4$ | 431.1865 | 431.1870 | 1.2 |
| $b_5$ | 558.2047 | 558.2055 | 1.4 | $y_5$ | 530.2549 | 530.2564 | 2.8 |
| $b_6$ | 629.2418 | 629.2422 | 0.6 | $y_6$ | 633.2641 | 633.2654 | 2.1 |
| $b_7$ | 757.3004 | 757.3005 | 0.1 | $y_7$ | 704.3012 | 704.3013 | 0.1 |
| $b_8$ | 814.3219 | 814.3226 | 0.9 | $y_8$ | 775.3383 | 775.3394 | 1.4 |
| $b_9$ | 901.3539 | 901.3531 | 0.9 | $y_9$ | 846.3754 | 846.3764 | 1.2 |
| $b_{10}$ | 1000.4223 | 1000.4243 | 2.0 | $y_{10}$ | 917.4125 | 917.4136 | 1.2 |
| | | | | $y_{11}$ | 1048.4530 | 1048.4552 | 2.1 |
| | | | | $y_{12}$ | 1147.5214 | 1147.5211 | 0.3 |
| | | | | $y_{13}$ | 1234.5535 | 1234.5518 | 1.4 |
| | | | | $y_{14}$ | 1291.5749 | 1291.5773 | 1.9 |
| | | | | $y_{15}$ | 1419.6335 | 1419.6300 | 2.5 |
| | | | | $y_{16}$ | 1490.6706 | 1490.6726 | 1.3 |

**1-C13A**

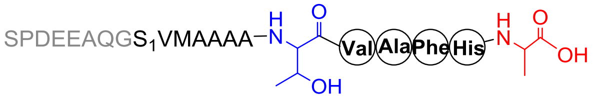


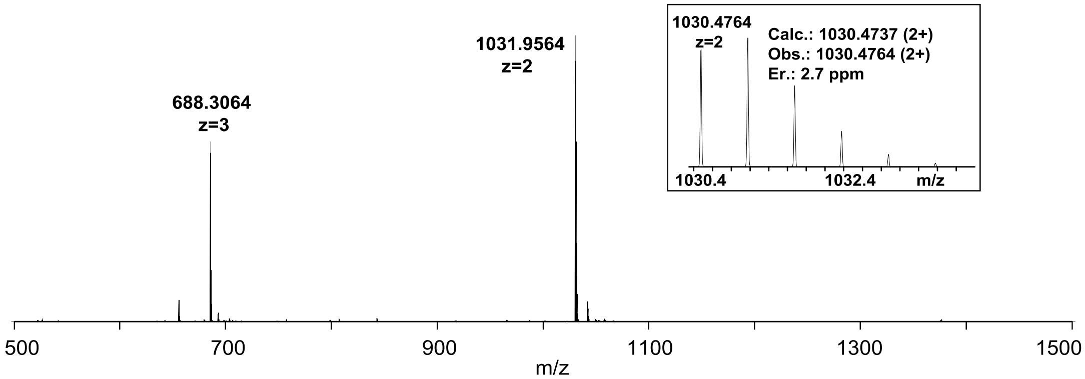


**1-C13S**

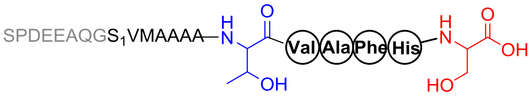


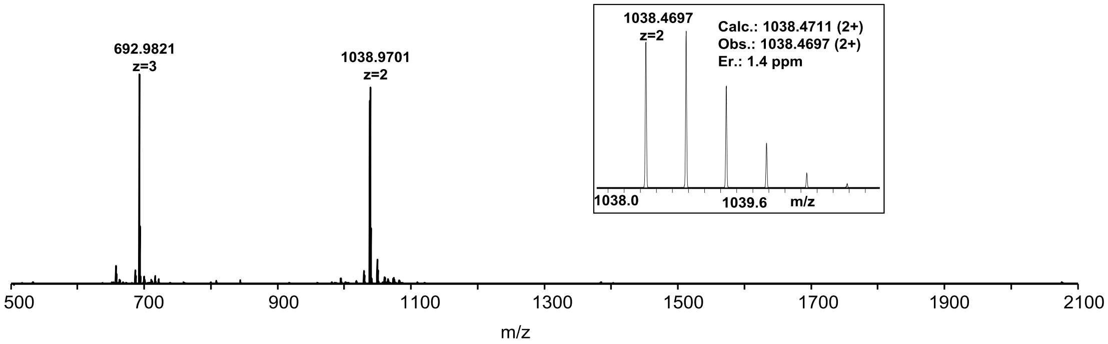


**1-C13T**

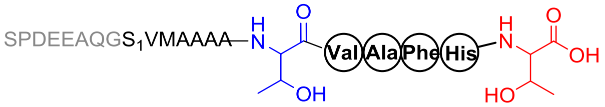


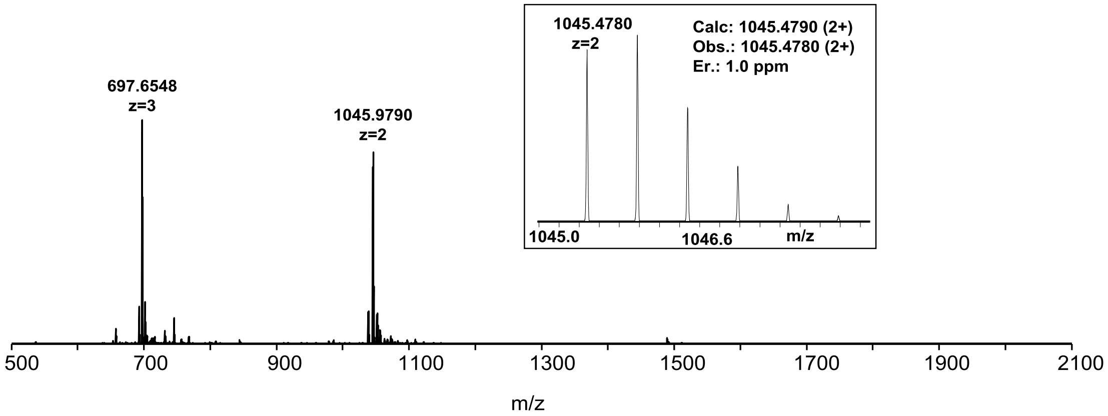

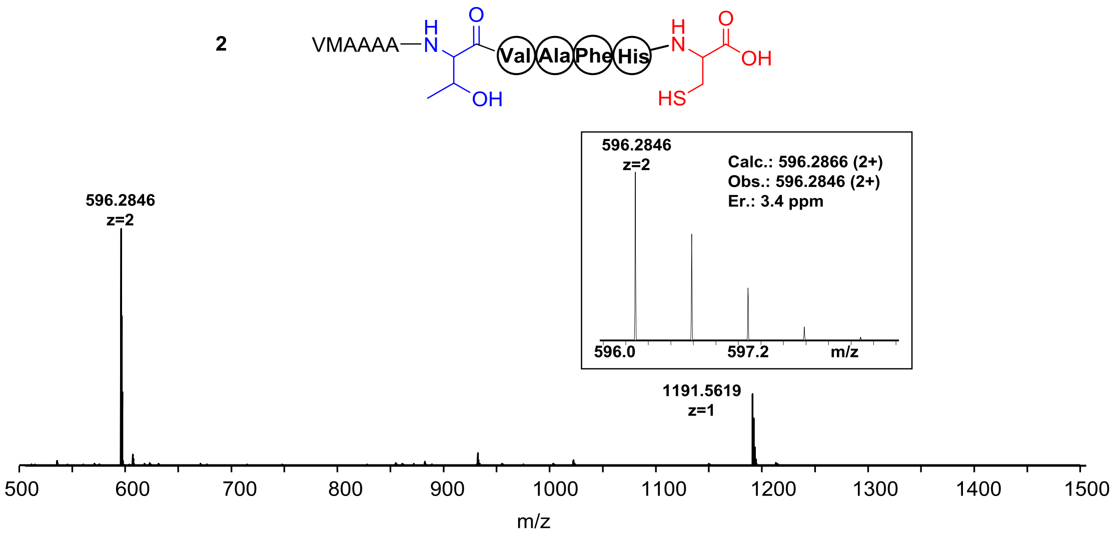
2
VMAAAA
Val
Ala
Phe
His
OH
HS
596.2846
z=2
Calc.: 596.2866 (2+)
Obs.: 596.2846 (2+)
Er.: 3.4 ppm
596.0
597.2
m/z
1191.5619
z=1
500
600
700
800
900
1000
1100
1200
1300
1400
1500
m/z

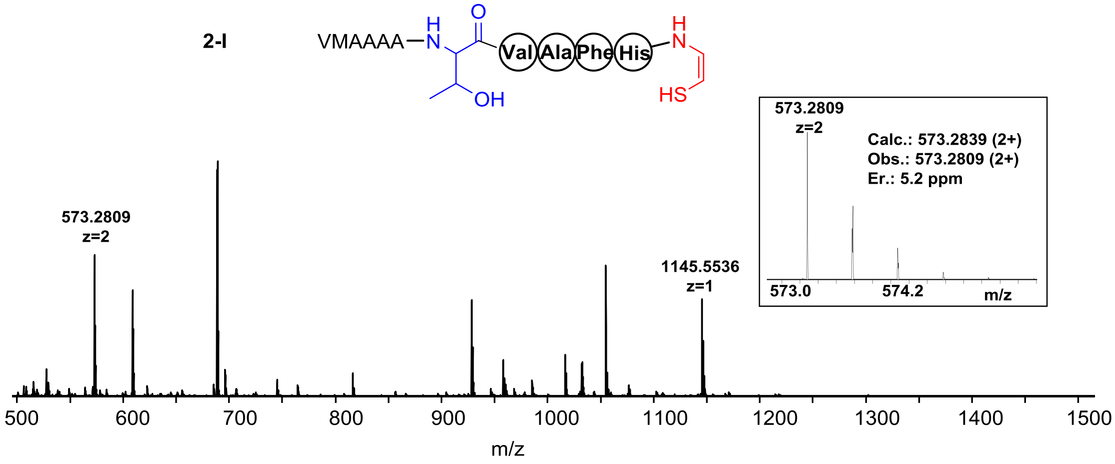
2-I
VMAAAA
Val
Ala
Phe
His
OH
HS
573.2809
z=2
Calc.: 573.2839 (2+)
Obs.: 573.2809 (2+)
Er.: 5.2 ppm
573.0
574.2
m/z
1145.5536
z=1
500
600
700
800
900
1000
1100
1200
1300
1400
1500
m/z

**2-III**

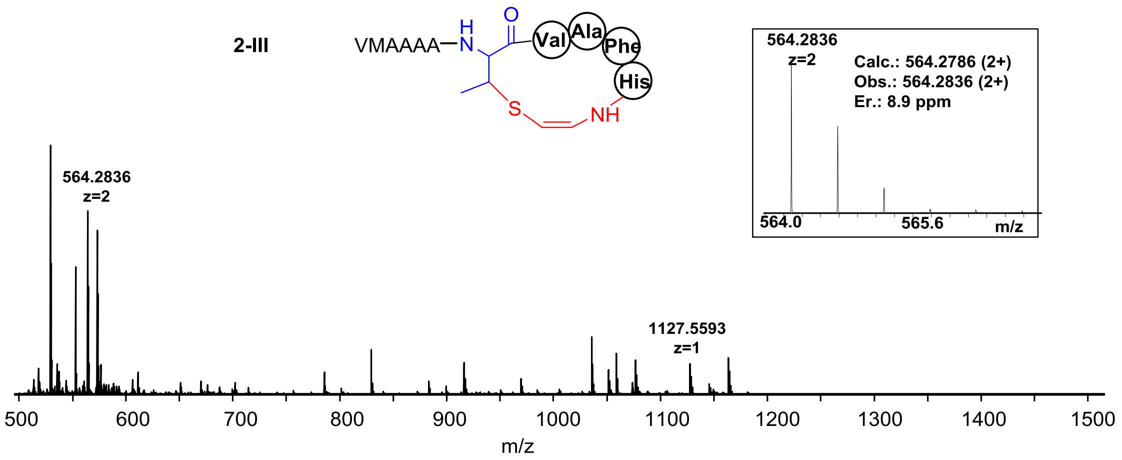
VMAAAA
H
N
O
Val
Ala
Phe
His
S
NH
564.2836
z=2
Calc.: 564.2786 (2+)
Obs.: 564.2836 (2+)
Er.: 8.9 ppm
564.0
565.6
m/z
564.2836
z=2
1127.5593
z=1
500
600
700
800
900
1000
1100
1200
1300
1400
1500
m/z

**2-II**

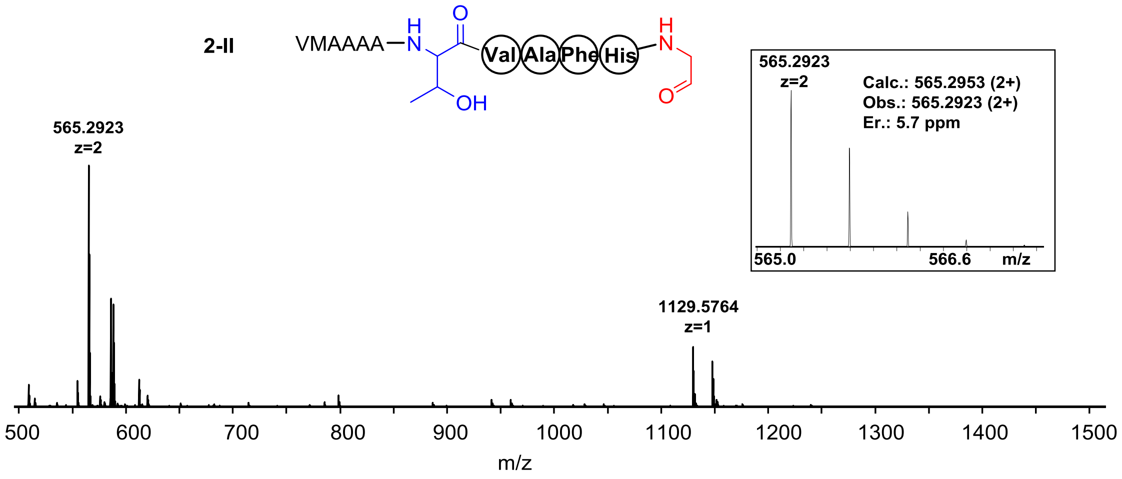
VMAAAA
H
N
O
Val
Ala
Phe
His
H
N
OH
O
565.2923
z=2
Calc.: 565.2953 (2+)
Obs.: 565.2923 (2+)
Er.: 5.7 ppm
565.0
566.6
m/z
565.2923
z=2
1129.5764
z=1
500
600
700
800
900
1000
1100
1200
1300
1400
1500
m/z

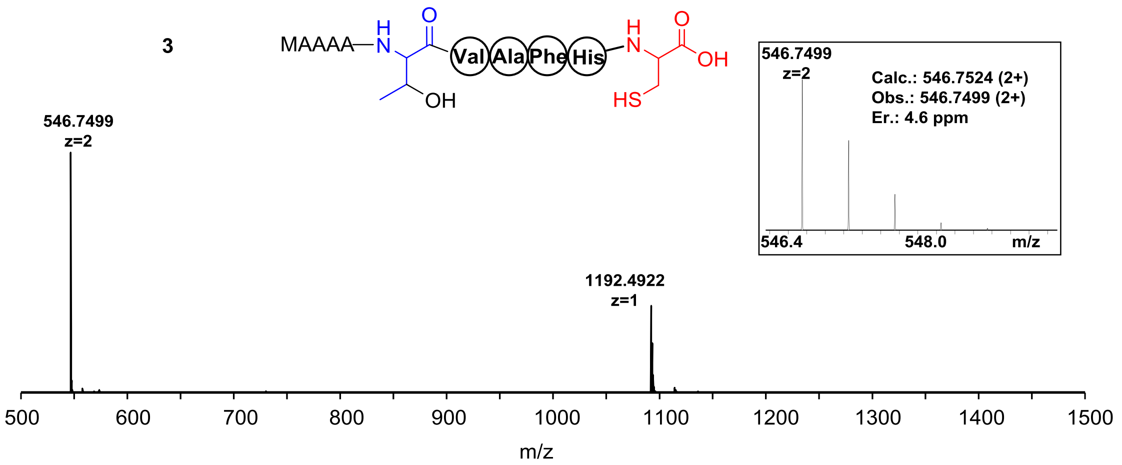
3
MAAAA
Val Ala Phe His
OH
HS
546.7499
z=2
1192.4922
z=1
Calc.: 546.7524 (2+)
Obs.: 546.7499 (2+)
Er.: 4.6 ppm
546.4
548.0
m/z
500
600
700
800
900
1000
1100
1200
1300
1400
1500
m/z

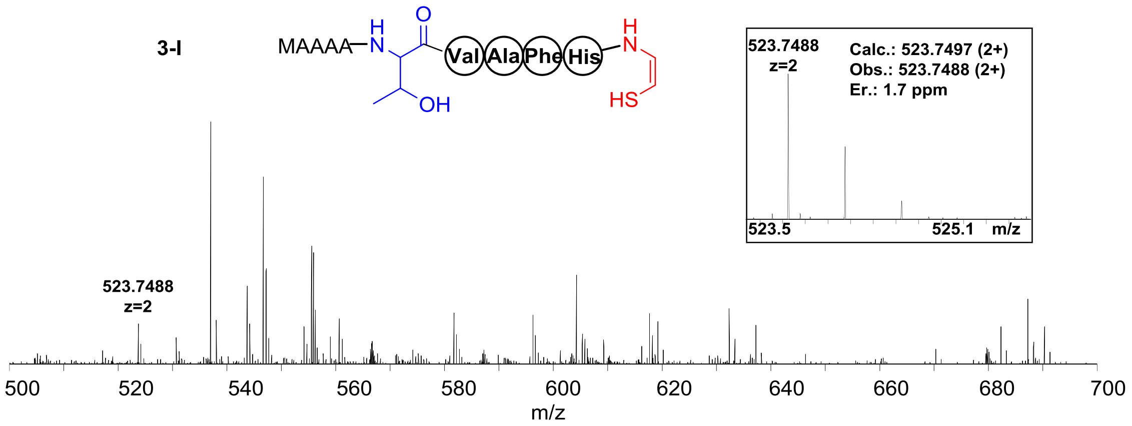
3-I
MAAAA
Val Ala Phe His
OH
HS
523.7488
z=2
Calc.: 523.7497 (2+)
Obs.: 523.7488 (2+)
Er.: 1.7 ppm
523.5
525.1
m/z
500
520
540
560
580
600
620
640
660
680
700
m/z

**3-III**

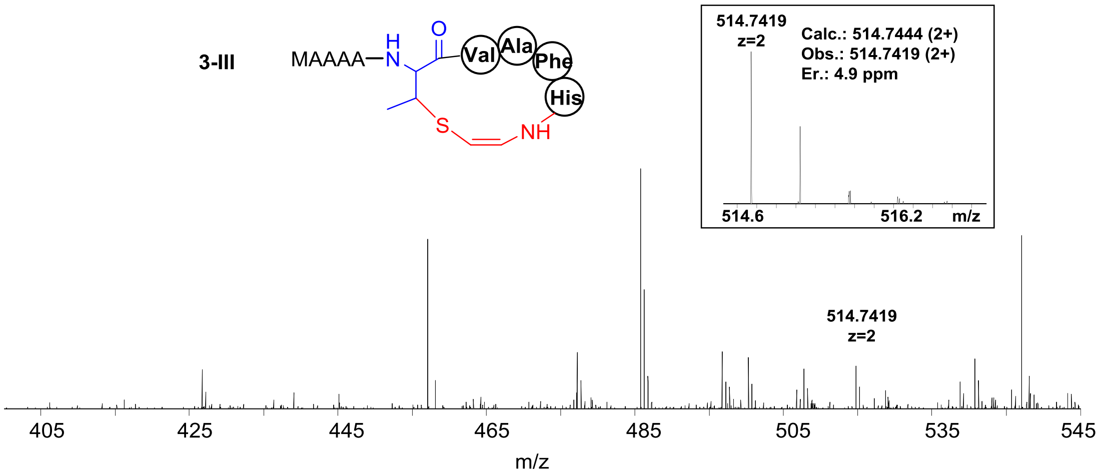
MAAAA
H
N
O
Val
Ala
Phe
His
S
NH
514.7419
z=2
Calc.: 514.7444 (2+)
Obs.: 514.7419 (2+)
Er.: 4.9 ppm
514.6
516.2
m/z
514.7419
z=2
405
425
445
465
485
505
535
545
m/z

**3-II**

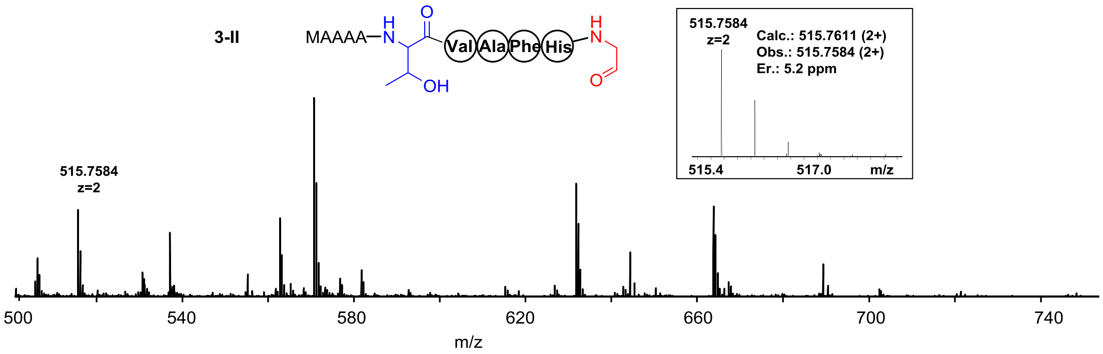
MAAAA
H
N
O
OH
Val
Ala
Phe
His
H
N
O
515.7584
z=2
Calc.: 515.7611 (2+)
Obs.: 515.7584 (2+)
Er.: 5.2 ppm
515.4
517.0
m/z
515.7584
z=2
500
540
580
620
660
700
740
m/z

**4**

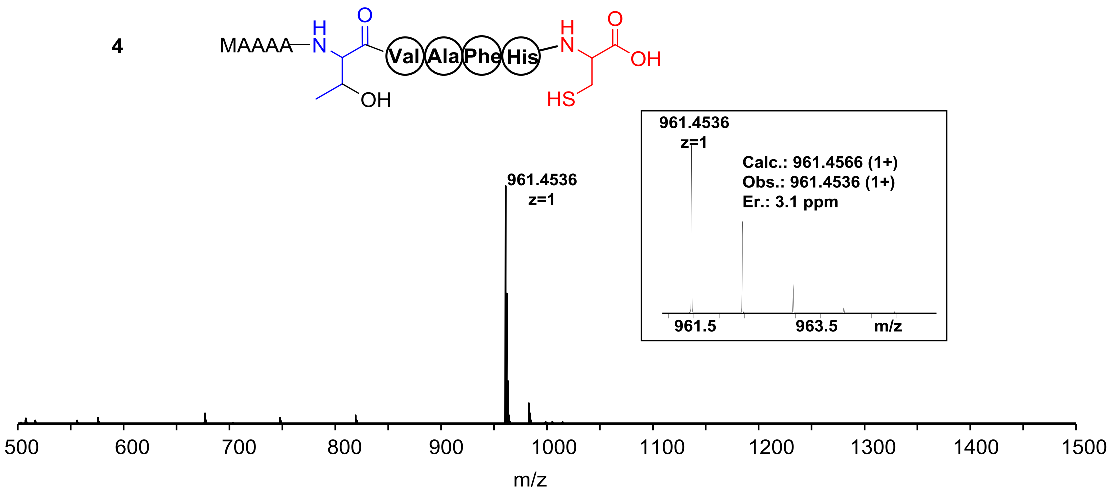
MAAAA
H
N
O
Val
Ala
Phe
His
H
N
O
OH
OH
HS
961.4536
z=1
Calc.: 961.4566 (1+)
Obs.: 961.4536 (1+)
Er.: 3.1 ppm
961.5
963.5
m/z
500
600
700
800
900
1000
1100
1200
1300
1400
1500
m/z

**4-I**

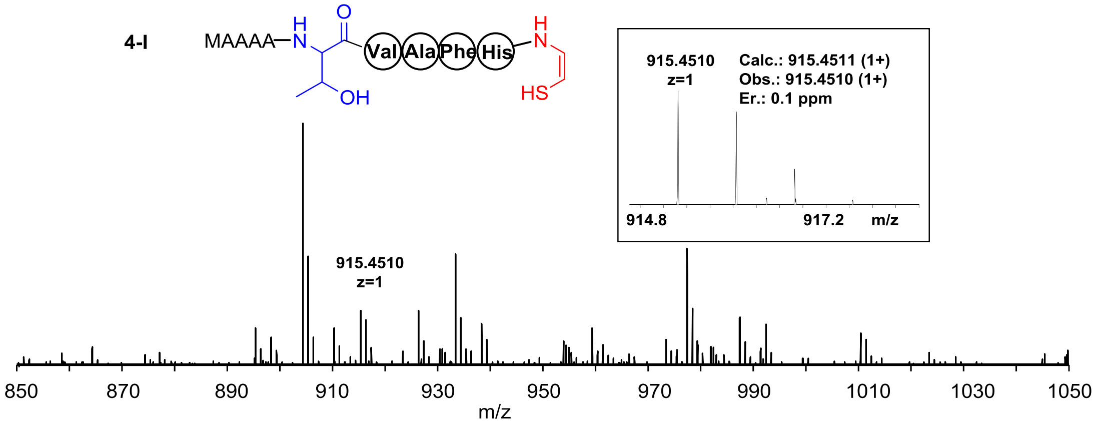
MAAAA
H
N
O
Val
Ala
Phe
His
H
N
OH
HS
915.4510
z=1
Calc.: 915.4511 (1+)
Obs.: 915.4510 (1+)
Er.: 0.1 ppm
914.8
917.2
m/z
850
870
890
910
930
950
970
990
1010
1030
1050
m/z

**5**

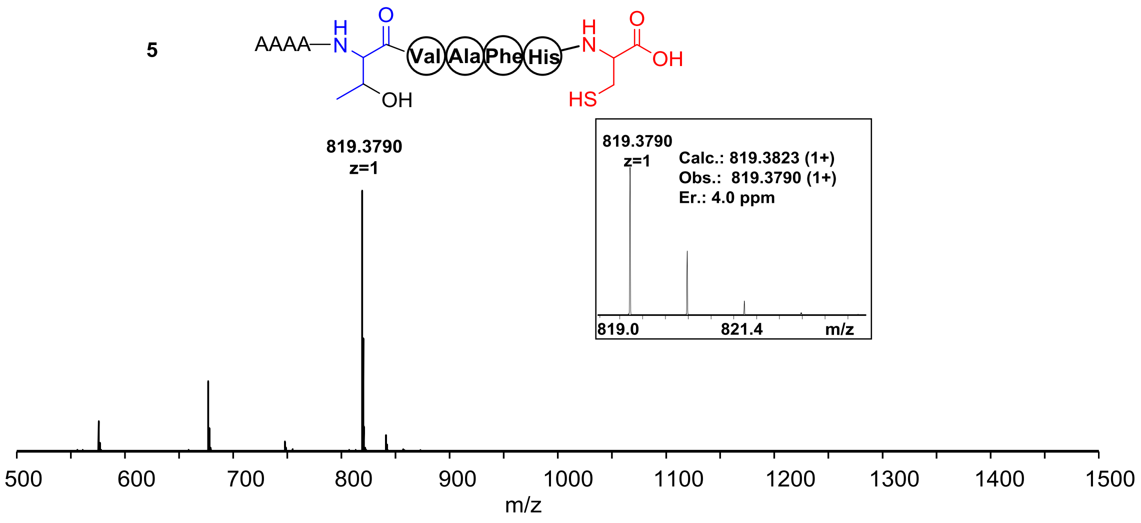
AAAA
Val
Ala
Phe
His
OH
HS
819.3790
z=1
Calc.: 819.3823 (1+)
Obs.: 819.3790 (1+)
Er.: 4.0 ppm
819.0
821.4
m/z

**5-I**

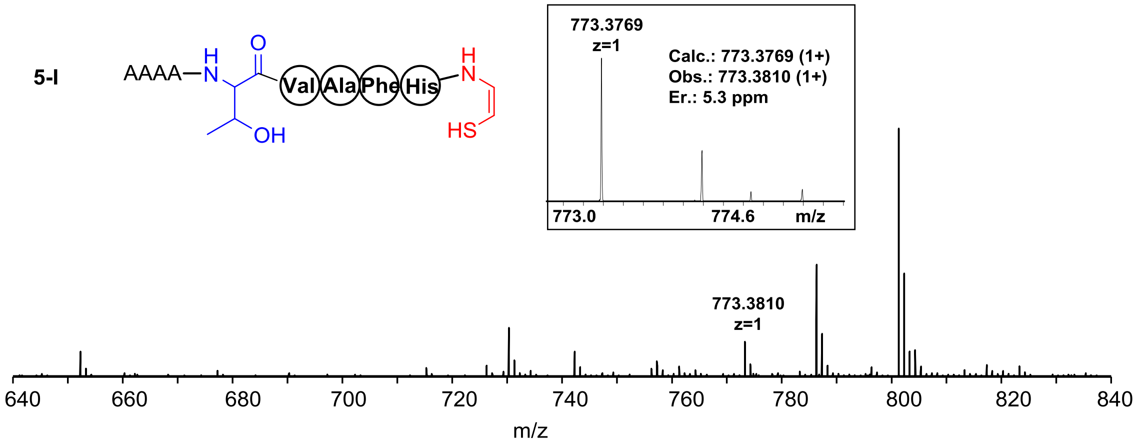
AAAA
Val
Ala
Phe
His
OH
HS
773.3769
z=1
Calc.: 773.3769 (1+)
Obs.: 773.3810 (1+)
Er.: 5.3 ppm
773.0
774.6
m/z
773.3810
z=1

6

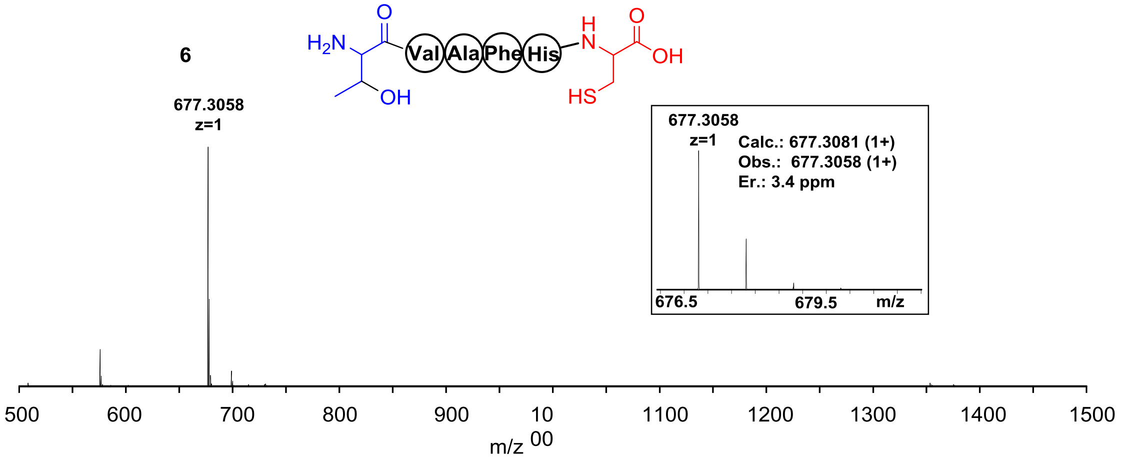
H2N
O
Val
Ala
Phe
His
H
N
O
OH
OH
HS
677.3058
z=1
677.3058
z=1
Calc.: 677.3081 (1+)
Obs.: 677.3058 (1+)
Er.: 3.4 ppm
676.5
679.5
m/z
500
600
700
800
900
10
00
1100
1200
1300
1400
1500
m/z

6-I

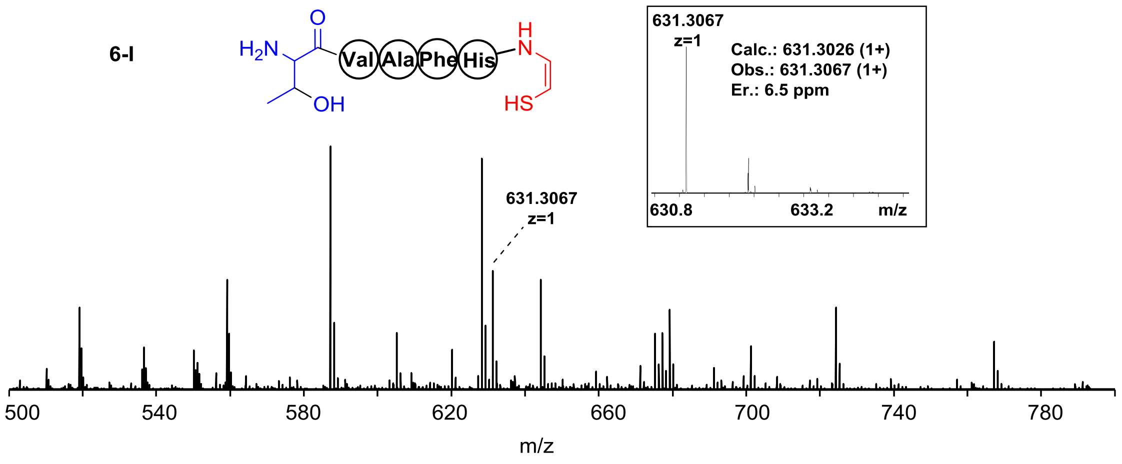
H2N
O
Val
Ala
Phe
His
H
N
OH
HS
631.3067
z=1
Calc.: 631.3026 (1+)
Obs.: 631.3067 (1+)
Er.: 6.5 ppm
630.8
633.2
m/z
631.3067
z=1
500
540
580
620
660
700
740
780
m/z

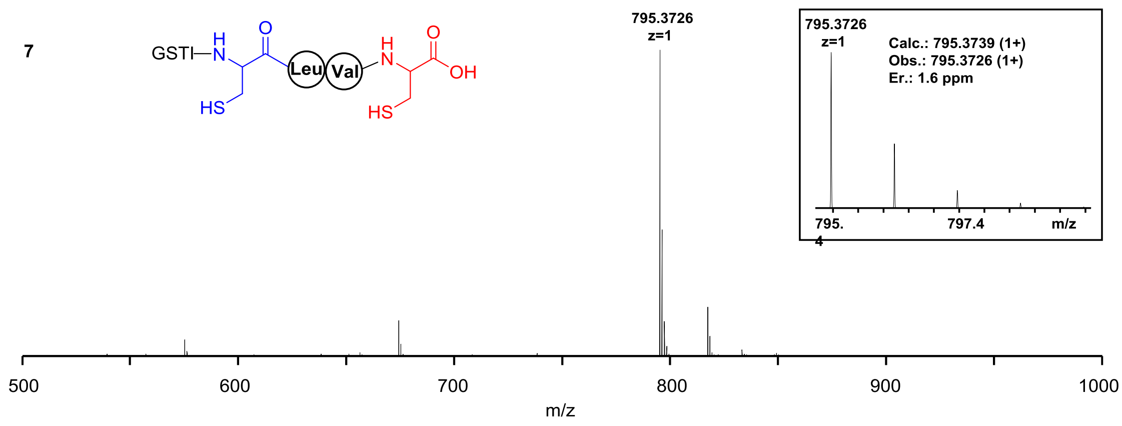


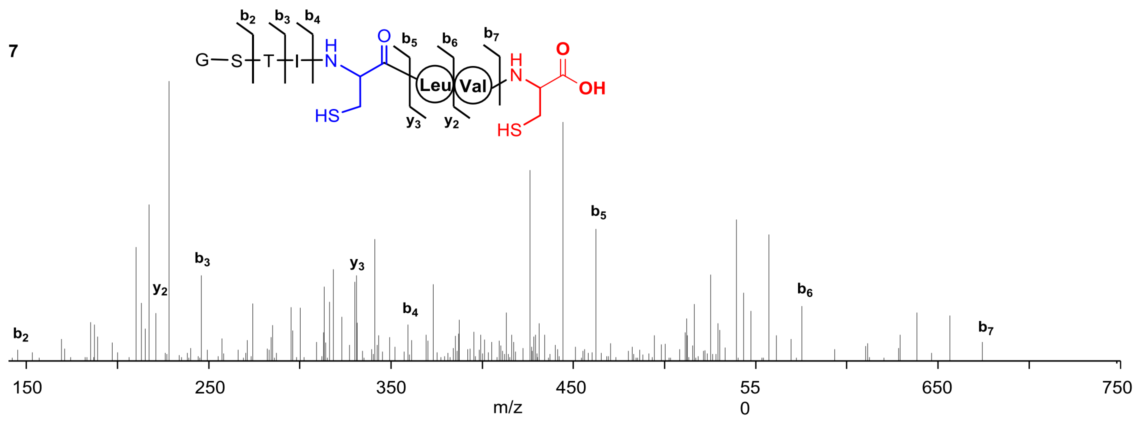


| Ions | Calcd. | Obs. | Er. (ppm) |
|---|---|---|---|
| $b_2$ | 145.0613 | 145.0606 | 5.5 |
| $b_3$ | 246.1090 | 246.1080 | 4.5 |
| $b_4$ | 359.1931 | 359.1917 | 3.6 |
| $b_5$ | 462.2022 | 462.2008 | 3.0 |
| $b_6$ | 575.2863 | 575.2847 | 3.1 |
| $b_7$ | 674.3547 | 674.3527 | 4.2 |
| $y_2$ | 221.0959 | 221.0951 | 4.1 |
| $y_3$ | 334.1800 | 334.1786 | 5.1 |

7-l

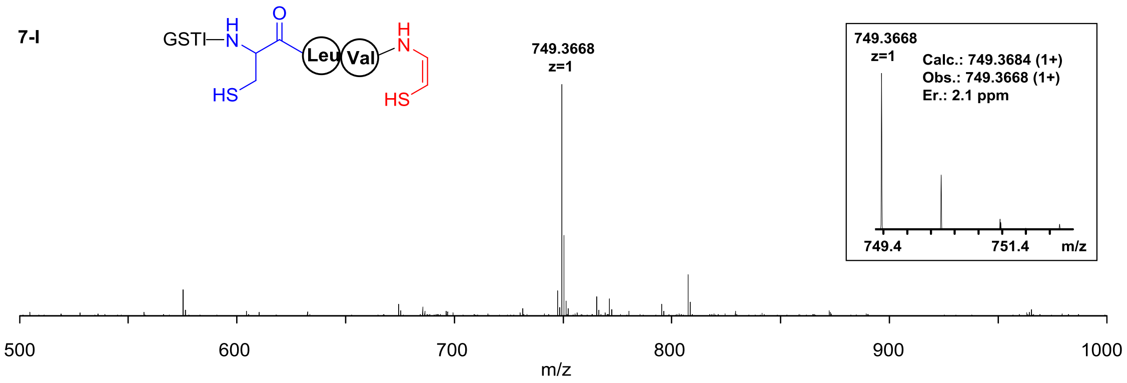


7-l

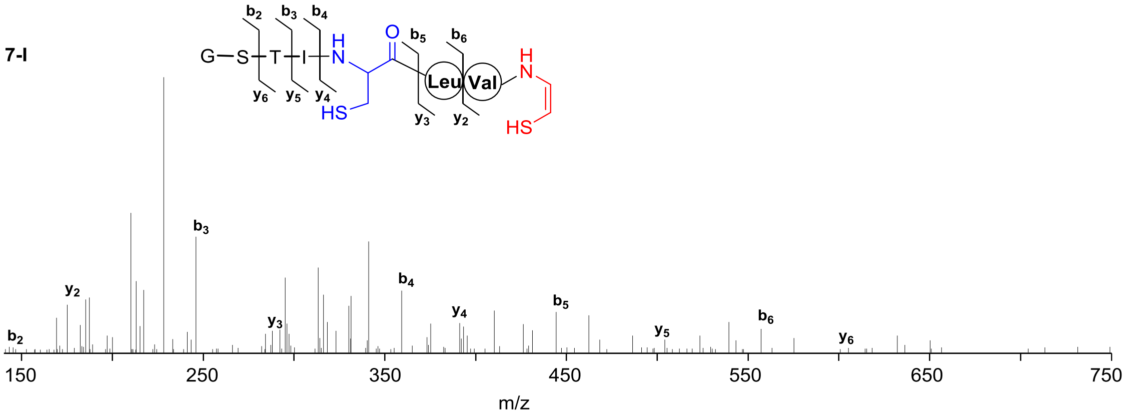


| Ions | Calcd. | Obs. | Er. (ppm) |
|---|---|---|---|
| $b_2$ | 145.0613 | 145.0606 | 4.8 |
| $b_3$ | 246.1090 | 246.1079 | 4.5 |
| $b_4$ | 359.1931 | 359.1917 | 3.9 |
| $b_5$ | 462.2022 | 462.2006 | 3.5 |
| $b_6$ | 575.2863 | 575.2836 | 4.7 |
| $y_2$ | 175.0905 | 175.0896 | 5.1 |
| $y_3$ | 288.1745 | 288.1731 | 4.9 |
| $y_4$ | 391.1837 | 391.1823 | 3.6 |
| $y_5$ | 504.2678 | 504.2656 | 4.4 |
| $y_6$ | 605.3154 | 605.3120 | 5.6 |

**7-II**

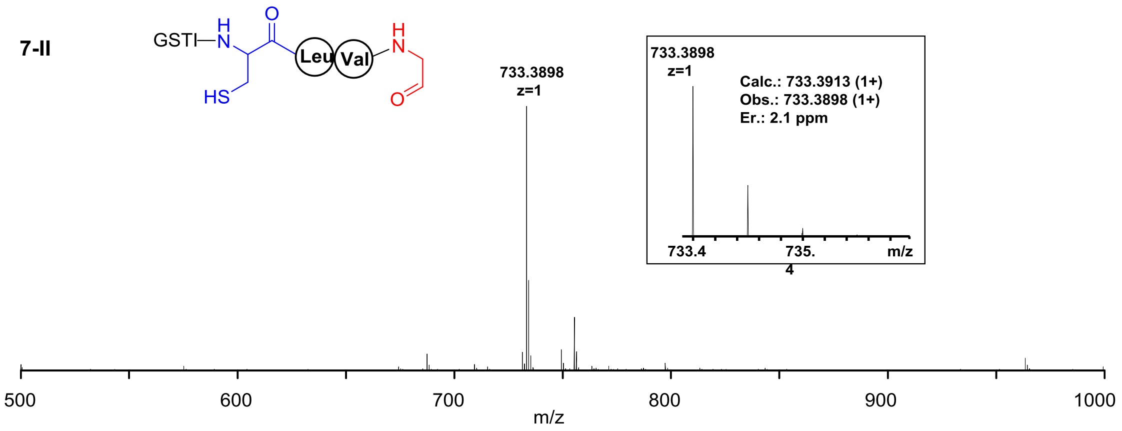


**7-II**

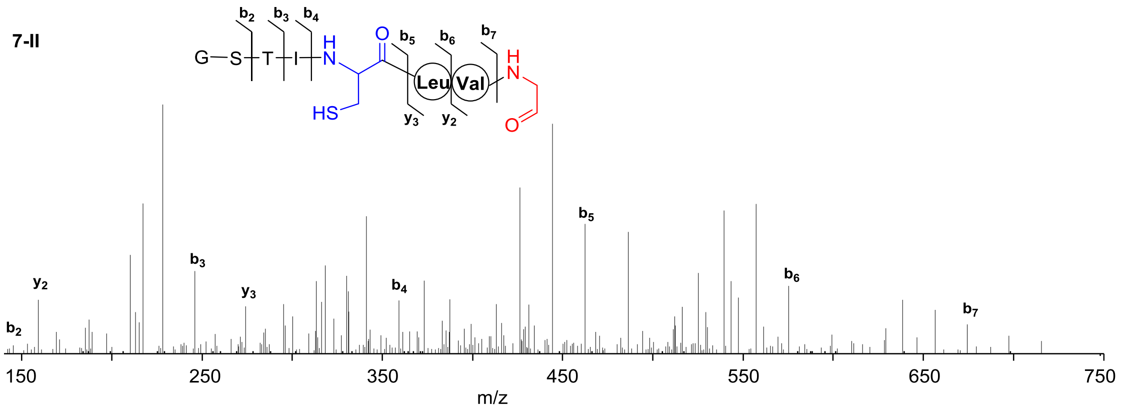


| Ions | Calcd. | Obs. | Er. (ppm) |
|---|---|---|---|
| $b_2$ | 145.0613 | 145.0605 | 5.5 |
| $b_3$ | 246.1090 | 246.1079 | 4.5 |
| $b_4$ | 359.1931 | 359.1917 | 3.9 |
| $b_5$ | 462.2022 | 462.2006 | 3.5 |
| $b_6$ | 575.2863 | 575.2844 | 3.3 |
| $b_7$ | 674.3547 | 674.3523 | 3.6 |
| $y_2$ | 159.1133 | 159.1125 | 5.0 |
| $y_3$ | 272.1974 | 272.1961 | 4.8 |

**7-III**

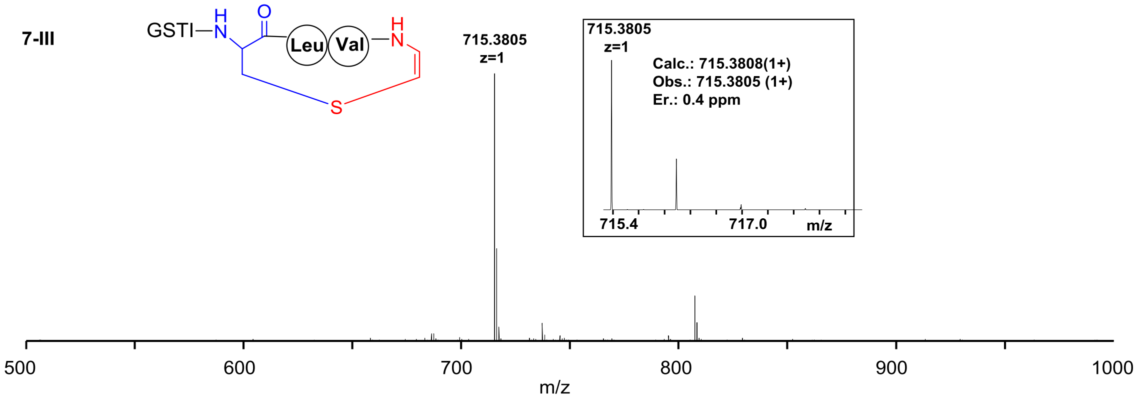


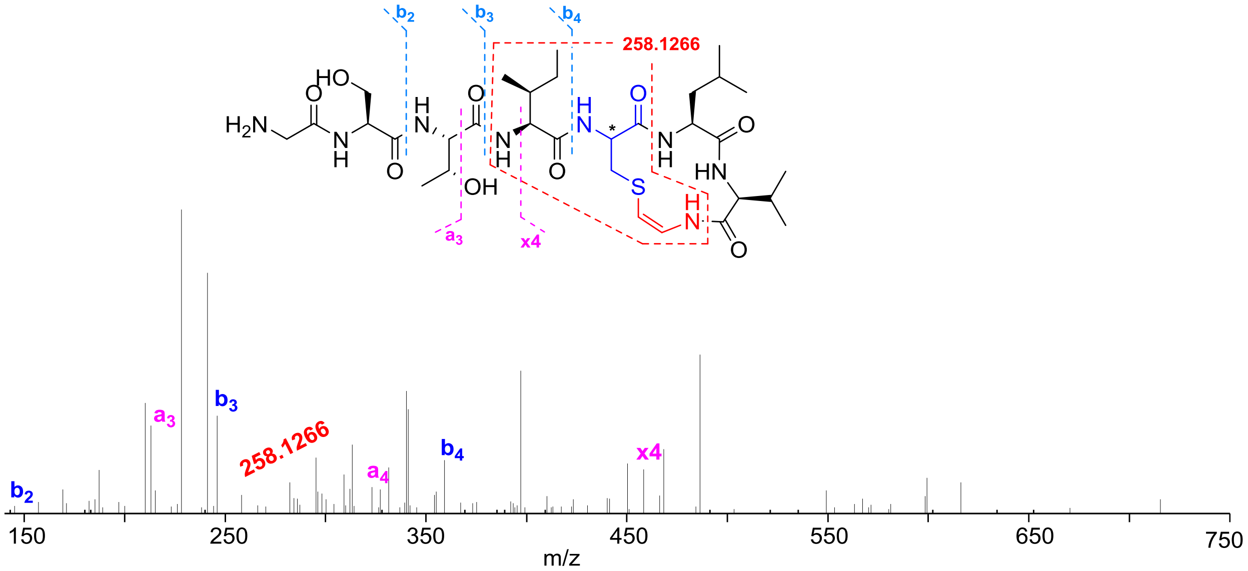


| Ions | Calcd. | Obs. | Er. (ppm) |
|---|---|---|---|
| **b₂** | 145.0613 | 145.0607 | 4.1 |
| **b₃** | 246.1090 | 246.1080 | 4.1 |
| **b₄** | 359.1931 | 359.1918 | 3.6 |
| **a₃** | 218.1135 | 218.1134 | 0.5 |
| **a₄** | 331.1976 | 331.1969 | 2.1 |
| **x₄** | 453.2530 | 453.2513 | 3.8 |
| | 258.1271 | 258.1266 | 1.9 |

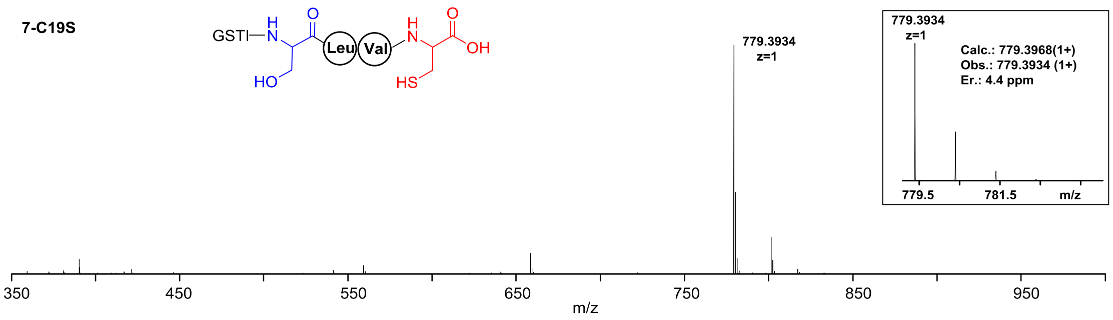


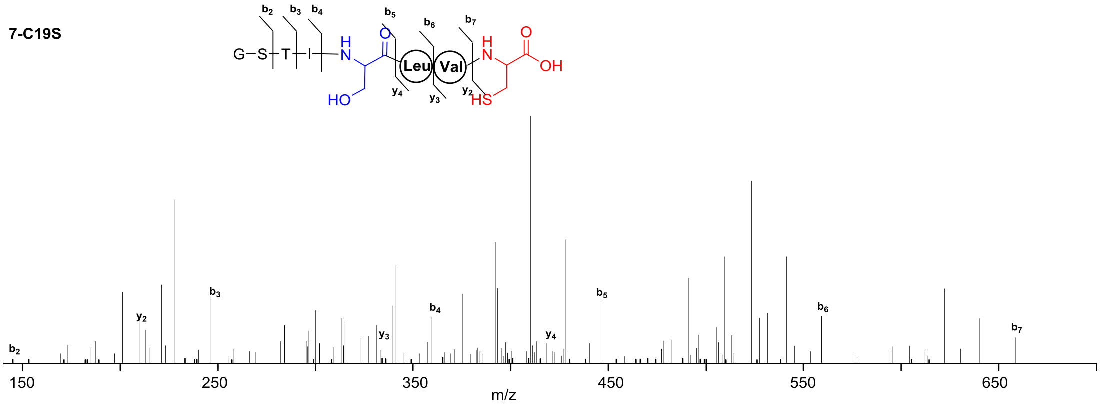


| Ions | Calcd. | Obs. | Er. (ppm) |
|---|---|---|---|
| **$b_2$** | 145.0613 | 145.0609 | 2.8 |
| **$b_3$** | 246.1090 | 246.1084 | 2.4 |
| **$b_4$** | 359.1931 | 359.1923 | 2.2 |
| **$b_5$** | 446.2251 | 446.2247 | 0.9 |
| **$b_6$** | 559.3092 | 559.3085 | 1.3 |
| **$b_7$** | 658.3776 | 658.3774 | 0.3 |
| **$y_2$** | 221.0959 | 221.0955 | 1.8 |
| **$y_3$** | 334.1800 | 334.1795 | 1.5 |
| **$y_4$** | 421.2120 | 421.2110 | 2.4 |

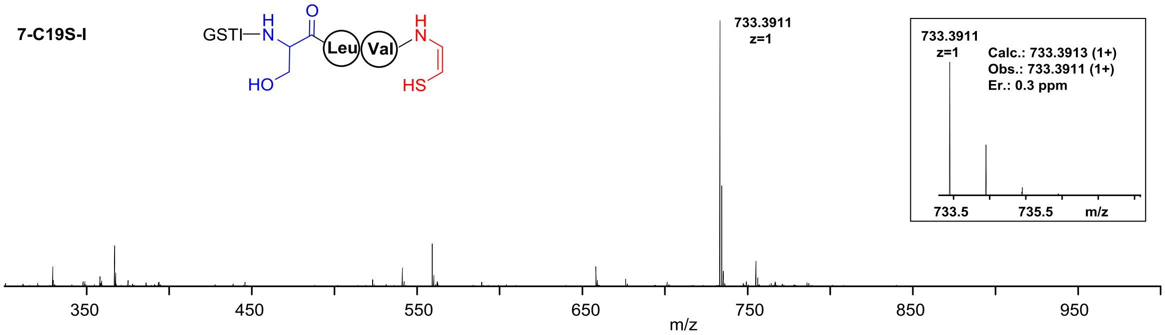


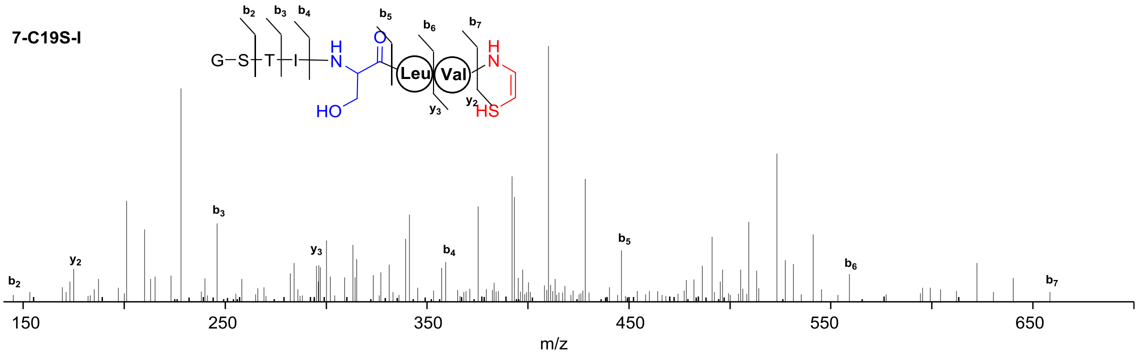


| Ions | Calcd. | Obs. | Er. (ppm) |
|---|---|---|---|
| $b_2$ | 145.0613 | 145.0608 | 3.4 |
| $b_3$ | 246.1090 | 246.1083 | 2.8 |
| $b_4$ | 359.1931 | 359.1922 | 2.5 |
| $b_5$ | 446.2251 | 446.2244 | 1.6 |
| $b_6$ | 559.3092 | 559.3081 | 2.0 |
| $b_7$ | 658.3776 | 658.3769 | 1.1 |
| $y_2$ | 175.0905 | 175.0900 | 2.9 |
| $y_3$ | 288.1745 | 288.1739 | 2.1 |

**7-C19S-III**

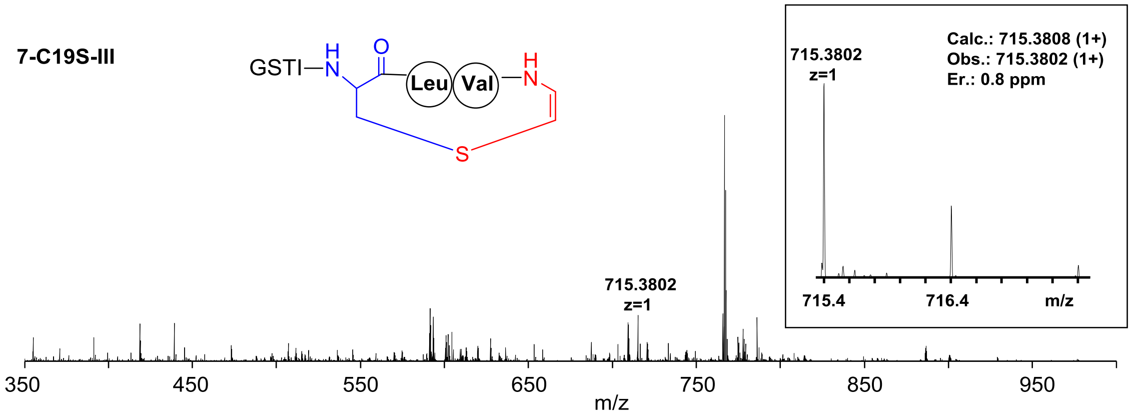


**7-C19S-II**

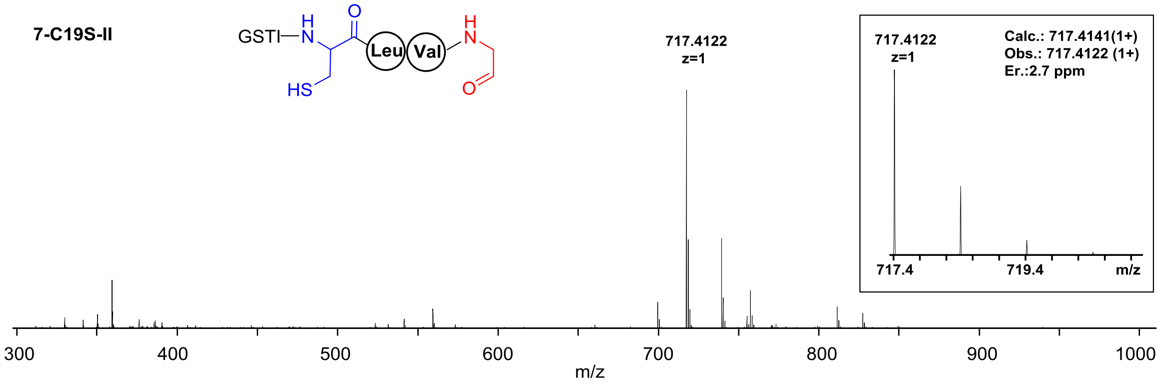

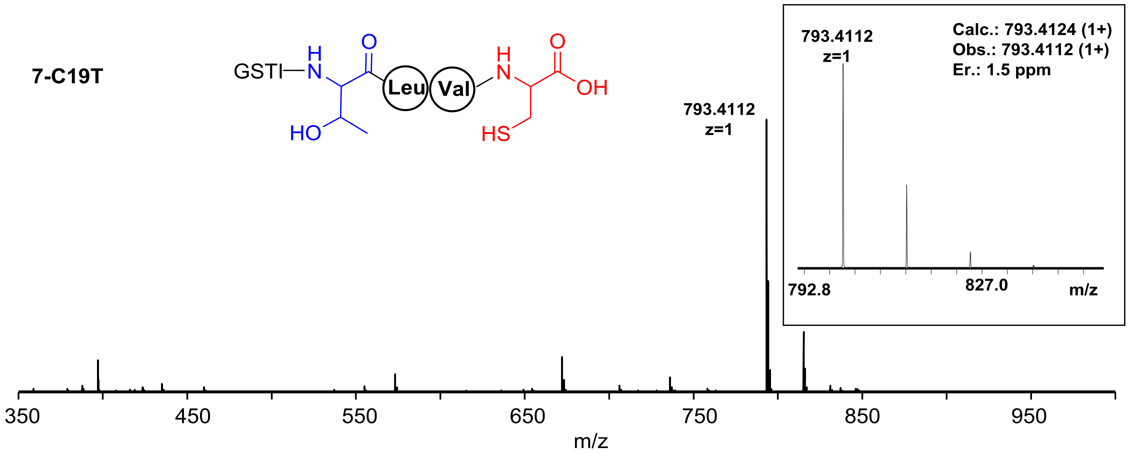


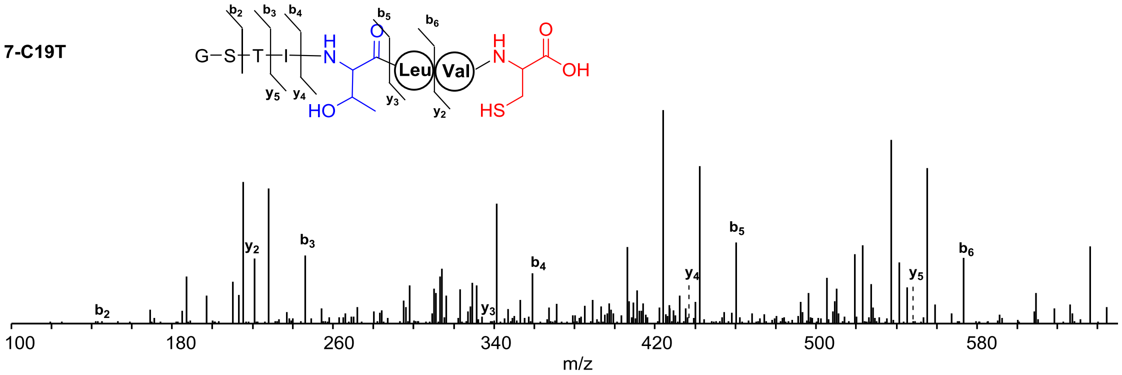


| Ions | Calcd. | Obs. | Er. (ppm) |
|---|---|---|---|
| $b_2$ | 145.0613 | 145.0607 | 4.1 |
| $b_3$ | 246.1090 | 246.1085 | 2.0 |
| $b_4$ | 359.1931 | 359.1925 | 1.7 |
| $b_5$ | 460.2407 | 460.2403 | 0.9 |
| $b_6$ | 573.3248 | 573.3245 | 0.5 |
| $y_2$ | 221.0959 | 221.0955 | 1.8 |
| $y_3$ | 334.1800 | 334.1795 | 1.5 |
| $y_4$ | 435.2277 | 435.2274 | 0.7 |
| $y_5$ | 548.3119 | 548.3126 | 1.3 |

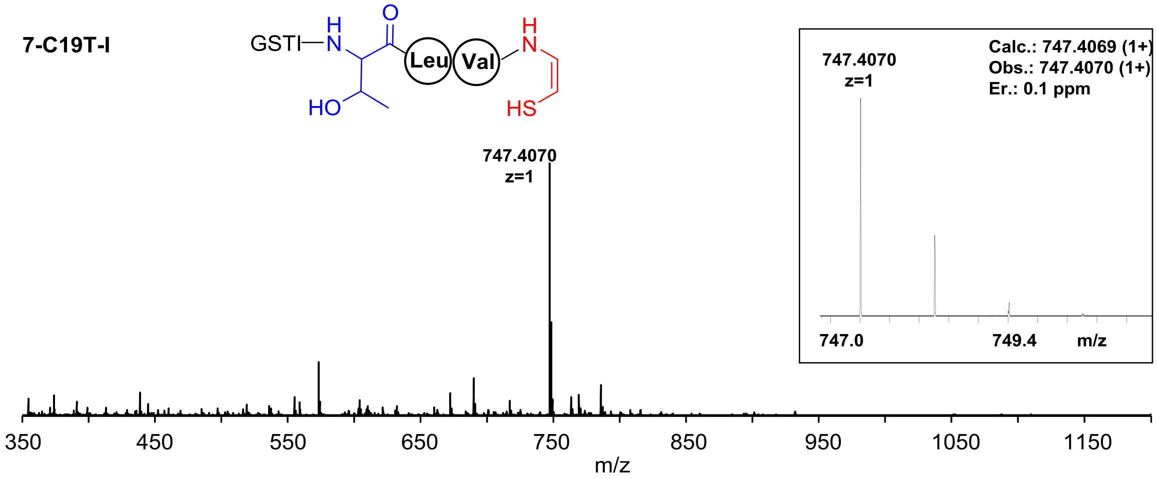


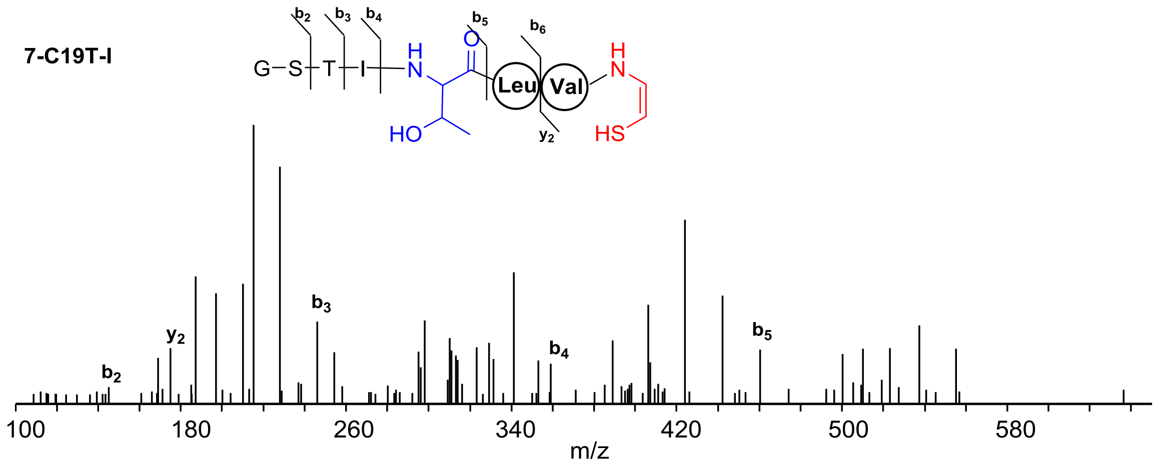


| Ions | Calcd. | Obs. | Er. (ppm) |
|---|---|---|---|
| $b_2$ | 145.0613 | 145.0607 | 4.1 |
| $b_3$ | 246.1090 | 246.1082 | 3.2 |
| $b_4$ | 359.1931 | 359.1920 | 3.1 |
| $b_5$ | 460.2407 | 460.2396 | 2.4 |
| $y_2$ | 175.0905 | 175.0900 | 2.9 |

**7-C19T-II**

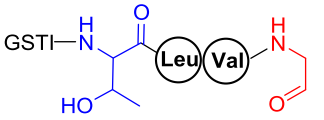
GSTI
H
N
O
HO
Leu
Val
H
N
O

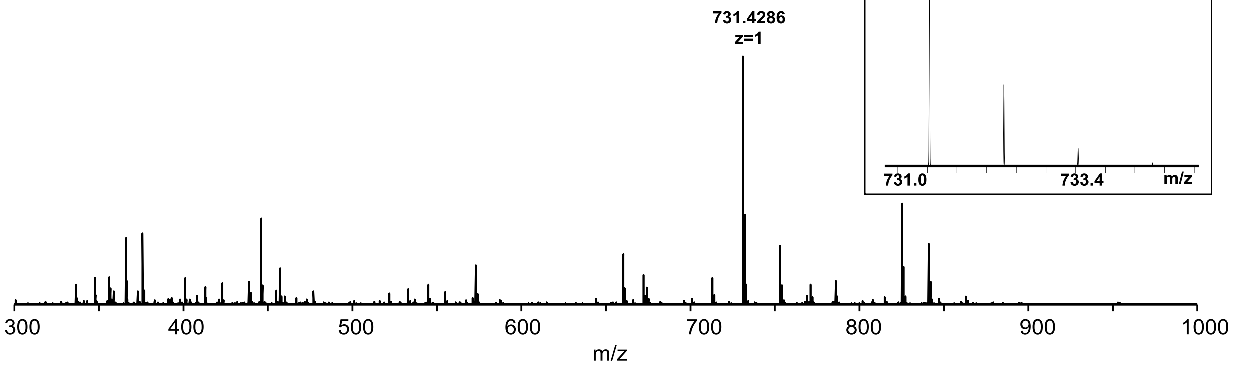
731.4286
z=1
Calc.: 731.4398 (1+)
Obs.: 731.4286 (1+)
Er.: 1.6 ppm
731.4286
z=1
731.0
733.4
m/z
300
400
500
600
700
800
900
1000
m/z

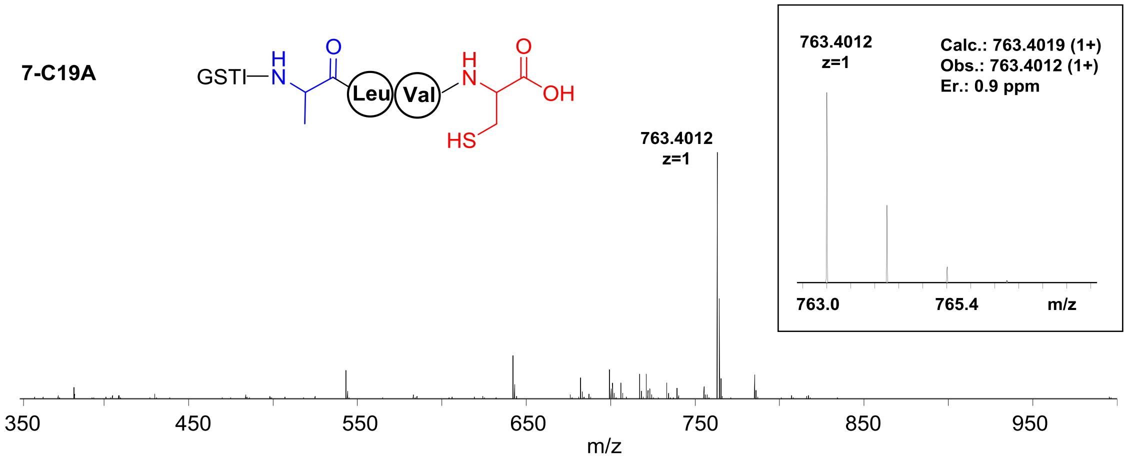


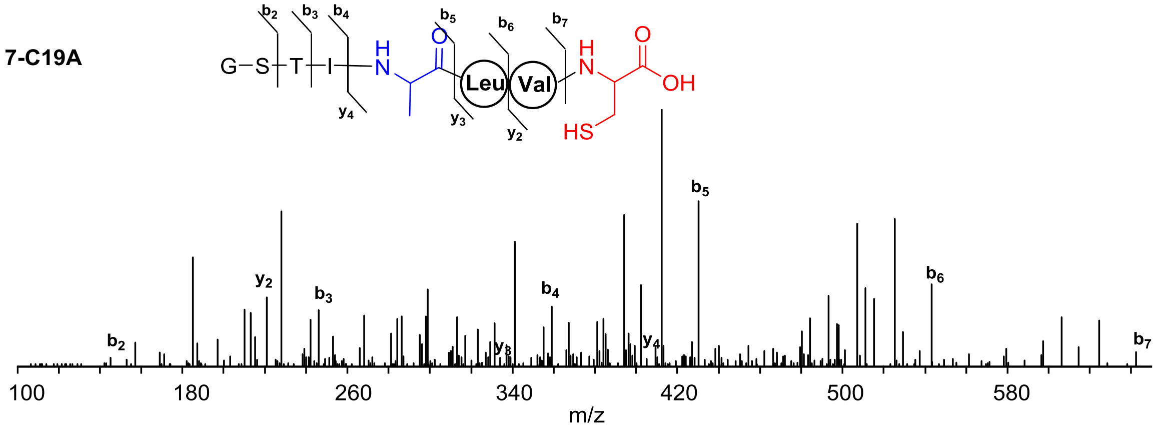


| Ions | Calcd. | Obs. | Er. (ppm) |
|---|---|---|---|
| $b_2$ | 145.0613 | 145.0603 | 6.9 |
| $b_3$ | 246.1090 | 246.1078 | 4.9 |
| $b_4$ | 359.1931 | 359.1915 | 4.5 |
| $b_5$ | 430.2302 | 430.2286 | 3.7 |
| $b_6$ | 543.3142 | 543.3124 | 3.3 |
| $b_7$ | 642.3827 | 642.3804 | 3.6 |
| $y_2$ | 221.0959 | 221.0949 | 4.5 |
| $y_3$ | 334.1800 | 334.1784 | 4.8 |
| $y_4$ | 405.2171 | 405.2165 | 1.5 |

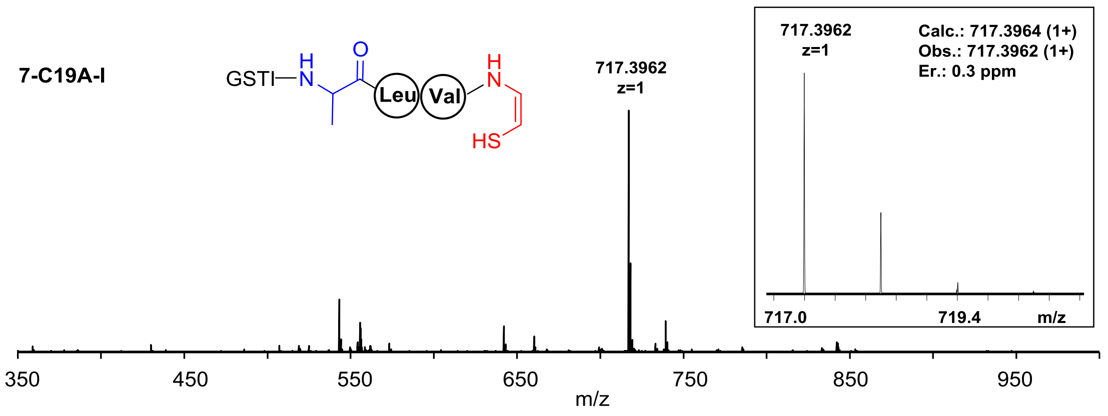


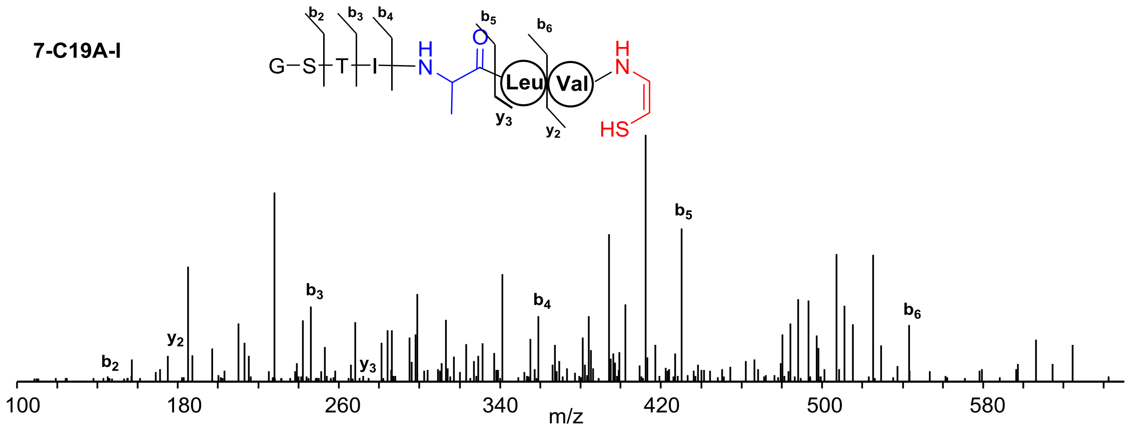


| Ions | Calcd. | Obs. | Er. (ppm) |
|---|---|---|---|
| $b_2$ | 145.0613 | 145.0606 | 4.8 |
| $b_3$ | 246.1090 | 246.1081 | 3.7 |
| $b_4$ | 359.1931 | 359.1918 | 3.6 |
| $b_5$ | 430.2302 | 430.2292 | 2.3 |
| $b_6$ | 543.3142 | 543.3132 | 1.8 |
| $y_2$ | 175.0905 | 175.0898 | 4.0 |
| $y_3$ | 288.1745 | 288.1736 | 3.1 |

**7-C19A-II**

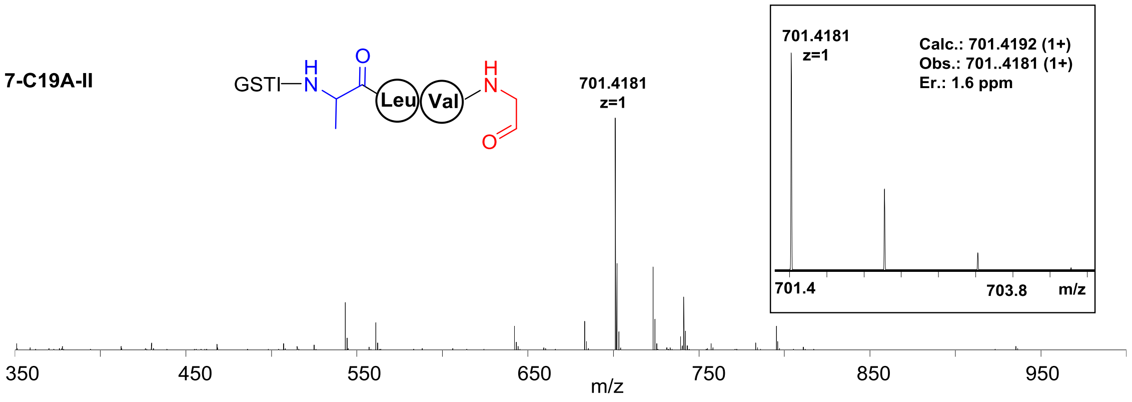

GSTI
H
N
O
Leu
Val
H
N
O
701.4181
z=1
Calc.: 701.4192 (1+)
Obs.: 701..4181 (1+)
Er.: 1.6 ppm
701.4
703.8
m/z
350
450
550
650
750
850
950
m/z

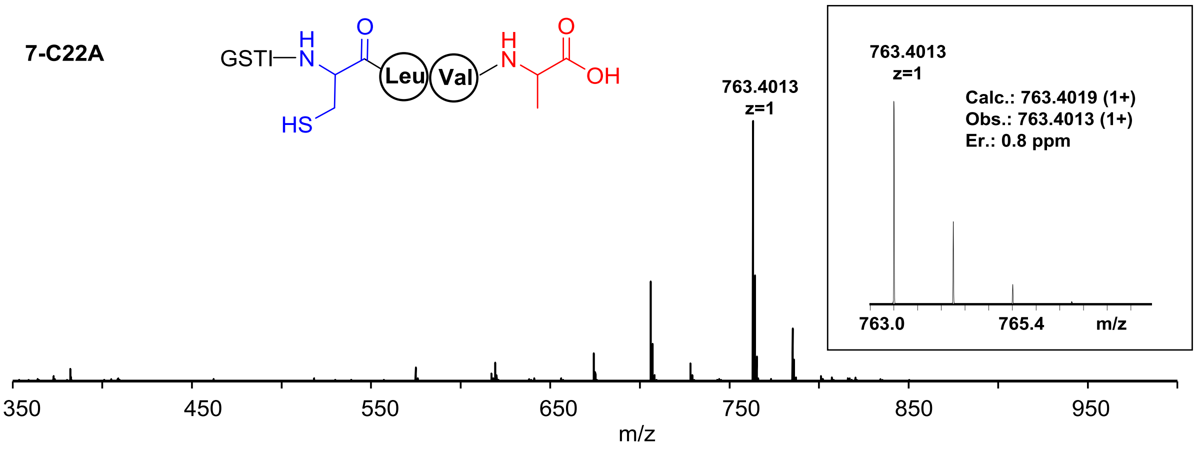
7-C22A
GSTI
H
N
O
Leu
Val
H
N
O
OH
HS
763.4013
z=1
763.4013
z=1
Calc.: 763.4019 (1+)
Obs.: 763.4013 (1+)
Er.: 0.8 ppm
763.0
765.4
m/z
350
450
550
650
750
850
950
m/z

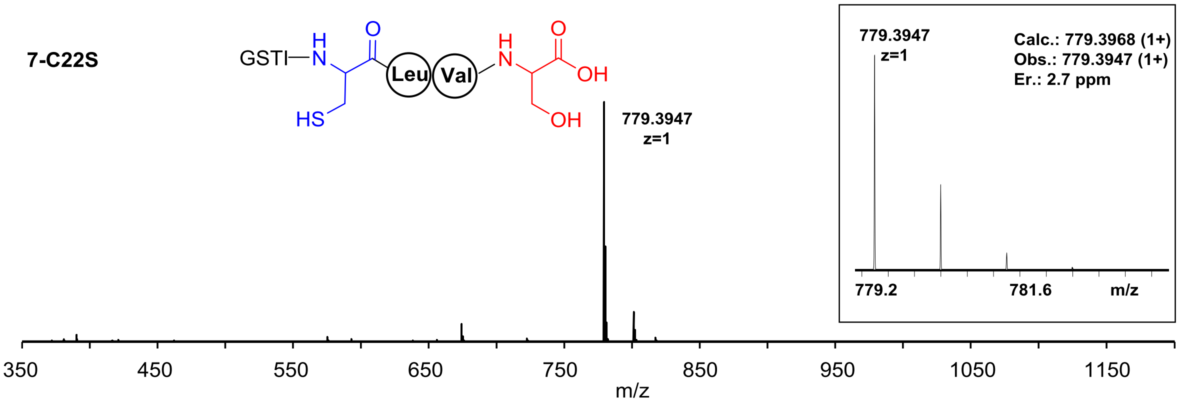
7-C22S
GSTI
H
N
O
Leu
Val
H
N
O
OH
HS
OH
779.3947
z=1
779.3947
z=1
Calc.: 779.3968 (1+)
Obs.: 779.3947 (1+)
Er.: 2.7 ppm
779.2
781.6
m/z
350
450
550
650
750
850
950
1050
1150
m/z

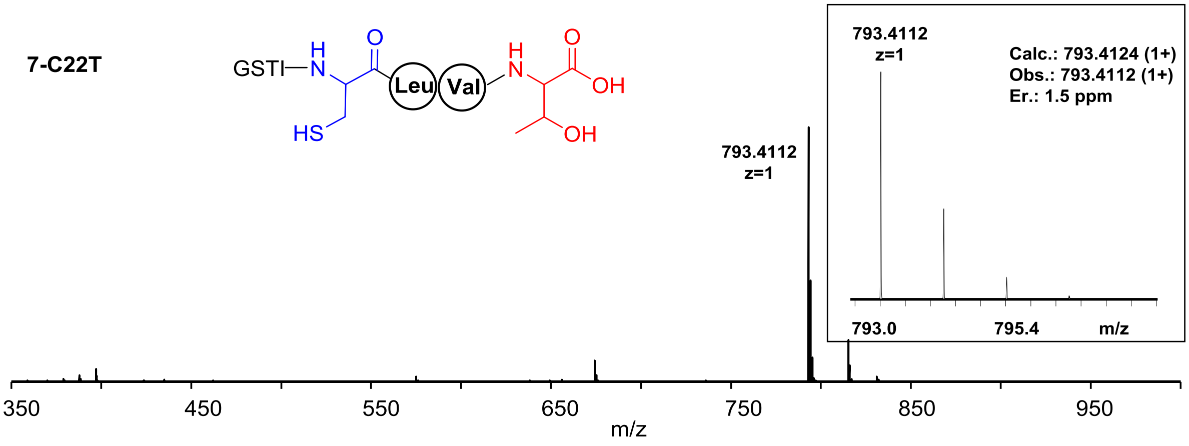
7-C22T
GSTI
H
N
O
Leu
Val
H
N
O
OH
HS
OH
793.4112
z=1
793.4112
z=1
Calc.: 793.4124 (1+)
Obs.: 793.4112 (1+)
Er.: 1.5 ppm
793.0
795.4
m/z
350
450
550
650
750
850
950
m/z

7-C19S-Ac

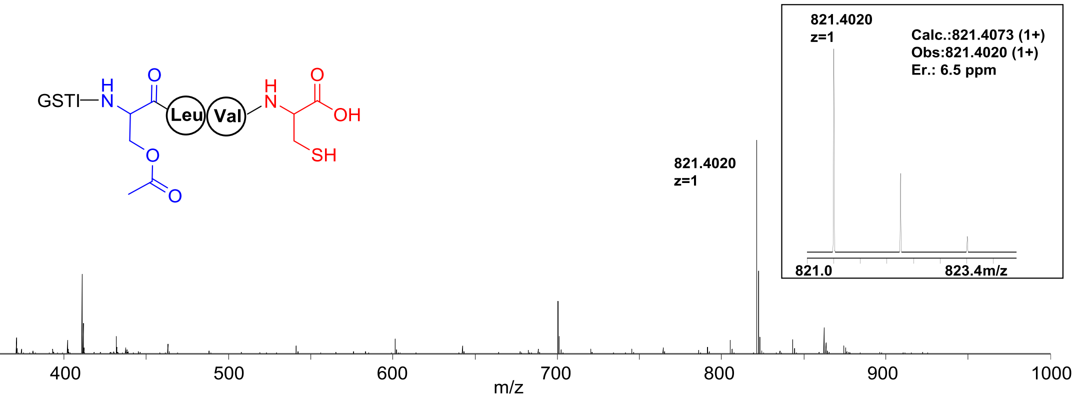


7-C19S-Ac-I

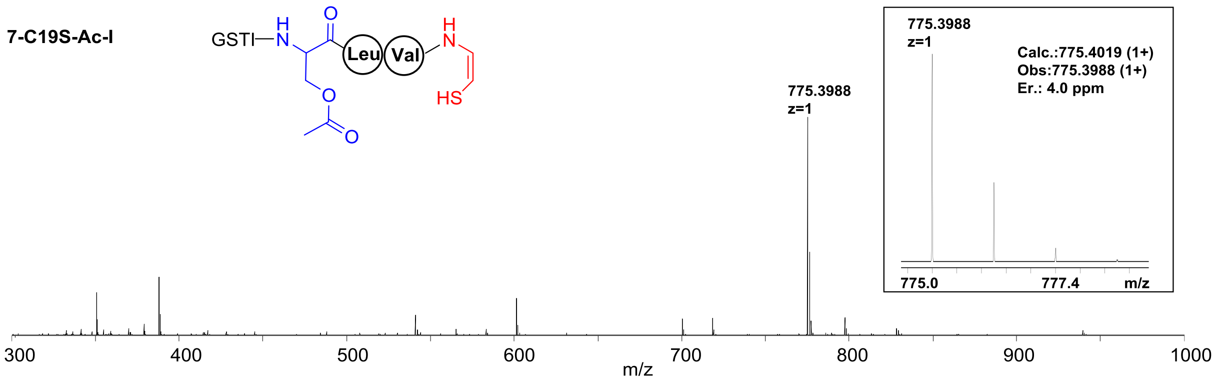


7-C19S-Ac-II

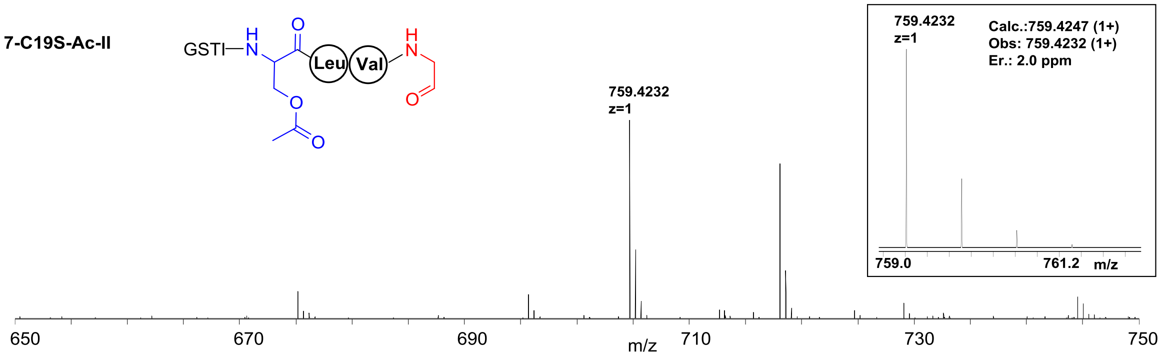


7-C19S-Ac-III

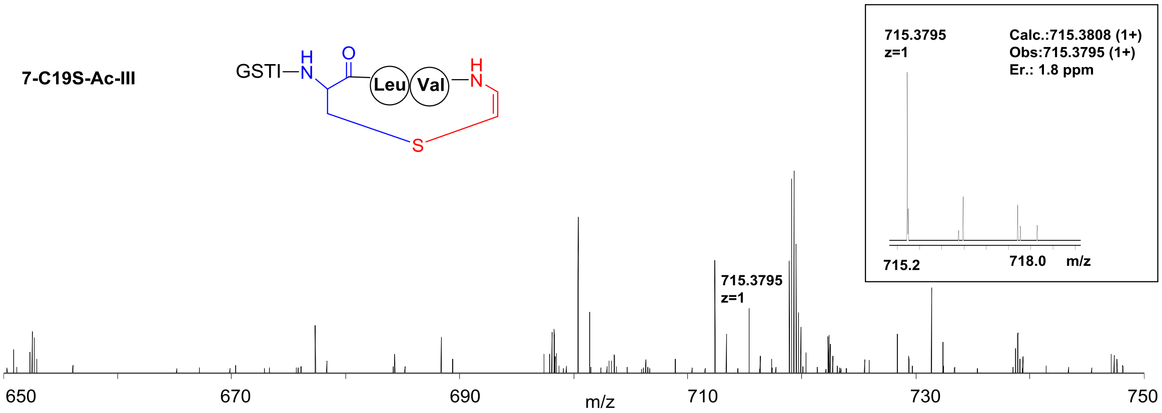

**7-C19S-P**

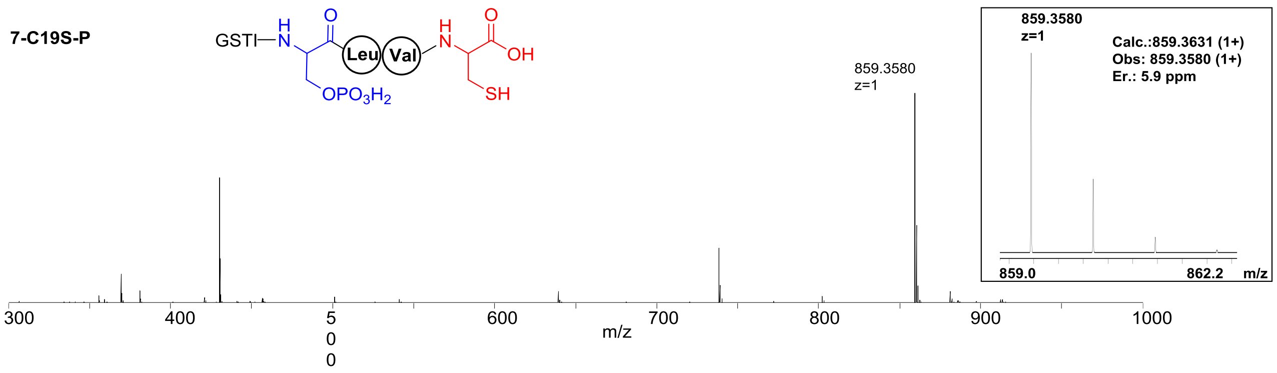


**7-C19S-Glu**

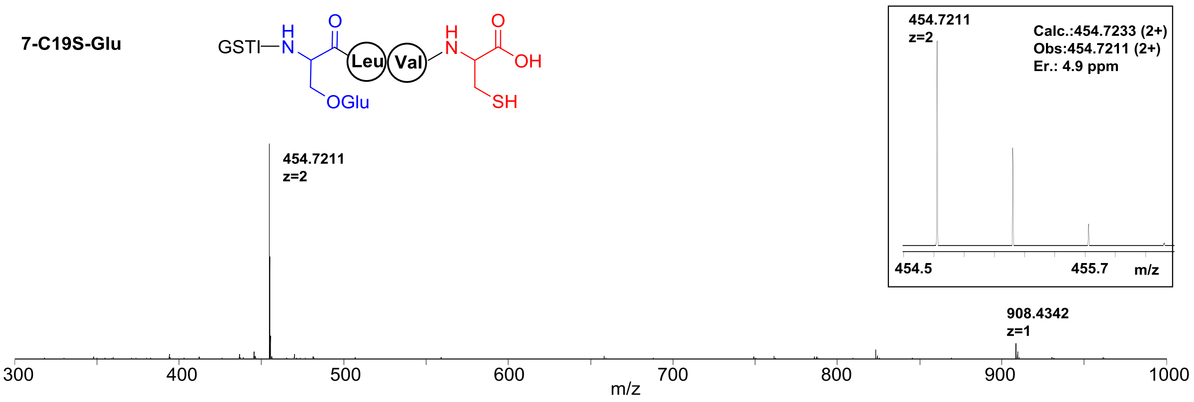

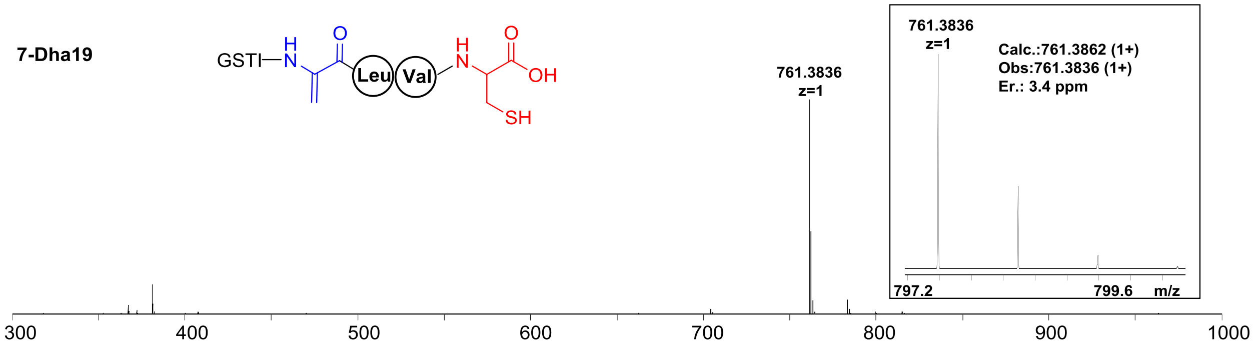


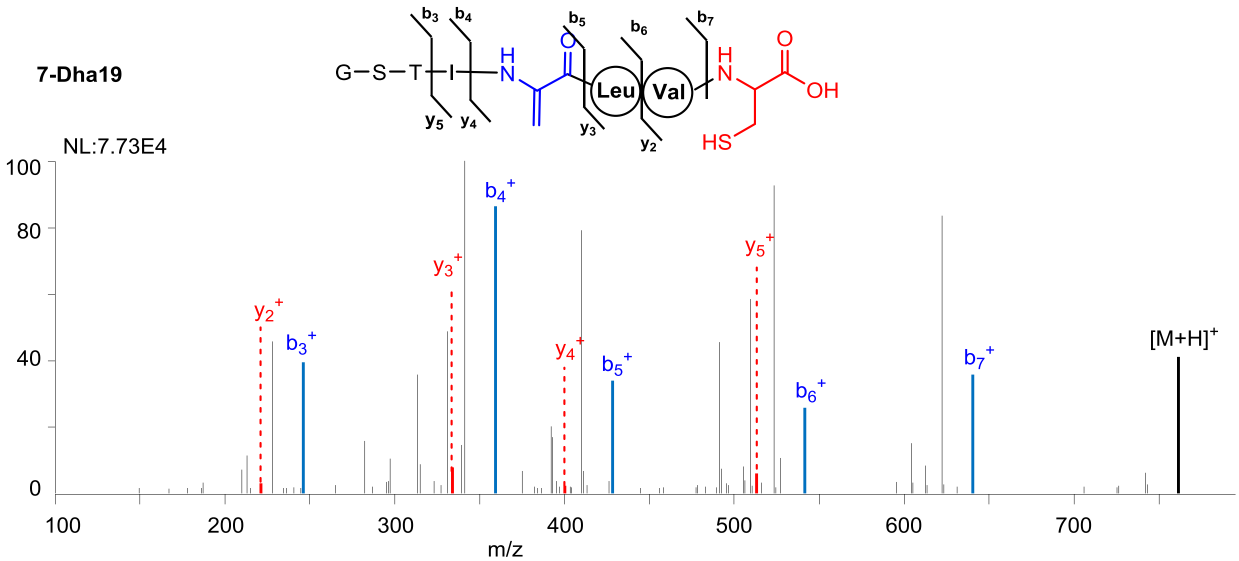


| Ions | Calcd. | Obs. | Er. (ppm) |
|---|---|---|---|
| $b_3$ | 246.1090 | 246.1082 | 3.3 |
| $b_4$ | 359.1931 | 359.1923 | 2.2 |
| $b_5$ | 428.2145 | 428.2140 | 1.2 |
| $b_6$ | 541.2986 | 541.2972 | 2.6 |
| $b_7$ | 640.3670 | 640.3655 | 2.3 |
| $y_2$ | 221.0960 | 221.0949 | 5.0 |
| $y_3$ | 334.1801 | 334.1785 | 4.8 |
| $y_4$ | 403.2015 | 403.2013 | 0.5 |
| $y_5$ | 516.2856 | 516.2870 | 2.7 |

**7-d-C19**

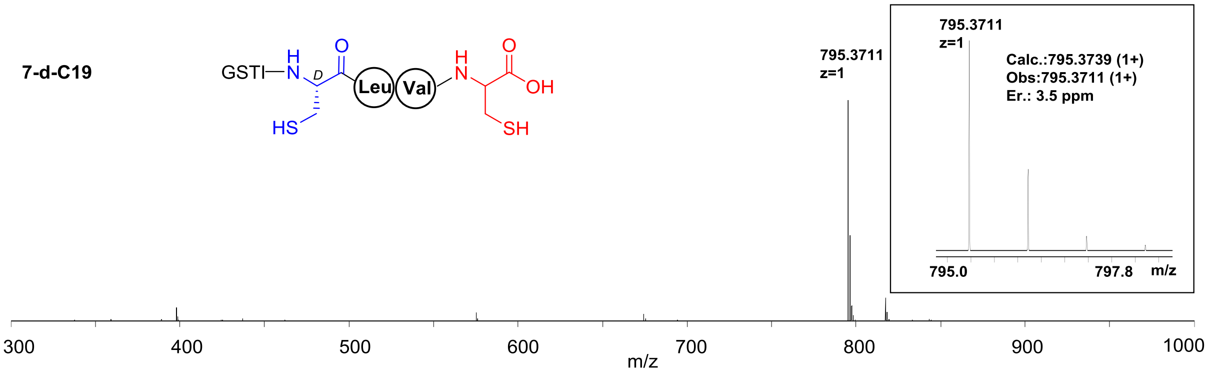
GSTI
H
N
D
O
Leu
Val
H
N
O
OH
HS
SH
795.3711
z=1
795.3711
z=1
Calc.:795.3739 (1+)
Obs:795.3711 (1+)
Er.: 3.5 ppm
795.0
797.8
m/z
300
400
500
600
700
800
900
1000
m/z

**7-IV**

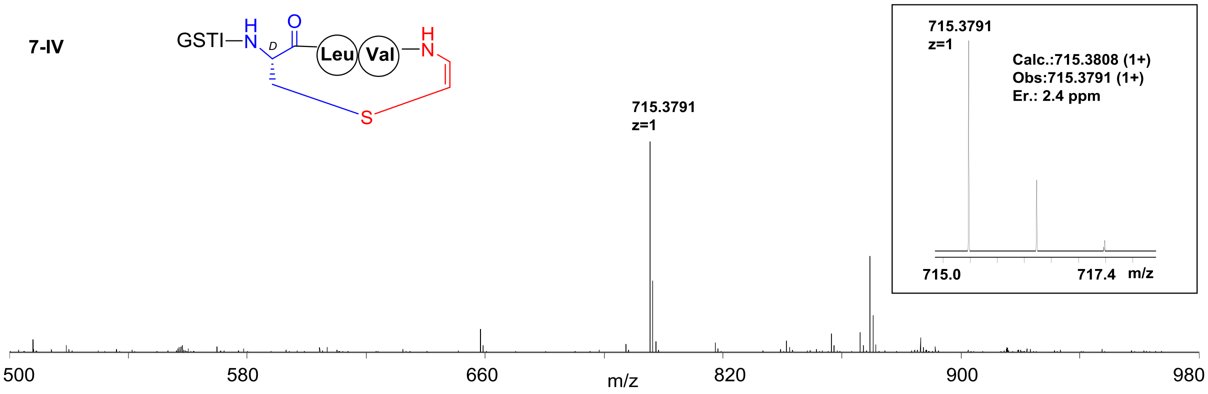
GSTI
H
N
D
O
Leu
Val
H
N
S
715.3791
z=1
715.3791
z=1
Calc.:715.3808 (1+)
Obs:715.3791 (1+)
Er.: 2.4 ppm
715.0
717.4
m/z
500
580
660
820
900
980
m/z

**7-C19S-D$_3$**

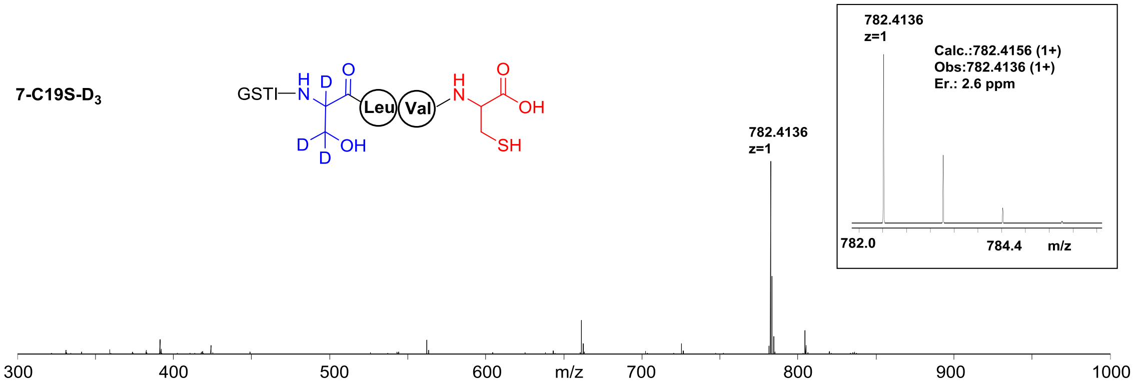


**7-C19S-D$_3$-I**

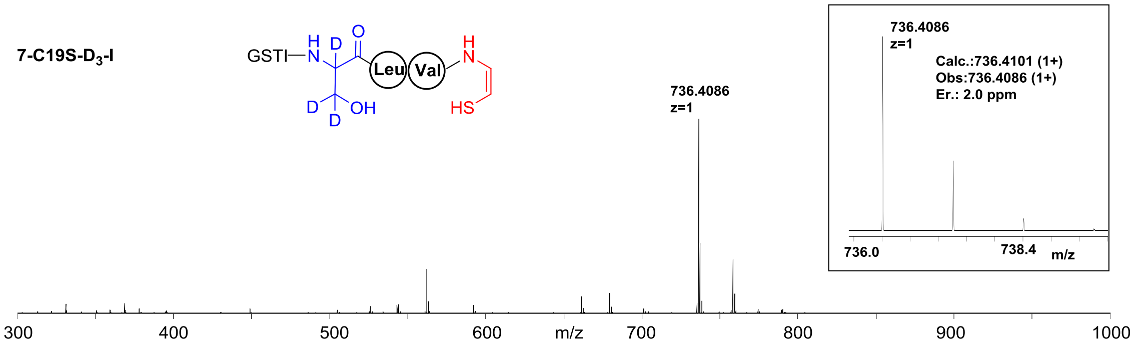


**7-C19S-D$_3$-II**

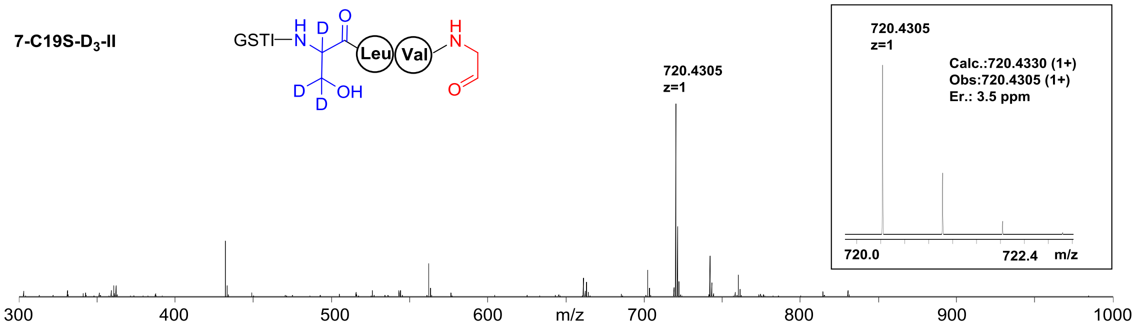

**Supplementary Table 5.** The $^{1}H$ and $^{13}C$ NMR data of **7** in DMSO-$d_6$.

| **Substructure** | $\delta_C$ | $\delta_H$, mult. ($J$, Hz) |
|---|---|---|
| **Gly** | 165.9 s | |
| | 40.2 t | 3.60 m (overlap)<br>3.60 m (overlap) |
| **Ser** | 169.9 s | |
| **Ser-NH** | | 8.55 d (7.8) |
| | 54.8 d | 4.56 q (6.7) |
| | 61.9 t | 3.59 m (overlap)<br>3.59 m (overlap) |
| **Thr** | 169.7 s | |
| **Thr-NH** | | 8.05 d (8.3) |
| | 58.3 d | 4.25 m (overlap) |
| | 66.3 d | 4.05 m (overlap) |
| | 19.8 q | 1.05 d (6.4) |
| **Ile** | 170.8 s | |
| **Ile-NH** | | 7.70 d (8.6) |
| | 56.8 d | 4.25 m (overlap) |
| | 36.9 d | 1.71 m (overlap) |
| | 24.1 t | 1.60 m (overlap)<br>1.43 m (overlap) |
| | 15.2 q | 0.82 m (overlap) |
| | 11.2 q | 0.79 m (overlap) |
| **Cys1** | 169.4 s | |
| **Cys1-NH** | | 8.20 m (overlap) |
| | 55.1 d | 4.40 m (overlap) |
| | 25.9 t | 2.75 m (overlap)<br>2.65 m (overlap) |
| **Leu** | 171.7 s | |
| **Leu-NH** | | 8.16 m (overlap) |
| | 51.3 d | 4.33 m (overlap) |
| | 40.6 t | 1.46 m (overlap)<br>1.46 m (overlap) |
| | 24.1 d | 1.06 m (overlap) |

| | | |
|---|---|---|
| | 21.5 q | 0.82 m (overlap) |
| | 23.1 q | 0.87 m (overlap) |
| **Val** | 170.9 s | |
| **Val-NH** | | 7.75 d (8.8) |
| | 57.4 d | 4.20 m (overlap) |
| | 30.7 d | 1.99 m (overlap) |
| | 19.1 q | 0.87 m (overlap) |
| | 18.0 q | 0.83 m (overlap) |
| **Cys2** | 171.4 s | |
| **Cys2-NH** | | 8.18 m (overlap) |
| | 54.4 d | 4.37 m (overlap) |
| | 25.4 t | 2.87 dd (13.7, 4.5)<br>2.75 m (overlap) |

**Supplementary Table 6.** Key data in the 2D-TOCSY f2-slice at f1 experiments in DMSO-$d_6$.

| | 2D-TOCSY f2-slice at f1 experiments (correlation signals) | |
|---|---|---|
| Residue (subunits) | Selective proton ($\delta_H$ in ppm, $J$ in Hz) | Target proton ($\delta_H$ in ppm, $J$ in Hz) |
| Ser | -NH, $\delta_H$ 8.50 | H2, $\delta_H$ 4.54 |
| Thr | -NH, $\delta_H$ 8.04 | H2, $\delta_H$ 4.25<br>H3, $\delta_H$ 4.22<br>H4, $\delta_H$ 1.21 |
| Ile | H2, $\delta_H$ 4.10 | H3, $\delta_H$ 1.62<br>H4a, $\delta_H$ 1.33, H4b, $\delta_H$ 1.32<br>H5, $\delta_H$ 0.84<br>H6, $\delta_H$ 0.81 |
| Leu | -NH, $\delta_H$ 8.21 | H2, $\delta_H$ 4.36<br>H3a, $\delta_H$ 1.48, H3b, $\delta_H$ 1.47<br>H4, $\delta_H$ 1.24<br>H5, $\delta_H$ 0.83<br>H6, $\delta_H$ 0.81 |
| Val | -NH, $\delta_H$ 7.00 | H2, $\delta_H$ 4.04<br>H3, $\delta_H$ 2.10<br>H4, $\delta_H$ 0.84<br>H5, $\delta_H$ 0.82 |
| -S-CH=CH-NH- | -NH, $\delta_H$ 7.16 | H1, $\delta_H$ 5.53 ($J$ = 7.3 Hz)<br>H2, $\delta_H$ 7.14 ($J$ = 7.3 Hz) |

**Supplementary Table 7.** Statistics of X-ray crystallographic data collection and model refinements of $TvaF_{S-87}$ and CypD in their apo-forms.

| **Data set** | SeMet-TvaF | SeMet-CypD | CypD |
|---|---|---|---|
| **Data collection** | | | |
| Wavelength (Å) | 0.97915 | 0.97890 | 0.97891 |
| Space group | *I*23 | *I*4 | *I*4 |
| Cell dimensions | | | |
| a, b, c (Å) | 193.96, 193.96, 193.96 | 140.11, 140.11, 197.53 | 139.62, 139.62, 197.21 |
| α, β, γ (°) | 90, 90, 90 | 90, 90, 90 | 90, 90, 90 |
| Resolution range (Å) | 96.98 - 2.27 (2.33 - 2.27) | 50.00 - 2.38 (2.42 - 2.38) | 50.00 - 2.40 (2.44 - 2.40) |
| $R_{merge}$ (%) [a] | 11.5 (79.3) | 9.1 (124.0) | 21.0 (81.3) |
| $I / \sigma I$ | 29.60 (7.30) | 30.20 (2.00) | 11.00 (2.44) |
| Completeness (%) | 100.00 (100.00) | 100.00 (92.60) | 99.60 (100.00) |
| Redundancy | 20.0 (21.7) | 13.8 (14.3) | 12.6 (13.3) |
| **Refinement** | | | |
| Resolution (Å) | 19.59 - 2.27 (2.35 - 2.27) | 44.31 - 2.38 (2.46 - 2.38) | 34.88 - 2.40 (2.49 - 2.40) |
| No. reflections | 55718 | 75830 | 72525 |
| $R_{work}$ / $R_{free}$ (%) [b] | 17.82 (21.83) /20.37 (25.30) | 18.03 (23.12) /20.14 (21.85) | 18.07 (20.25) /20.19 (26.80) |
| No. of atoms | 5694 | 8890 | 8917 |
| Protein | 5347 | 8203 | 8172 |
| Ligand | 124 | 318 | 364 |
| Water | 223 | 369 | 381 |
| Average B-factor (Å$^2$) | 32.06 | 40.02 | 39.38 |
| R.m.s.deviations | | | |
| Bond lengths (Å) | 0.009 | 0.010 | 0.009 |
| Bond angles (°) | 1.230 | 1.370 | 1.260 |
| Ramachandran plot [c] | | | |
| Favored region (%) | 97.45 | 98.44 | 97.48 |
| Allowed region (%) | 2.55 | 1.56 | 2.52 |
| Outliers (%) | 0 | 0 | 0 |

[a] $R_{merge} = \sum|I_i - I_m|/\sum I_i$, where $I_i$ is the intensity of the measured reflection and $I_m$ is the mean intensity of all symmetry related reflections.

[b] $R_{work} = \Sigma||F_{obs}| - |F_{calc}||/\Sigma|F_{obs}|$, where $F_{obs}$ and $F_{calc}$ are observed and calculated structure factors.

$R_{free} = \Sigma_T||F_{obs}| - |F_{calc}||/\Sigma_T|F_{obs}|$, where T is a test data set of about 5% of the total reflections randomly chosen and set aside prior to refinement.

[c] Defined by Molprobity.

Numbers in parentheses represent the value for the highest resolution shell.

**Supplementary Table 8.** Statistics of X-ray crystallographic data collection and model refinement of CypD in complex with the **ISLVS** peptide

| **Data set** | CypD/ISLVS peptide complex |
|---|---|
| **Data collection** | |
| Wavelength (Å) | 0.97891 |
| Space group | *F*23 |
| Cell dimensions | |
| a, b, c (Å) | 228.74，228.74，228.74 |
| α, β, γ (º) | 90, 90, 90 |
| Resolution range (Å) | 50.00 - 2.30 (2.34 - 2.30) |
| $R_{merge}$ (%) [a] | 24.5 (56.4) |
| $I$ / σ$I$ | 18.45 (8.14) |
| Completeness (%) | 99.30 (99.30) |
| Redundancy | 37.4 (34.6) |
| **Refinement** | |
| Resolution (Å) | 44.02 - 2.30 (2.38 - 2.30) |
| No. reflections | 44029 |
| $R_{work}$ / $R_{free}$ (%) [b] | 20.34 (20.06) /22.25 (23.18) |
| No. of atoms | 3106 |
| Protein | 2820 |
| Ligand | 106 |
| Water | 180 |
| Average B-factor (Å$^2$) | 31.91 |
| R.m.s.deviations | |
| Bond lengths (Å) | 0.009 |
| Bond angles (°) | 1.26 |
| Ramachandran plot [c] | |
| Favored region (%) | 98.29 |
| Allowed region (%) | 1.71 |
| Outliers (%) | 0 |

[a] $R_{merge} = \sum|I_i - I_m|/\sum I_i$, where $I_i$ is the intensity of the measured reflection and $I_m$ is the mean intensity of all symmetry related reflections.

[b] $R_{work} = \Sigma||F_{obs}| - |F_{calc}||/\Sigma|F_{obs}|$, where $F_{obs}$ and $F_{calc}$ are observed and calculated structure factors.

$R_{free} = \Sigma_T||F_{obs}| - |F_{calc}||/\Sigma_T|F_{obs}|$, where T is a test data set of about 5% of the total reflections randomly chosen and set aside prior to refinement.

[c] Defined by Molprobity.

Numbers in parentheses represent the value for the highest resolution shell.
